# Plant Genotype Influences Physicochemical Properties of Substrate as well as Bacterial and Fungal Assemblages in the Rhizosphere of Balsam Poplar

**DOI:** 10.1101/2020.06.19.161398

**Authors:** Karelle Rheault, Denis Lachance, Marie-Josée Morency, Évelyne Thiffault, Marie Guittonny, Nathalie Isabel, Christine Martineau, Armand Séguin

**Affiliations:** Natural Resources Canada, Canadian Forest Service, Laurentian Forestry Centre, Quebec City, QC, Canada; Research Centre on Renewable Materials, Department of Wood and Forest Sciences, Université Laval, Quebec City, QC, Canada; Research Institute of Mining and Environment (RIME), Université du Québec en Abitibi-Témiscamingue, Rouyn-Noranda, QC, Canada

**Keywords:** *Populus balsamifera*, Genotype-by-environment interactions, Microbiome, Tree genetics, Mine waste revegetation, Plant-microbe interactions

## Abstract

Abandoned unrestored mines are an important environmental issue since they typically remain unvegetated for decades, exposing vast amounts of mine waste to erosion. Several factors limit the revegetation of these sites, including extreme abiotic conditions and unfavorable biotic conditions. However, some pioneer tree species having high level of genetic diversity, such as balsam poplar (*Populus balsamifera*), are able to naturally colonize these sites and initiate plant succession. This suggests that some tree genotypes are likely more suited for acclimation to the conditions of mine wastes. In this study, two contrasting mine waste storage facilities (waste rock versus tailings) from the Abitibi region of Quebec (Canada), on which poplars have grown naturally, were selected. First, we assessed *in situ* the impact of vegetation presence on each type of mine wastes. The presence of balsam poplars improved soil health locally by improving physicochemical properties (e.g. higher nutrient content and pH) of the mine wastes and causing an important shift in their bacterial and fungal community compositions, going from lithotrophic communities that dominate mine waste environments to heterotrophic communities involved in nutrient cycling. Next, in a greenhouse experiment, ten genotypes of *P. balsamifera* collected on both mine sites and from a natural forest nearby were grown in these mine wastes. Tree growth was monitored during two growing seasons, after which the effect of genotype-by-environment interactions was assessed by measuring the physicochemical properties of the substrates and the changes in microbial communities, using a metabarcoding approach. Although substrate type was identified as the main driver of rhizosphere microbiome diversity and community structure, a significant effect of tree genotype was also detected, particularly for bacterial communities. Plant genotype also influenced aboveground tree growth and the physicochemical properties of the substrates. These results highlight the influence of balsam poplar genotype on the soil environment and the potential importance of tree genotype selection in the context of mine waste revegetation.

## 1 Introduction

Abandoned and unrestored mine sites represent an important environmental issue since they typically remain unvegetated for decades. They often constitute a major eyesore to adjacent communities and pose further risk to surrounding ecosystems as vast amounts of soil, waste rock and tailings are exposed to aeolian and water erosion (Mendez and Maier, 2008). Soil microorganisms are in part responsible for the negative impacts of mine sites, mainly through the formation of acid mine drainage, mostly at pH lower than five (Bussière *et al.*, 2005). In addition, the nature and composition of mine wastes make them challenging substrates for plant growth: they are nutrient poor, have either a very low or very high pH and have poor physical structure and deficient water holding capacity (Kumaresan *et al.*, 2017).

Revegetation could help initiate the restoration of these ecosystems. Established plants, together with their associated microbiome, have the ability to reduce acid mine drainage and modify the physicochemical properties of their environment, improving soil quality and fertility (Hubbard *et al.*, 2018). Furthermore, plants form a biological cap that reduces soil erosion while increasing water retention and organic matter content in coarse mine wastes, which contributes to soil stability (Tordoff *et al.*, 2000).

Parameters used to assess the success of revegetation are based on plant, soil and microbial criteria. Plant criteria include plant survival and biomass, leaf and shoot metal concentrations, establishment of other native colonizers and ability to self-propagate (Tordoff *et al.*, 2000). Soil criteria include improvement in soil structure such as increased soil aggregation and reduced erosion; and improvement in soil physicochemical properties such as less acidic pH, increased organic matter content and increased metal bioavailability and mobility (Mendez and Maier, 2008; Lauber *et al.*, 2009). Although currently not widely used, microbial criteria include a decrease in autotrophic bacteria followed by an increase in heterotrophic bacteria and fungi (Rosario *et al.*, 2007) and an increase in bacterial diversity and richness (Garbeva *et al.*, 2004).

The efficiency of revegetation is largely dependent on the establishment of a large root network and beneficial root-soil microbe interactions (Callender *et al.*, 2016). Poplars with their high level of genetic diversity could thus be ideal candidates for revegetation purposes. Indeed, poplars are pioneer trees: they rapidly grow in a vast range of environmental conditions and they easily propagate by root suckering and crown breakage (Dickmann and Kuzovkina, 2014). They also have a large and deep root system (Braatne *et al.*, 1996). As a perennial species, they can tolerate harsh environments and promote the establishment of primary and successive plant species through the addition of soil nutrients from their abundant litter production, increasing ecosystem health and function (Pardon *et al.*, 2017). They can establish root associations with both arbuscular mycorrhizal and ectomycorrhizal fungi (Gehring *et al.*, 2006), as well as other endophytic and rhizospheric organisms, thereby increasing access to nutrients, relieving abiotic stresses such as hydric stress, suppressing plant pathogens, altering phenology and promoting plant growth (Bulgarelli *et al.*, 2013). Finally, *Populus* is considered as a model genus for the study of woody perennials and therefore massive genomic resources are available (e.g. fully sequenced genome, Tuskan *et al.*, 2006) which provides unique opportunity to support selection of most suited genotypes to address ecosystem restoration issues (Fini *et al.*, 2017).

Plant species produce specific root exudates that will attract specific microorganisms, resulting in dissimilar selective pressures on microbial communities (Philippot *et al.*, 2013). In addition, differences between plant genotypes can have a significant impact on the microbiomes of their rhizosphere (Badri *et al.*, 2009; Lebeis *et al.*, 2015; Veach *et al.*, 2019). This suggests that plant genotype could significantly affect the recruitment of specific microorganisms, thus leading to further differences in plant growth and ecosystem functioning. Many studies have attempted to characterize the root microbiome of *Populus*. It has been shown that soil type is the main driver of microbial community assembly, since physicochemical properties, such as granulometry, pH and nutrient content, influence microbial composition and functional group prevalence (Gottel *et al.*, 2011; Danielsen *et al.*, 2012; Shakya *et al.*, 2013; Cregger *et al.*, 2018). However, genetic variations of the host plant are also associated with differential microbial colonization (Bonito *et al.*, 2014). Differentiating between the effects of soil properties and those of the host plant genotype has not yet been sufficiently addressed (Bonito *et al.*, 2019).

The aim of this study was to assess the suitability of *P. balsamifera* for the revegetation of abandoned mine sites and to determine the impact of genotype-by-environment interactions on the improvement of soil health and the rhizosphere microbiome. To do so, two contrasting mine sites from Abitibi, Quebec, where poplars have naturally grown, were selected. First, in a field study, the impact of balsam poplar presence on the waste rock of the first site and on tailings of the second site, was assessed, regarding their physicochemical properties and the composition of their microbiome using a metabarcoding approach. Then, in a greenhouse experiment, ten genotypes of *P. balsamifera* collected from both mine sites and from a natural forest nearby were grown in mine wastes (waste rock and tailings) and evaluated. Tree growth was monitored over two growing seasons, after which the effect of genotype-by-environment interactions on microbial community dynamics were investigated using the same genomic tools as for the field study. The main objectives were (1) to assess the effect of vegetation presence on two contrasting mine wastes by measuring physicochemical properties and characterizing bacterial and fungal communities of the mine wastes *in situ*; (2) to assess the effect of balsam poplar genotype on the physicochemical properties of their substrate and the diversity and composition of their rhizosphere microbiome; and (3) to assess the effect of genotype-by-environment interactions on the rhizosphere microbiome in contrasting substrates.

## 2 Material and Methods

### 2.1 Field site description and sampling methods

Two mine sites, 50 km apart, located in Abitibi (western Quebec, Canada) were chosen for the contrasting characteristics of their mine wastes and for the fact that they both have balsam poplars naturally growing on the periphery, among other plant species such as willows (*Salix*), trembling aspen (*P. tremuloides)*, alder (*Alnus*) and birch (*Betula*). The Westwood site, formerly the Doyon site, is characterized by its acid generating and coarse waste rock piles; the La Corne Mine site is dominated by neutral and fine-grained tailing piles. Both mine wastes are nutrient poor. The Westwood site is a former gold mine, recently put back into operation and owned by IAMGOLD Corporation. The La Corne Mine site is a former molybdenum and bismuth mine out of operation since 1972 and owned by Romios. See Figures S1 and S2 for pictures of the sites and Figure S3 for a summary of the field and the greenhouse experiments. Mine wastes, vegetated soil samples and tree cuttings were sampled in November 2016.

#### 2.1.1 Soil sampling on site

Approximately 150 L of mine waste was collected from the tailing stockpiles at the La Corne Mine site as well as from waste rock stockpiles from the former Doyon mine site at the Westwood site. Samples were collected from the top 20 cm using a shovel and were stored in 25 L plastic boxes at 4°C until used. Five subsamples of 15 g from both mine wastes were also taken for DNA extraction and physicochemical analyses. In addition, from each site, five samples of 15 g of bulk soil were collected from areas colonized by balsam poplars to assess the impact of vegetation on mine substrates in a natural setting. Upon arrival to the laboratory, subsamples were taken from each 15 g samples, placed in 1.5 mL tubes and stored at −20°C until DNA extraction. The remainder of each sample was air-dried for physicochemical analyses.

#### 2.1.2 Cuttings sampling and genotyping

Eight mature balsam poplars per mine site and two from a natural forest near the La Corne Mine site were selected, from which branches were harvested to produce cuttings. Branches were dormant when collected. Harvested branches were kept on ice during transport and stored frozen at −5°C immediately upon arrival to the laboratory, until used. In order to reduce the risk of harvesting the same genotype (clone), trees were sampled at a minimum distance of approximately 200 m. Their unique genotypes were then verified using a 40 SNP-array designed to reveal *P. balsamifera* intraspecific variations (Table S1), as described by Meirmans *et al.* (2017). For this, DNA was extracted from bud tissue using a Nucleospin 96 Plant II kit (Macherey-Nagel, Bethlehem, PA, USA) following the manufacturer’s protocol for centrifugation processing with the following modification: buffer PL2 was used at the cell lysis step and was heated for 2 h at 65°C instead of 30 min. All samples were sent to the Genome Quebec Innovation Centre at McGill University to be genotyped.

### 2.2 Cuttings selection and growth

From November 2016 to January 2017, cuttings were first started using a hydroponic system (see below) to better allow for root development, then transferred into pots and grown until the end of April 2017. Trees were at least 30 cm tall at the start of the experiment. Genotypes for which a minimum of nine replicates (three replicates per substrate type) remained at the end of the tree production process were kept for the experiment, leaving four genotypes per mine site and two genotypes from the natural forest.

Ten cuttings, each containing three buds, were prepared from the tree branches collected from each tree in the field. The base of each was cut diagonally, dipped in a rooting powder (STIM-ROOT No3, Plant Product Co. LTD) and inserted into a rooting medium (ROOTCUBES® 1½’ square, Smithers-OASIS) in the hydroponic system. Buds containing flowers were removed. The cuttings were watered automatically twice a day, at 08:00 and 20:00, for 4 minutes (just enough time for the container to be filled and drained slowly). Day/night greenhouse temperatures were set at 22/18°C with 16 hours supplemented lighting (less than 250 W/m^2^) between 08:00 and 24:00. Every four weeks of growth in hydroponic, a rooting fertilizer (8 mL / 40 L; Roots&Rhizo, Fred T. Lizer) was added to the irrigation system to help root development.

After two months of growth in the hydroponic system, cuttings were transferred to pots. The potting mix consisted of four parts peat (Agro Mix G6, Fafard), two parts vermiculite (Perlite Canada Inc.) and one part Turface (calcined clay particles, Turface MVP, Turface Athletics®). The potting mix was watered and autoclaved to kill insects potentially present in the peat. Nine grams of slow release fertilizer 18.6.8 (Nutricote Total, Type:100, Chisso-Asahi Fertilizer Co. LTD) was then added per liter of potting mix. Cuttings were planted in 250 mL square pots. Temperature and light conditions were the same as for the hydroponic growth. For the first two months, the trees were watered automatically by a drip irrigation system twice a day for 3 minutes at 08:00 and 20:00. For the last month, the watering program was changed to three times per day for 5, 3 and 5 minutes at 08:00, 16:00 and 24:00. Trees were given extra manual watering as required.

### 2.3 Greenhouse experiment

#### 2.3.1 Soil sieving and mixing

Mine wastes were sieved through a 10 mm sieve before use. A peat mix composed of four parts peat, two parts vermiculite and one part Turface was also prepared. Using a cement mixer, equal volumes of mine wastes and the peat mix were combined. Thus, three treatments were obtained: (1) the mix containing waste rock from the Westwood site and the peat mix (waste rock; WR); (2) the mix containing tailings from the La Corne Mine site and the peat mix (tailing; TA); and (3) a control substrate composed of peat, vermiculite, Turface and slow release fertilizer as described in section 2.2 (control; CO).

#### 2.3.2 Experimental design

At the end of April, three trees from each genotype were repotted in 4 L pots with each mixture treatment and randomly distributed in a greenhouse. The trees were again watered automatically using the drip irrigation system three times per day for 5 minutes at 08:00, 16:00 and 24:00. Temperature and light settings were as described previously.

The trees were transferred outside in August after they formed buds and started to lose leaves so as to harden off naturally. Plants were watered manually as required. The trees were returned to the greenhouse in January set to a day/night temperature of 10/5°C with 10 hours of supplemented lighting from 08:00 to 16:00 hours. The trees were immediately cut back to between 30 cm and 50 cm high so as to keep 10 buds per tree, including both lateral and terminal buds. Temperature and lighting were gradually raised by 5°C and two hours respectively at two weeks interval until reaching maximum day/night temperatures of 22/18°C and 14 hours lighting (between 06:00 and 20:00). Irrigation started one week after the first buds started to flush, again using the drip irrigation system. Trees were initially watered for 3 minutes at 08:00 once every three days. After ten days, this was increased to once every two days, then to once a day a week later, and finally to twice a day, for 5 minutes, at 08:00 and 20:00 11 days after that. The trees were grown for about three months, until they naturally set bud.

#### 2.3.3 Tree growth and health measurements

The following parameters were measured to assess tree growth and health during the two growing seasons: variation in height (growth); chlorophyll content of leaves; shoot diameter; and dry biomass of shoots and leaves. Height was measured from the soil surface to the terminal bud at the beginning and end of the first season. Height was not measured during the second season because trees were cut back to 40-50 cm at the start of the season. Growth was expressed as a percentage of the difference in heights between the beginning of the experiment and the end of the first season of growth:

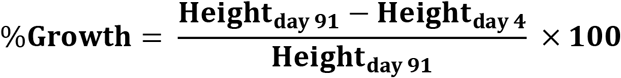

As an estimation of leaf nitrogen content, the leaf chlorophyll content, or “greenness”, was measured with a spectrophotometer, following the manufacturer’s instructions (SPAD 502 Plus Chlorophyll Meter, Spectrum Technologies, Inc.). Chlorophyll measurements were made around the middle of each growing season, starting on the fifth leaf from the base of the plant and on every third leaf thereafter, avoiding diseased or immature leaves that were not representative of the whole tree. Shoot diameter was measured 20 cm above the initial cutting at the end of the experiment. Dry biomass was the total mass of the shoot and leaves from the second growth season after drying at 50°C for 7 days. Appearance of leaves was noted once a week during the two growing seasons.

#### 2.3.4 Rhizosphere and bulk sampling

Trees were removed from their pots and the shallow roots removed. Fine roots less than 2 mm in diameter were then sampled, avoiding the taproot in the center of the pots. Bulk soil was obtained by collecting the soil that detached from those roots with gentle shaking. Rhizosphere soil, the soil still attached to the roots after shaking, was then collected by placing the roots into 50 mL Falcon tubes containing 25 mL of sterile PBS (8 g NaCl, 0.2 g KCl, 1.44 g Na_2_HPO_4_, 0.24 g KH_2_PO_4_, pH 7.4, 1 L distilled water). After briefly shaking the tubes, the roots were removed, the tubes centrifuged at 4700 RPM for 10 minutes at 4°C, and the supernatant discarded. The pellet or rhizosphere soil, was then collected and placed on sterile filter papers to absorb excess moisture, then stored in 1.5 mL tubes at −20°C until DNA extraction.

### 2.4 Processing of samples

#### 2.4.1 Physicochemical analyses of bulk soil

Samples of bulk soil, either collected directly on the mine sites or from the greenhouse experiment, were air-dried, sieved at 2 mm, and kept in plastic bags until further processing. One gram of soil was ground to a fine powder of 0.5 mm prior to physicochemical analyses for carbon, nitrogen and sulfur.

Carbon (C), nitrogen (N) and sulfur (S) were quantified using the TruMac® CNS analyzer (LECO Corporation, MI, USA) following the manufacturer’s protocol. Water pH and buffer pH were measured using the methods described by the Canadian Society of Soil Science (Gregorich and Carter, 2007) using the Thermo Scientific™ Orion™ 2-Star Benchtop pH meter. Extractable phosphorus (P) and exchangeable cations (potassium (K), calcium (Ca), magnesium (Mg), manganese (Mn), iron (Fe), aluminum (Al), and sodium (Na)) were extracted with a Mehlich III extraction buffer (Gregorich and Carter, 2007) and analyzed by inductively coupled plasma (ICP) using an optical emission spectrometer (OES) (Optima 7300 DV, Perkin Elmer, Waltham, MA).

#### 2.4.2 DNA isolation and library preparation

Bulk soil samples from the field experiment and rhizosphere soil samples from the greenhouse experiment were kept for microbiome analyses. Up to 250 mg of these samples were transferred to PowerBead tubes for DNA extraction using the DNeasy PowerSoil DNA Isolation Kit (Qiagen, Valencia, CA, USA), in accordance with the manufacturer’s instructions, except that DNA was eluted in 50 μL instead of 100 μL. DNA was quantified using a Qubit dsDNA HS Assay Kit and Qubit Fluorometer (Thermo Fisher Scientific, Waltham, MA, USA). Library preparation for Illumina sequencing was performed according to the manufacturer’s instructions for user-defined primers (Illumina, 2013)^1^, with the following modifications. Each sample was amplified in triplicate to ensure reproducibility (Schmidt *et al.*, 2013; Kennedy *et al.*, 2014). Bacterial communities were amplified using primers 515F-Y (5’-GTGYCAGCMGCCGCGGTAA-3’) and 926R (5’-CCGYCAATTYMTTTRAGTTT-3’) targeting the V4-V5 regions of the 16S rRNA gene of bacteria and archaea (Parada *et al.*, 2016; Rivers, 2016). The ITS2 region of the fungal ribosomal DNA was amplified using the primer set ITS9F (5’-GAACGCAGCRAAIIGYGA-3’) and ITS4R (5’-TCCTCCGCTTATTGATATGC-3’) (White *et al.*, 1990; Rivers, 2016). Primers contained the required Illumina adaptors at the 5′ end of the primer sequences (5′-TCGTCGGCAGCGTCAGATGTGTATAAGAGACAG-3′ for the forward primer and 5′-GTCTCGTGGGCTCGGAGATGTGTATAAGAGACAG-3′ for the reverse primer). PCR reactions were set up by first mixing 37.5 μl of HotStarTaq Plus Master Mix (QIAGEN Inc., Germantown, MD, USA), 27 μL RNase-free water, 1.5 μL of each 10 μM primer and 7.5 μL of gDNA at 5 ng/μL. The final volume of 75 μL was then equally distributed in three 96-well plates placed in distinct thermocyclers. Thermal cycling conditions were as follows: initial denaturation at 95°C for 5 minutes; 40 cycles (for ITS2 amplification; and 35 cycles for 16S amplification) at 94°C for 30 s, 50°C for 30 s, 72°C for 1 minute; and a final elongation at 72°C for 10 minutes. PCR products were pooled and purified using 81 μL of magnetic beads solution (Agencourt AMPure XP), then unique codes were added to each sample using the Nextera XT Index Kit, in accordance with the above-mentioned Illumina’s protocol. Indexed amplicons were purified with magnetic beads, quantified using a Qubit dsDNA BR Assay Kit (Thermo Fisher Scientific, Waltham, MA, USA) and combined at equimolar concentration. Paired-end sequencing (2 × 250 bp) of the pools was carried out on an Illumina MiSeq at the Illumina Sequencing Platform, Nucleic Acids Solutions, Aquatic and Crop Resource Development, National Research Council Canada-Saskatoon. To compensate for low base diversity when sequencing amplicon libraries, PhiX Control v3 Library was denatured and diluted to 12.5 pM before being added to the denatured and diluted amplicon library at 15% v/v. The amplicon libraries were sequenced at a concentration of 6.5 pM for most of the sequencing runs. The Illumina data generated in this study was deposited in the NCBI Sequence Read Archive and is available under the project number PRJNA615167.

### 2.5 Bioinformatic analyses

#### 2.5.1 Sequences assignation

All bioinformatics analyses were performed in QIIME (version 1.9.1) (Caporaso *et al.*, 2010). Briefly, sequence reads were merged with their overlapping paired-end (fastq_mergepairs), trimmed to remove primers (fastx_truncate), and filtered for quality (fastq_filter) using USEARCH (Edgar, 2010). Unique identifiers were inserted into the header of the remaining high-quality sequences, and sequences from the different samples were pooled together (add_qiime_labels) prior to further analyses.

UPARSE (Edgar, 2013) was then used to dereplicate the sequences (derep_fulllength), discard singletons (sortbysize), group high quality reads into operational taxonomic units (OTUs) using a 97% identity threshold (cluster_otus) (Schloss *et al.*, 2009), and identify chimeras (uchime_ref). The taxonomic assignment of OTUs was done using the QIIME “assign_taxonomy” command with Mothur as the assignment method and Greengenes Database (McDonald *et al.*, 2012) files as the reference for bacteria and the UNITE database (Abarenkov *et al.*, 2010) files for fungi. The “make_OTU_table’’ command was then used to generate the OTU table in the “biom” format, which was then used by QIIME in the next steps. OTUs from nonbacterial (or nonfungal) taxa were excluded using the “filter_taxa_from_otu_table’’ command. For the 16S rRNA analysis, sequences corresponding to chloroplasts, mitochondria and to the kingdom *Plantae* were removed; for the ITS2 analysis, sequences assigned to the kingdom *Protozoa*, *Protista*, *Chromista* and *Plantae* were removed.

OTUs with relative abundances below 0.005% were excluded as previously described (Bokulich *et al.*, 2013) prior to diversity analyses using the “filter_otus_from_otu_table” command. Each sample was rarefied to the lowest number of reads observed among libraries from each data set with QIIME’s “single_rarefaction” command, so the rarefied samples all contained the same number of sequences. Finally, the OTU rarefied list file was used in QIIME’s “core_diversity_analyses” and “alpha_diversity” commands to generate alpha diversity measures (chao1 and Shannon indices), calculate the beta-diversity between samples (Bray-Curtis dissimilarity), and generate community composition profiles at different taxonomic levels.

Fungal functional groups were predicted using a homemade Python script based on a predetermined list of fungi genus associated with their respective function (Tedersoo *et al.*, 2014). First, the script optimizes the list to detect and remove non-unique entries (these data are compared to each other to determine which data contains the most information). The corrected data is then compared to an excel file, where the genus is the search key: the script analyzes each line of the file to extract the genus of each OTU; this genus is then compared to a reference library to identify the associated biological function. OTUs unidentified to the genus level are assigned unidentified function.

#### 2.5.2 Statistical analyses

Statistical analyses were conducted in R version 3.5.3 (Rproject.org) and figures were produced using the package “ggplot2” package. Statistical significance was determined at *p* < 0.05 throughout the analyses. Parametric assumptions were verified before analysis: data normality was checked graphically with normal quantile–quantile plots and computationally with the Shapiro–Wilk test of normality using the “shapiro.test” function. Homoscedasticity was verified using both the Bartlett test (“bartlett.test” function) and the Fligner-Killeen test (“fligner.test” function). Data were transformed using square root (“sqrt” function) or Tukey’s Ladder of Powers (“transformTukey” function, “rcompanion” package) when necessary to meet parametric ANOVA assumptions. A generalized least squares model (“nlme” package) with a stepwise selection and Akaike’s Information Criterion (AIC) minimization approach was performed with vegetation presence (vegetated or unvegetated soil) and waste type (waste rock or tailings) as explanatory variables for the field experiment and tree genotype and substrate type (WR, TA or CO) as explanatory variables for the greenhouse experiment. Genotype origin (La Corne Mine site, Westwood site or natural forest) was also included in these models (“corComSymm” correlation) to account for its overall large influence. Substrate types were weighted (“varIdent”, weights) to reduce variance due to the fact that they were highly different in their physicochemical properties.

Two-way ANOVAs (“anova” function) were used to discern how waste type, vegetation presence, substrate type, genotype, and their interactions influenced taxa relative abundances, physicochemical properties of substrates, and alpha diversity indices. When a factor was revealed as a statistically significant predictor, a Tukey HSD post-hoc pairwise comparison test (“predictmeans” function, adjusted to “tukey”, “predictmeans” package) was performed between all treatments. For the greenhouse experiment, if the interaction between substrate type and genotype was deemed statistically significant, additional analyses were performed for each substrate type separately to better assess the effect of genotype.

Spearman linear correlation analyses were performed using the “corr.test” function (“psych” package) to determine if there were correlations between physicochemical properties of the substrates, tree growth and taxa relative abundances for both field and greenhouse experiments.

Non-Euclidean distances were calculated from Bray Curtis dissimilarity matrices and implemented in a non-metric multidimensional scaling plot (NMDS, “metaMDS” function, “vegan” package) to visualize both bacterial and fungal community compositional differences between vegetation presence and waste type on the field, and balsam poplar genotypes and substrate type in the greenhouse experiment. A permutational multivariate analysis of variance model (PERMANOVA, “adonis2” function, “vegan” package; (Oksanen *et al.*, 2019)) was also implemented to discern the amount of variation attributed to each factor and their interaction (with 999 permutations). Additional multivariate analyses were performed on communities from each substrate type of the greenhouse experiment separately to better assess the effect of genotype. Variance heterogeneity between the a priori selected groups was tested with the functions “betadisper” and “permutest”, in the “vegan” package. Clusters between treatments were determined by a multilevel pairwise comparison test (“pairwise.adonis2” function, “pairwiseAdonis” package). Correlations between NMDS axes and physicochemical properties of substrates were determined using the “envfit” function (with 999 permutations, “vegan” package).

One sample (genotype C21 in control substrate) was removed from analyses because its results were considered aberrant. Two samples (genotype W13 in tailings and waste rock) failed to amplify and/or get sequenced for ITS and were therefore excluded from further analyses.

## 3 Results

### 3.1 Assessment of vegetation’s effect on mine wastes under field conditions

#### 3.1.1 Soil physicochemical properties

Physicochemical analyses of the mine wastes indicated absence of N, low concentration of C and macronutrients such as P and K, and a relatively high concentration of elements like Fe and S in the waste rock from the Westwood site (Table 1). Additionally, the pH of waste rock was highly acidic with values varying between 2.47 and 3.02, whereas in tailings from La Corne Mine site the pH was almost neutral with values between 5.93 and 7.77. For both tailings and waste rock, the vegetated soils contained significantly higher levels of C, N, K, P, Ca, Mg and Mn than the mine wastes (Table 1). Vegetation also reduced S and Fe concentrations and increased pH in waste rock, and reduced pH in tailings.

**Table 1.** Physicochemical properties of substrates from the field. Legend: CEC: Cation exchange capacity; BCSR: Base cation saturation ratio. Significant differences are highlighted in bold.

|  | Reference forest soil | Westwood |  |  | La Corne Mine |  |  |
| --- | --- | --- | --- | --- | --- | --- | --- |
|  |  | Waste rock | Vegetated | p-value | Tailings | Vegetated | p-value |
| **C total (%)** | 6.218 | 0.084 | 0.863 | **0.002** | 0.042 | 2.983 | < **0.001** |
| **N total (%)** | 0.253 | < 0.006 | 0.021 | 0.053 | < 0.006 | 0.106 | < **0.001** |
| **S total (%)** | 0.045 | 0.700 | 0.134 | < **0.001** | 0.017 | 0.030 | 0.091 |
| **pH** | 4.9 | 2.8 | 4.7 | < **0.001** | 6.4 | 4.5 | < **0.001** |
| **P (mg/kg)** | 9.83 | 7.02 | 23.98 | < **0.001** | 0.68 | 3.80 | **0.004** |
| **K (cmol<sub>c</sub>/kg)** | 0.418 | 0.011 | 0.199 | < **0.001** | 0.053 | 0.202 | < **0.001** |
| **Ca (cmol<sub>c</sub>/kg)** | 7.69 | 8.96 | 1.80 | 0.069 | 0.43 | 2.93 | < **0.001** |
| **Mg (cmol<sub>c</sub>/kg)** | 1.590 | 0.564 | 0.676 | 0.413 | 0.272 | 0.902 | < **0.001** |
| **Mn (cmol<sub>c</sub>/kg)** | 0.143 | 0.011 | 0.040 | **0.024** | 0.024 | 0.073 | **0.003** |
| **Fe (cmol<sub>c</sub>/kg)** | 2.00 | 6.26 | 2.68 | < **0.001** | 2.85 | 3.10 | 0.550 |
| **Na (cmol<sub>c</sub>/kg)** | 0.035 | 0.022 | 0.021 | 0.805 | 0.028 | 0.029 | 0.929 |
| **CEC (cmol<sub>c</sub>/kg)** | 13.6 | 17.0 | 8.7 | 0.102 | 5.3 | 9.4 | **0.003** |
| **BCSR** | 71.6 | 46.3 | 31.3 | 0.344 | 14.1 | 43.2 | < **0.001** |

#### 3.1.2 Microbiome analyses

##### 3.1.2.1 Alpha diversity

Factorial analyses of alpha diversity indices (Figure 1) indicated that vegetation presence (vegetated vs unvegetated soil), waste type (tailings vs waste rock) and the interaction between both factors had a consistently significant effect on bacterial richness (chao1 index: *p* < 0.005) and diversity (Shannon index: *p* < 0.001). For fungal richness and diversity, the interaction between factors was not significant (*p* = 0.066 and 0.355). Pairwise comparison of alpha diversity indices indicated that vegetation presence significantly (*p* < 0.05) increased bacterial richness in both waste types and bacterial diversity in waste rock; and reduced fungal richness and diversity in both waste types.

**Figure 1.**
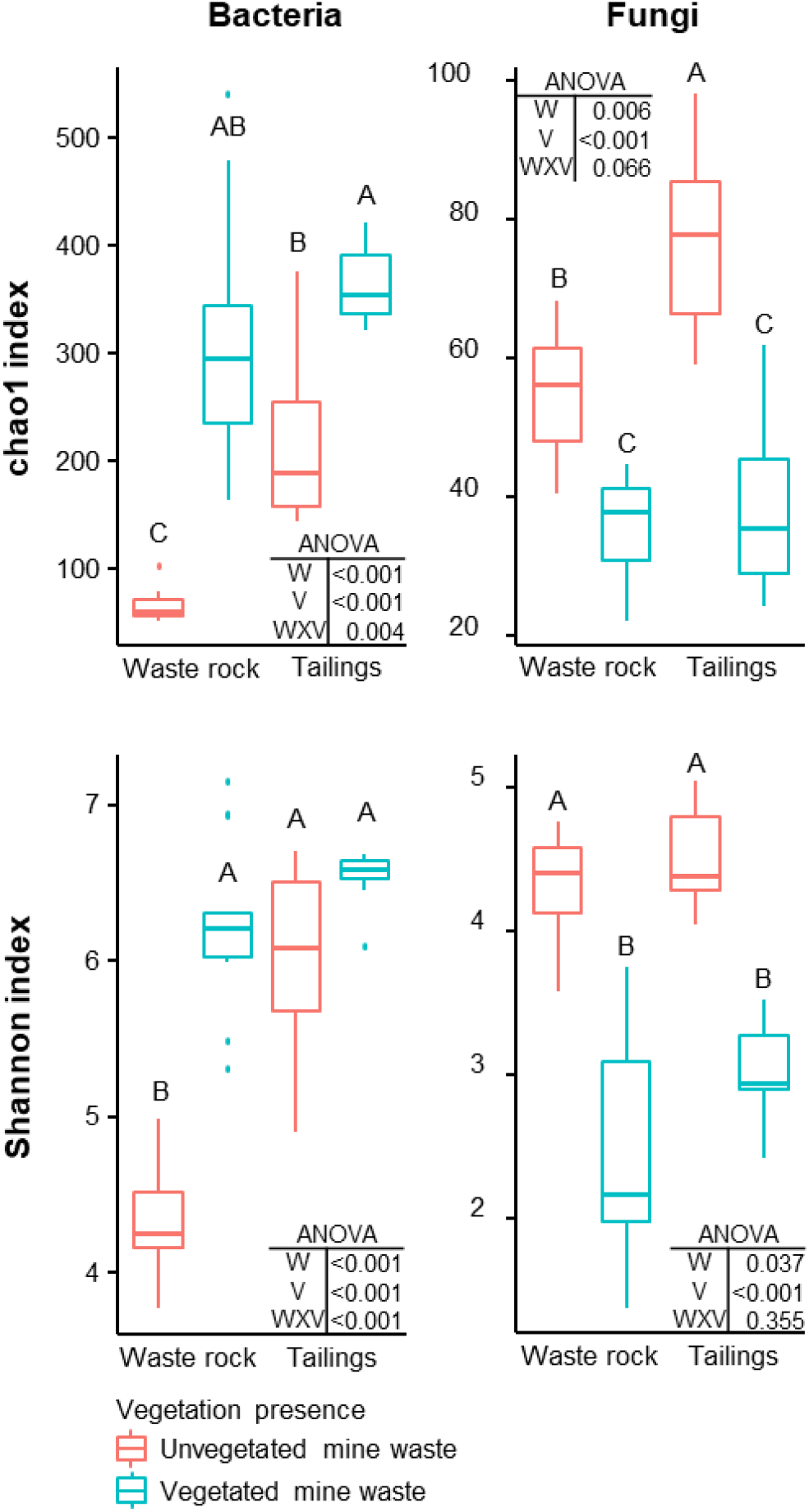
Alpha diversity indices of bacterial and fungal community profiles in field samples. A two-way ANOVA was used to discern how waste type, vegetation presence and their interaction influenced alpha diversity indices. In the ANOVA table, W is waste type; V is vegetation presence; and WXV is the interaction between waste type and vegetation presence. Treatments are composed of two factors: waste rock and tailings as waste type; and vegetated or unvegetated mine waste as vegetation presence. Shared letters between treatments means there is no significant difference between these treatments, as determined by Tukey HSD post-hoc pairwise comparison test (n ≥ 8). Significance level is *p* < 0.05.

##### 3.1.2.2 Beta diversity

Analysis of beta diversity (Figure S4) indicates that bacterial and fungal community structures highly differed between waste types and vegetation presence. The model explains 62% and 35% of the variation in bacterial and fungal community structure, respectively. The main driver of bacterial and fungal community structure was vegetation presence (R^2^ = 31.4% and 18.9%, respectively). Waste type also had a significant effect on community structure (R^2^ = 14.6% and 9.7%). The interaction between these two factors being significant (R^2^ = 14.7% and 6.7%), a multilevel pairwise comparison test was performed on all combinations of waste type by vegetation presence. All combinations clustered separately, indicating that bacterial and fungal community structures differed between each treatment.

##### 3.1.2.3 Taxonomic profiles

Taxonomic profiling of bacterial and fungal communities revealed high heterogeneity as shown by the variability among field replicates. Many bacterial and fungal taxa were only identified at a high taxonomic level. Figure 2 illustrates the relative abundance of the most abundant taxa (>1%, at the genus level) in vegetated soil and mine wastes, at the La Corne Mine site and the Westwood site.

**Figure 2.**
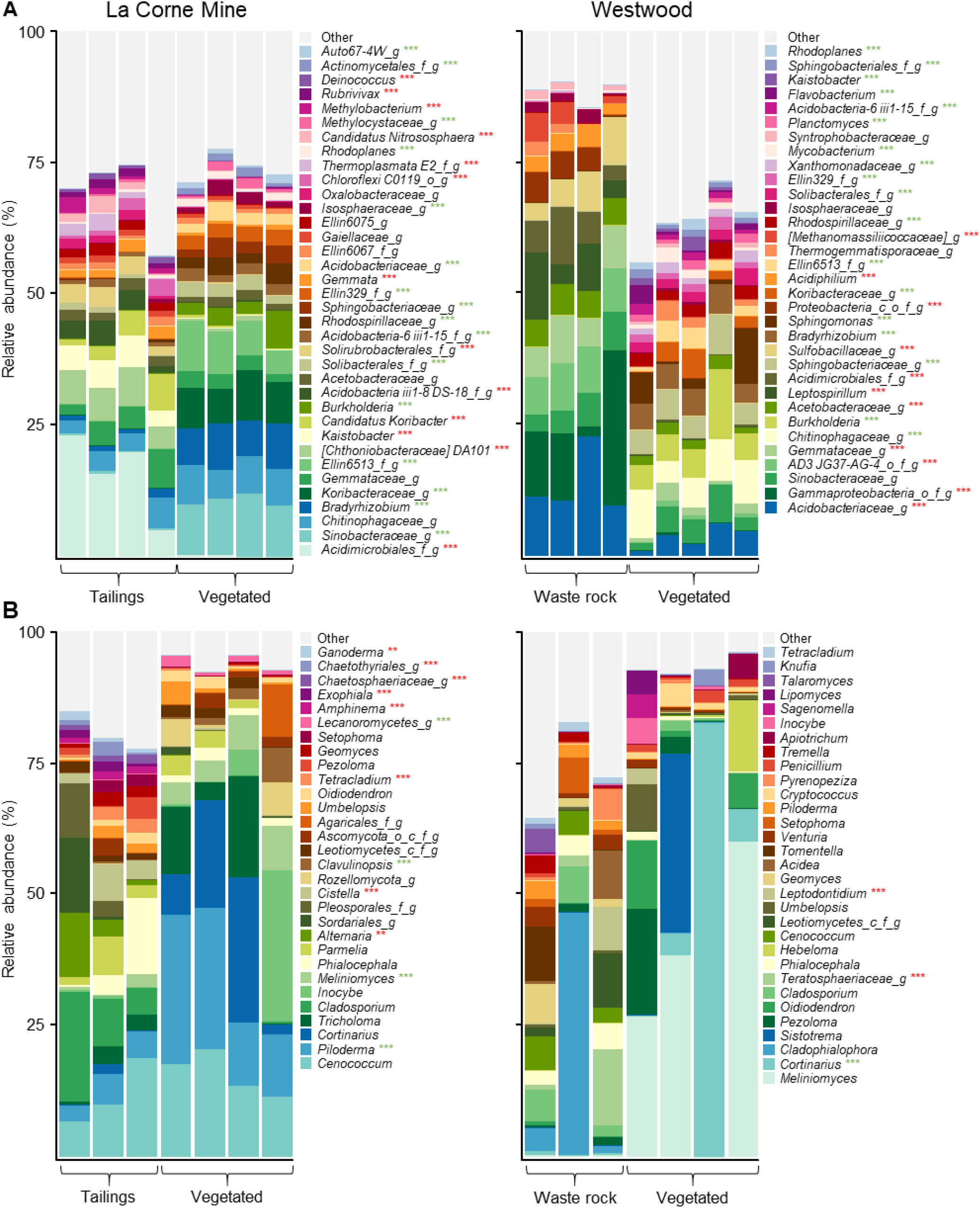
Taxonomic profiles of bacterial and fungal communities in vegetated and unvegetated samples of waste rock from the Westwood site and tailings from the La Corne Mine site. Taxonomic profiles of bacterial communities **(A)** and fungal communities **(B)** at the genus level. Only bacterial and fungal taxa with a relative abundance >1% in at least one treatment are shown. Green and red stars represent a significant increase and decrease, respectively, in the relative abundance of the taxa in the vegetated soil compared to the unvegetated soil as determined by a Tukey HSD post-hoc pairwise comparison test (n ≥ 3). Significance level is represented as follows: *p* < 0.05 *, *p* < 0.01 **, *p* < 0.001 ***.

Factorial analyses of the relative abundances of bacterial taxa (for taxa >1%, Table S2) showed that waste type had a significant effect on 32 taxa, vegetation presence had a significant effect on 44 taxa, and a significant interaction between both factors was detected for 40 taxa. Pairwise comparisons between all treatments revealed that 50 of the 51 most abundant bacterial taxa (>1%, at the genus level) had a significant difference in at least one treatment (*p* < 0.001). In fungal communities (Table S2), waste type had a significant effect on 11 taxa, vegetation presence had a significant effect on 21 taxa and a significant interaction between both factors was detected for 11 taxa. Pairwise comparison between all treatments showed a significant difference in at least one treatment for 21 of the 47 fungal taxa (*p* < 0.01).

The presence of vegetation on mine wastes reduced the relative abundance of microorganisms associated with acid mine drainage like *Acidiphilium*, *Leptospirillum* and *Sulfobacillaceae_g* on waste rock (Harrison Jr, 1981; Hippe, 2000; Hottenstein *et al.*, 2019) (Figure 2). Additionally, it also reduced the relative abundance of fungal plant pathogens like *Alternaria* and *Ganoderma* on tailings and *Teratosphaeriaceae* on waste rock. Conversely, the presence of vegetation increased the relative abundance of many microorganisms previously found in rhizosphere samples and associated with beneficial ecological functions like the ectomycorriza *Meliniomyces* on tailings and the rhizobacteria *Burkholderia* (Caballero-Mellado *et al.*, 2004) and *Rhodoplanes* (Sun *et al.*, 2015) on both mine wastes.

Functions associated with fungal community in all samples comprised ectomycorrhizae (35%), saprotrophs (27%), ericoid mycorrhizae (13%), plant pathogens (3%), white rot (1%) and lichenized (1.8%) and arbuscular (0.05%) mycorrhizae. Factorial analyses of the relative abundances of fungal functions (Table S2) showed that there was no effect of waste type nor interaction between waste type and vegetation presence on functional group prevalence. Pairwise comparisons between all treatments revealed that there was no effect of vegetation on the relative abundance of fungal functions in waste rock. However, the presence of vegetation on tailings significantly increased (*p* < 0.001) the relative abundance of ectomycorrhizae and ericoid mycorrhizae; and significantly decreased the relative abundance of saprotrophs (*p* < 0.001), plant pathogens (*p* < 0.001) and arbuscular mycorrhizae (*p* = 0.003).

All bacterial and fungal taxa were significantly correlated with at least one physicochemical property of the substrates (Table S3). Bacterial genera like *Leptospirillum* and *Acidiphilium*, two bacterial genera associated with the oxidoreduction of iron and acid mine drainage (Harrison Jr, 1981; Hippe, 2000) were strongly correlated with iron content and the reduction of pH. Similarly, *Sulfobacillaceae*, a bacterial family associated with the oxidation of sulfur and acid mine drainage (Hottenstein *et al.*, 2019) was strongly correlated with sulfur content and the reduction of pH.

### 3.2 Assessment of genotype-by-environment effect under greenhouse experiment

#### 3.2.1 Tree growth

Tree growth measurements showed a significant effect of genotype for all parameters, an effect of substrate on three parameters and an interaction between both factors for one parameter only (Figure 3). Tree growth (Fig. 3A), the number of days before buds start to open at the beginning of the second season (Fig. 3B) and chlorophyll content during the first season (Fig. 3C) differed among plant genotypes (*p* < 0.001) but not between substrate types (*p* = 0.155, 0.323 and 0.628 respectively); the interaction between both factors was not significant for these parameters (*p* = 0.526, 0.498 and 0.595). Shoot diameter (Fig. 3E) and biomass produced during the second season (Fig. 3F) differed among plant genotypes (*p* < 0.001) as well as between substrate types (*p* < 0.001 and 0.008 respectively), but again no significant interaction was detected (*p* = 0.937 and 0.361). Shoot diameter and plant biomass measurements were greater in the nutrient-rich control substrate versus mine substrates. Chlorophyll content during the second season (Fig. 3D) differed between genotypes and it was greater in the waste rock for some genotypes (significant interaction between genotype and substrate type, *p* < 0.001).

**Figure 3.**
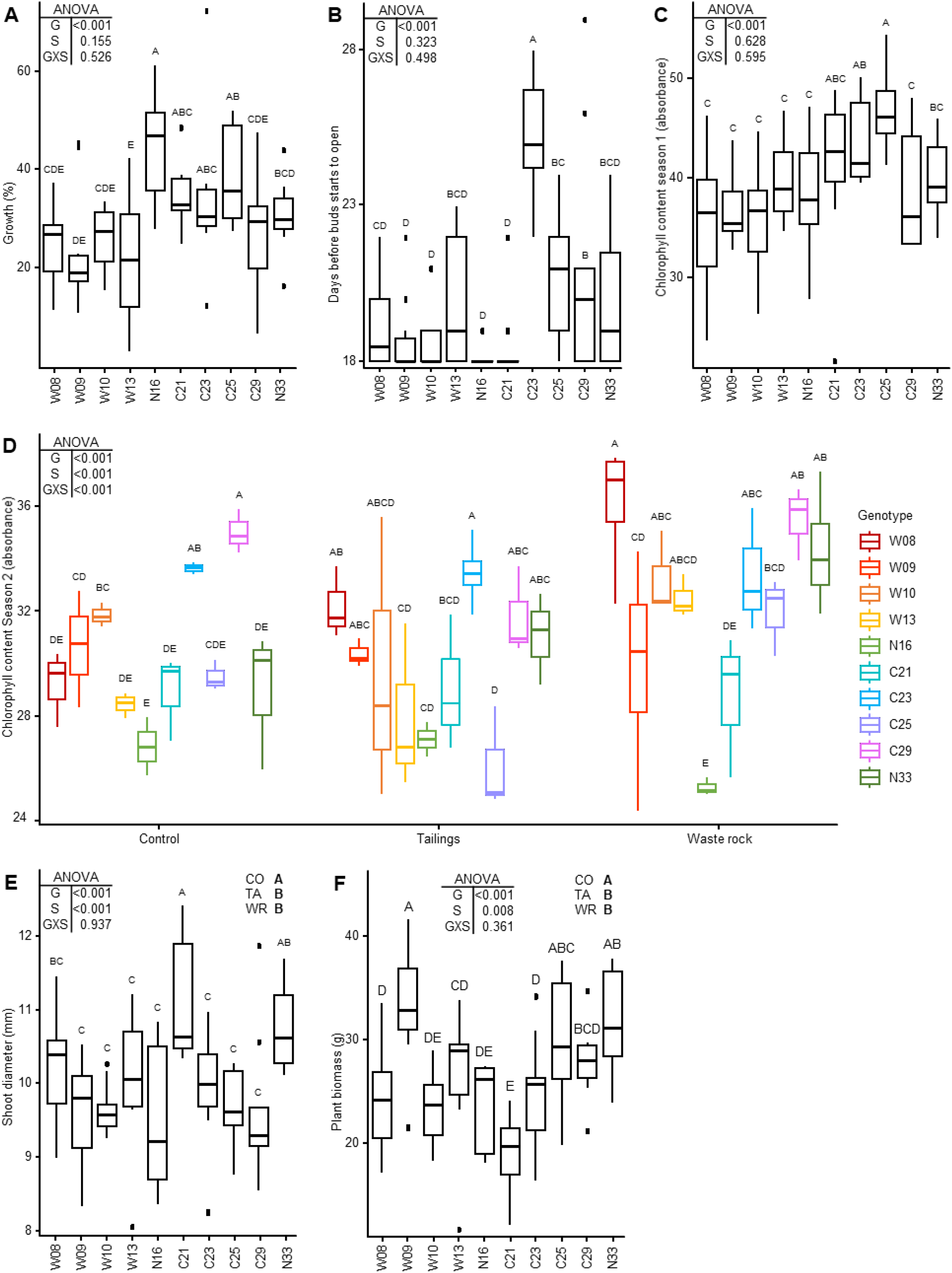
Effect of genotype and substrate type on tree growth measurements. Tree growth during the first season **(A)**; chlorophyll content at the end of the first **(B)** and second **(C)** seasons; blooming during the second season **(D)**; shoot diameter **(E)**; and biomass produced during the second season **(F)**. A two-way ANOVA was used to discern how genotype, substrate type and their interaction influenced tree growth measurements. Shared letters between treatments means there is no significant difference between these treatments as determined by Tukey HSD post-hoc pairwise comparison test (n ≥ 3). In the ANOVA tables, G is genotype; S is substrate type; and GXS is the interaction between genotype and substrate type. In panels **(E)** and **(F)** CO is control; TA is tailings; and WR is waste rock. Significance level is p < 0.05. In panels **(A)**, **(B)** and **(C)**, because only the effect of genotype is significant, measures from all substrate types were pooled. In panel **(D)**, because the interaction of both factors is significant, all treatments were shown separately. In panels **(E)** and **(F)**, pooled measures from all substrate types are shown for each genotype as a boxplot and the effect for each substrate type is shown in the upper right corner of the plots.

The genotypes having lower or higher values strongly differed depending on the measured growth parameter (Figure 3). As an example, genotypes W08, W10, W13 and C29 had lower values for most parameters except for their chlorophyll content after the second season of growth in mine substrates, for which they had the highest values. Besides, genotypes W09 and N16 had the highest biomass and highest growth, respectively, but had lower values for other parameters. Genotype N33 had intermediate to high values for all parameters.

There was an overall effect of cuttings origin (either the Westwood site: W; the La Corne Mine site: C; or the Natural forest: N) on measured growth parameters when the effect of substrate type was not significant (Figure S5): W genotypes grew significantly less than C and N genotypes (Fig. S4A, *p* < 0.001); C genotypes flushed later than W and N genotypes (Fig. S4B, *p* < 0.001); and C genotypes had a higher chlorophyll content during the first season than W genotypes (Fig. S4C, *p* < 0.001).

When the effect of substrate type was significant, there was no obvious association between cuttings origin and their growth in the substrate from which they came from. For example, the genotypes that had a higher chlorophyll content in tailings were not only those originating from the La Corne Mine site (Fig. 3D). For shoot diameter (Fig. 3E) and plant biomass (Fig. 3F), there was no significant difference between mine substrates.

#### 3.2.2 Soil physicochemical properties

A factorial analysis of the physicochemical properties of substrates revealed that all parameters were significantly affected by substrate type, 11 of the 13 parameters had a significant effect of genotype, and the interaction between substrate type and genotype was significant for 10 parameters (Table S4). To better assess the effect of genotype, pairwise comparisons were made on each substrate type separately (Tables 2 and S5). All physicochemical properties were significantly affected by genotype in at least one substrate. Although significant, the differences between genotypes were small, generally resulting in only two genotypes being different from the others.

**Table 2.** Physicochemical properties of substrates after the greenhouse experiment. See Supplementary Table 5 for the other parameters.

|  | C total (%) |  |  | N total (‰) |  |  | P (mg/kg) |  |  |
| --- | --- | --- | --- | --- | --- | --- | --- | --- | --- |
|  | Control | Tailings | Waste rock | Control | Tailings | Waste rock | Control | Tailings | Waste rock |
| W08 | 9.43 ab | 1.47 b | 2.50 abc | 17.8 ab | 0.35 c | 3.09 bcd | 15.7 b | 2.43 | 6.2 b |
| W09 | 8.23 ab | 1.63 ab | 2.56 ab | 17.1 ab | 0.52 c | 3.98 abc | 30.9 ab | 4.12 | 10.9 a |
| W10 | 6.08 b | 1.93 ab | 2.55 abc | 11.7 b | 1.16 bc | 3.19 bcd | 38.0 a | 6.11 | 12.6 a |
| W13 | 10.96 a | 1.83 ab | 3.21 a | 22.2 a | 0.98 bc | 4.87 ab | 34.6 ab | 7.10 | 10.9 a |
| N16 | 10.24 ab | 2.21 ab | 2.25 bc | 21.8 ab | 1.87 abc | 2.74 cd | 27.6 ab | 5.07 | 11.1 a |
| C21 | 10.02 ab | 2.19 a | 1.79 c | 19.3 ab | 1.91 abc | 1.32 d | 33.0 ab | 5.14 | 9.2 ab |
| C23 | 7.60 ab | 2.05 ab | 2.61 abc | 13.9 ab | 1.22 bc | 3.50 abcd | 37.9 a | 6.06 | 11.5 a |
| C25 | 7.81 ab | 2.14 a | 2.21 bc | 15.6 ab | 1.47 bc | 4.39 abc | 35.2 a | 3.96 | 11.9 a |
| C29 | 10.59 a | 1.95 ab | 2.71 ab | 21.1 a | 3.22 a | 5.39 a | 37.3 a | 7.07 | 11.9 a |
| N33 | 7.16 ab | 1.59 ab | 2.22 bc | 14.3 ab | 2.50 ab | 4.29 abc | 29.8 ab | 2.68 | 10.5 a |
| p-value | 0.009 | 0.006 | 0.002 | 0.009 | < 0.001 | < 0.001 | 0.017 | 0.027 | < 0.001 |

|  | K (cmol/kg) |  |  | Ca (cmol/kg) |  |  | Mg (cmol/kg) |  |  |
| --- | --- | --- | --- | --- | --- | --- | --- | --- | --- |
|  | Control | Tailings | Waste rock | Control | Tailings | Waste rock | Control | Tailings | Waste rock |
| W08 | 4.42 cd | 1.14 b | 2.16 | 9.20 c | 1.40 b | 3.56 c | 3.97 | 0.64 b | 0.86 ab |
| W09 | 9.61 abcd | 1.60 ab | 2.20 | 24.80 a | 2.85 ab | 5.93 ab | 7.07 | 1.12 a | 1.28 a |
| W10 | 11.06 a | 2.59 ab | 2.00 | 21.50 abc | 3.45 a | 5.46 abc | 6.43 | 1.45 a | 1.12 ab |
| W13 | 8.75 bc | 1.88 ab | 2.00 | 23.90 ab | 3.49 a | 6.47 ab | 7.26 | 1.30 a | 1.28 a |
| N16 | 7.56 d | 2.22 ab | 1.67 | 20.00 abc | 4.66 a | 4.66 abc | 6.49 | 1.62 a | 1.18 ab |
| C21 | 9.07 abcd | 2.67 a | 1.37 | 19.50 abc | 4.10 a | 3.10 c | 6.13 | 1.45 a | 0.67 b |
| C23 | 10.11 ab | 1.98 ab | 1.72 | 17.60 bc | 3.35 a | 5.99 ab | 5.70 | 1.22 a | 1.24 a |
| C25 | 9.35 abcd | 2.84 a | 1.58 | 20.90 abc | 3.59 a | 4.27 bc | 6.40 | 1.46 a | 1.01 ab |
| C29 | 7.10 d | 1.67 ab | 2.05 | 19.70 abc | 4.06 a | 6.74 a | 5.95 | 1.22 a | 1.28 a |
| N33 | 8.61 abcd | 1.83 ab | 1.57 | 21.10 abc | 3.19 a | 4.88 abc | 5.77 | 1.24 a | 1.06 ab |
| p-value | 0.013 | 0.013 | 0.068 | 0.005 | < 0.001 | < 0.001 | 0.094 | < 0.001 | 0.003 |

|  | pH |  |  | S total (‰) |  |  | Fe (cmol/kg) |  |  |
| --- | --- | --- | --- | --- | --- | --- | --- | --- | --- |
|  | Control | Tailings | Waste rock | Control | Tailings | Waste rock | Control | Tailings | Waste rock |
| W08 | 5.81 | 5.83 | 4.66 ab | 1.04 ab | 0.52 | 10.60 a | 0.450 | 0.910 d | 2.100 |
| W09 | 5.59 | 5.51 | 4.86 a | 1.27 ab | 0.54 | 10.30 a | 1.054 | 1.800 ab | 2.320 |
| W10 | 5.68 | 5.58 | 4.73 ab | 0.62 ab | 0.70 | 12.70 a | 0.869 | 1.890 a | 2.450 |
| W13 | 5.77 | 5.64 | 4.92 a | 1.84 a | 0.14 | 10.40 a | 0.957 | 1.720 abc | 2.170 |
| N16 | 5.64 | 6.03 | 4.84 a | 1.21 ab | 0.05 | 10.20 a | 0.813 | 1.220 cd | 2.130 |
| C21 | 5.39 | 5.61 | 4.32 b | 1.03 ab | 0.89 | 11.20 a | 0.922 | 1.420 bc | 2.660 |
| C23 | 5.63 | 5.64 | 4.83 a | 1.19 ab | 0.48 | 3.40 b | 1.089 | 1.580 abc | 2.700 |
| C25 | 5.65 | 5.48 | 4.66 ab | 0.30 b | 0.11 | 3.50 b | 1.028 | 1.430 abc | 2.140 |
| C29 | 5.54 | 5.62 | 4.92 a | 0.34 b | 0.02 | 2.60 b | 0.973 | 1.650 abc | 2.270 |
| N33 | 5.44 | 5.53 | 4.75 ab | 0.03 b | 0.03 | 4.10 b | 0.998 | 1.780 ab | 2.260 |
| p-value | 0.282 | 0.111 | 0.015 | 0.011 | 0.684 | < 0.001 | 0.354 | < 0.001 | 0.497 |

Genotypes having a significant effect on physicochemical properties varied depending on the substrate type (Table 2). For example, genotypes associated with higher carbon content were W13 and C29 in the control substrate; C21 and C25 in tailings; and W13 in waste rock.

General trends were observable for some genotypes (Table 2). In all substrates, genotypes W08 and N16 were associated with less favorable soil conditions (lower nutrient and higher sulfur content). In waste rock, genotype C21 led to the lowest content of all elements and the lowest pH, but, in tailings, it was associated with higher nutrient content. In both mine substrates, genotype C29 was associated with higher nutrient content and pH.

There was an overall effect of the origin of the cuttings on the physicochemical properties of the substrates. Indeed, several elements showed higher concentrations when genotypes were grown in their original mine waste: in tailings, C (*p* = 0.003), N (*p* < 0.001), Ca (*p* = 0.019) and Mn (*p* = 0.029) contents were higher for genotypes originating from La Corne Mine; in waste rock, C (*p* = 0.022), N (*p* < 0.001) and K (*p* = 0.002) contents were higher for genotypes originating from Westwood. Lastly, plant biomass was significantly, albeit weakly, correlated with N and Na contents of the substrates (Table S6).

#### 3.2.3 Microbiome analyses

##### 3.2.3.1 Alpha-diversity

Factorial analyses of alpha diversity indices indicated that substrate type had a significant effect on bacterial and fungal richness (chao1: *p* < 0.001) and diversity (Shannon: *p* < 0.001 and 0.004; Figure 4). There was a significant interaction between substrate type and genotype on bacterial diversity (*p=* 0.031). Pairwise comparison between substrate types indicated that bacterial richness (*p* < 0.001) and diversity (*p* < 0.001) were significantly higher in tailings compared to waste rock and control substrate, but that fungal richness (*p* < 0.001) and diversity (*p* = 0.001) were higher in the control substrate than in mine substrates. Alpha diversity was analyzed by substrate type to better assess the effect of genotype. Bacterial richness was higher in genotype C29 compared to genotype C21 in the waste rock (*p* = 0.040; chao1 index, Figure 4). There was no effect of genotype on fungal alpha diversity.

**Figure 4.**
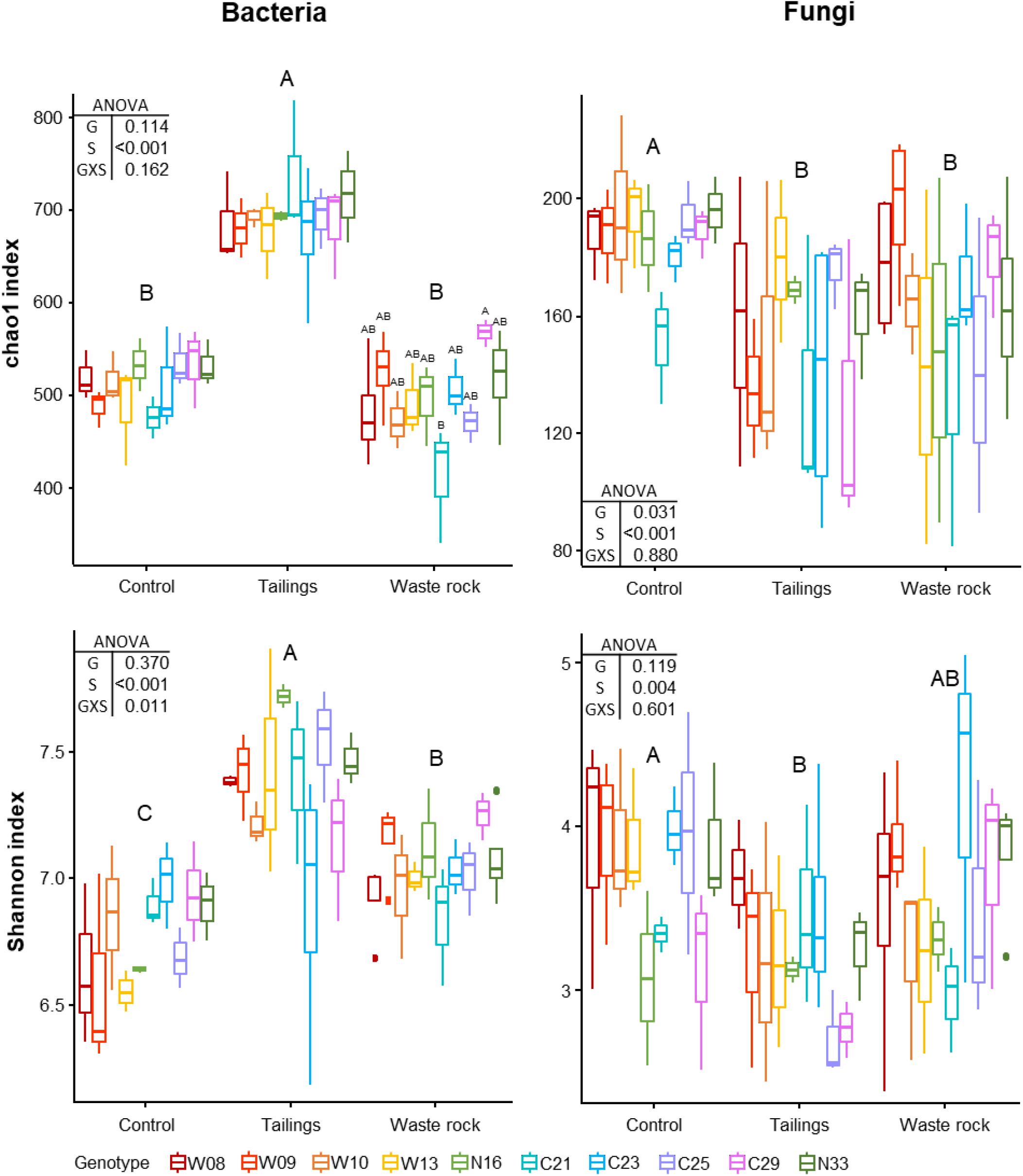
Alpha diversity indices of bacterial and fungal community profiles in greenhouse samples. A two-way ANOVA was used to discern how genotype, substrate type and their interaction influenced alpha diversity indices. In the ANOVA table, G is genotype; S is substrate type; and GXS is the interaction between genotype and substrate type. Shared letters between treatments means there is no significant difference between these treatments, as determined by Tukey HSD post-hoc pairwise comparison test (n ≥ 3). Significance level is *p* < 0.05. Genotype had a significant effect on bacterial richness in waste rock only (chao1 index; *p* = 0.040).

##### 3.2.3.2 Beta-diversity

Bacterial and fungal community structure highly differed between substrate types as shown by variation in beta diversity (Figure 5). The main driver of bacterial and fungal community structure was substrate type (R^2^ = 54.9% and 47.0%, respectively), and a multilevel pairwise comparison test revealed that all substrate types, for bacterial and fungal communities, clustered separately (*p* < 0.001). In bacterial beta diversity, genotype and the interaction between substrate type and genotype, explained, respectively, 7.4% and 11.3% of the variation in rhizosphere community structure. Genotype and the interaction were not significant for the fungal community structure. All physicochemical properties of the substrates were significantly (*p* < 0.001) correlated with bacterial and fungal community structures as shown by the arrows on Figure 5.

**Figure 5.**
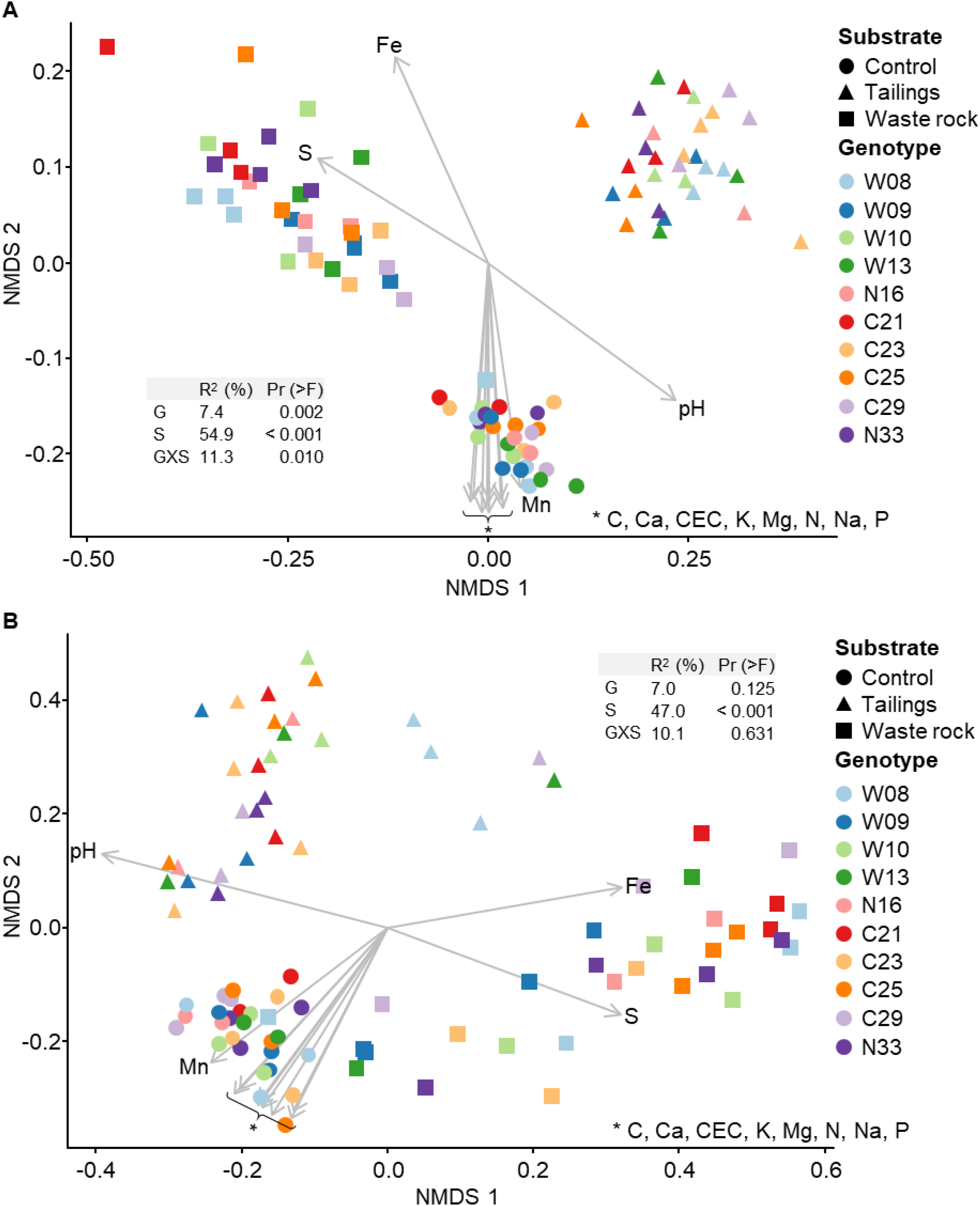
Non-metric multidimensional scaling (NMDS) ordination of variation in bacterial and fungal community structure of balsam poplar rhizosphere in the greenhouse experiment. Bacterial community **(A)** and fungal community **(B)**. Points represent samples and arrows represent the significant (*p* < 0.001) correlations between NMDS axes and the physicochemical properties of the substrates. In PERMANOVA tables, G is for genotype; S for substrate type and GXS for the interaction between genotype and substrate type. The model explains 73.6% and 64.1% of the variation in bacterial and fungal community structure, respectively (n ≥ 3).

Beta diversity was analyzed by substrate type separately to better assess the effect of genotype on bacterial and fungal community structure (Figure S6). There were differences in bacterial community structure between a few genotypes in both mine substrates (Figures S6B, C) and in tailings for fungal community structure (Figure S6E). In all substrates, bacterial and fungal community structure was correlated with at least one physicochemical property (arrows in Figure S6).

##### 3.2.3.3 Taxonomic profiles

Figure 6 illustrates the relative abundance of the most abundant bacterial and fungal taxa (>1%, at the genus level) in the rhizosphere of balsam poplars after two seasons of growth in tailings, waste rock and control substrates. Many bacterial and fungal taxa were only identified at a high taxonomic level.

**Figure 6.**
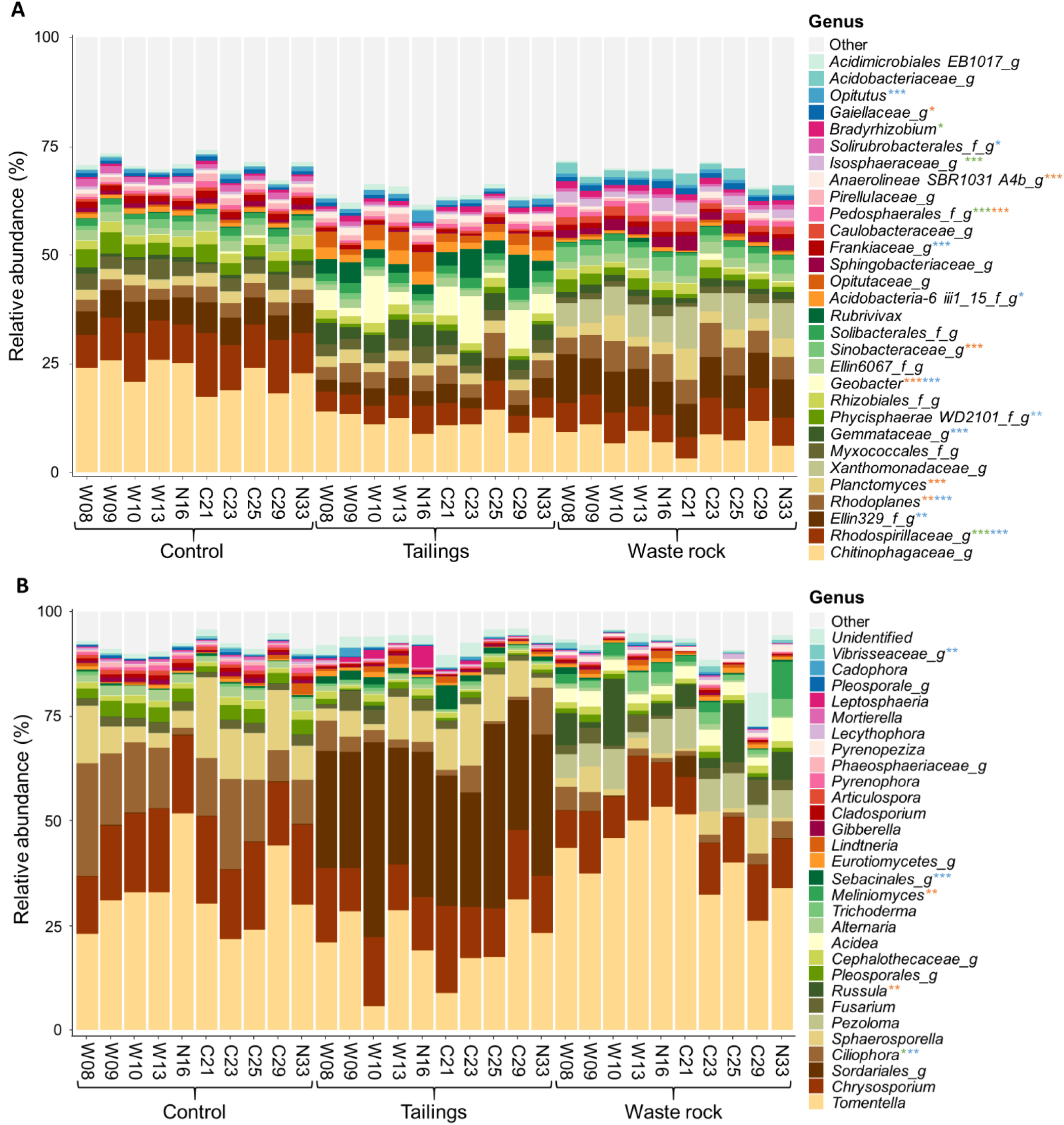
Taxonomic profiles of bacterial and fungal communities in treatments from the greenhouse experiment. Bacterial **(A)** and fungal **(B)** communities at the genus level. Only bacterial and fungal taxa with a relative abundance >1% in at least one treatment are shown. Replicates for each genotype were pooled for visual simplification (n ≥ 3). Stars represent levels of significant difference between genotypes in the three substrates analyzed separately: green for control, blue for tailings and orange for waste rock. Significant differences were determined by a Tukey HSD post-hoc pairwise comparison test. Significance level is represented as follows: *p* < 0.001 ***, *p* < 0.01 **, *p* < 0.05 *.

Functions associated with fungal community of the rhizosphere comprised mostly ectomycorrhizae (40%), saprotrophs (22%), plant pathogens (5%), ericoid mycorrhizae (1%) and brown rot (1%). Factorial analyses of the relative abundance of each function (Table S7) revealed that substrate type had a significant effect on the relative abundance of ectomycorrhizae, saprotrophs, plant pathogens, ericoid mycorrhizae and white rot. Pairwise comparisons between genotypes among each substrate type were performed to better assess the effect of genotype. The effect of genotype was significant on the relative abundance of ericoid mycorrhizae in waste rock: they were more abundant in the rhizosphere of genotype N33 compared to genotypes W09, C25 and C29 (*p* = 0.005).

Factorial analyses of taxa abundance in bacterial and fungal communities (for taxa >1%, Table S8) showed that the effect of substrate type was significant for all bacterial taxa and most fungal taxa (24/29); the effect of genotype was significant for many bacterial and fungal taxa (17/30 and 7/29, respectively); and the interaction between substrate type and genotype was significant for a few bacterial and fungal taxa (13/30 and 5/29, respectively). Pairwise comparisons between genotypes among each substrate type were made to better assess the effect of genotype. The effect of genotype was significant in at least one substrate for half of bacterial taxa (17/30) and for a few fungal taxa (5/29). The bacterial and fungal taxa significantly affected by genotype were marked with stars in Figure 6.

The relative abundance of most bacterial and fungal taxa was significantly correlated with physicochemical properties of substrates (Table S9). In waste rock, bacterial and fungal communities were characterized by acid tolerant taxa like the bacterial family *Xanthomonadaceae* (Callender *et al*., 2016) and the fungal genus *Acidea* (Hujslová and Gryndler, 2019), while the tailings were dominated by microorganisms tolerant to stress like the oligotrophic bacterial genus *Geobacter* (Wilkins *et al*., 2008). As for the community composition of the control substrate, it was mostly composed of decomposer microorganisms like the bacterial family *Chitinophagaceae* (Rosenberg, 2014) and the fungal genus *Chrysosporium* (Tedersoo *et al*., 2014). A few plant growth parameters were weakly correlated (|0.3| < r < |0.5|) with the relative abundance of a few bacterial and fungal taxa (Table S10).

There was an overall effect of the origin of the cuttings on the relative abundance of bacterial and fungal taxa (Table S11). For example, the bacterial genus *Bradyrhizobium* was more abundant in the rhizosphere of genotypes originating from the La Corne Mine site compared to the genotypes originating from the Westwood site, in the control substrate (*p* = 0.030).

## 4 Discussion

### Vegetation improved physicochemical properties of mine wastes *in situ*

In this study, the impact of naturally grown *P. balsamifera* on two contrasting mine wastes was assessed *in situ*. These mine wastes were considered unfavorable for plant growth because they contain only small concentrations of essential nutrients, have either a very low or very high pH and have poor physical structure and deficient water holding capacity. The pioneer tree *P. balsamifera* has previously been found to naturally grow on mine sites (van Haveren and Cooper, 1992), a phenomenon that was also observed in this study on the highly distinct mine wastes from our two sites. On both sites, well established vegetation significantly improved most of the physicochemical properties of mine wastes, with an increase in carbon and nutrient (N, K, P, Ca and Mg) content, pH values closer to the natural forest soil samples, and a decrease in S and Fe concentrations in waste rock.

Increase in C and N comes from organic matter provided by the growth of balsam poplars; organic matter is further degraded by heterotrophic microorganisms in the soil. This increase in organic matter is responsible for the variations in pH as well as S and Fe. Indeed, soil organic matter buffers soil pH by binding to H^+^ in acidic soil. Similarly, Fe binds to organic matter making it less available. Consequently, it has been shown that S adsorption decreases when pH is higher, Fe and Al oxides contents are lower and organic matter content is higher, leading to S uptake by plants and leaching (Johnson *et al.*, 1992).

### Vegetation caused a beneficial shift in microbial communities of mine wastes

Results from the field experiments are in line with other studies (Chen *et al.*, 2013; Li *et al.*, 2015, 2016) that found a succession of microbial communities shifting from lithotrophic to heterotrophic microorganisms during plant growth on mine wastes. These shifts in microbial community structure suggest that initial soil conditions of mine wastes favoring the growth of lithotrophic microorganisms changed during the establishment of balsam poplars on site, confirming the positive influence of these trees on microbial community structure, function and ecosystem health. For example, *Leptospirillum* and *Acidiphilium*, two bacterial genera associated with the oxidoreduction of iron and acid mine drainage (Harrison Jr, 1981; Hippe, 2000), as well as *Sulfobacillaceae*, a bacterial family associated with the oxidation of sulfur and acid mine drainage (Hottenstein *et al.*, 2019), were found to be more abundant in unvegetated than in vegetated zones of the waste rock pile. A previous study also reported that vegetation growth on acid mine tailings lowered the abundance of these key iron and sulfur oxidizing bacteria and lowered acidity (Li *et al.*, 2016).

Surprisingly, the presence of vegetation on mine wastes also reduced the relative abundance of fungal taxa typically known as plant pathogens, suggesting that these fungi may have other ecological functions in disturbed lands. Some of these taxa, like *Alternaria*, *Ganoderma* and the *Teratosphaeriaceae* family, have also been previously isolated in mine wastes and various acidic environments (Wong, 1981; Hujslová *et al.*, 2013; Callender *et al.*, 2016; Mosier *et al.*, 2016). It was also surprising that the presence of vegetation on mine wastes reduced the relative abundance of fungal saprotrophs, as it would be expected that an increase in organic matter would also increase the presence of these microorganisms. These results illustrate a well-known limitation of the use of relative abundances in metabarcoding studies (Zhang *et al.*, 2017; Lin *et al.*, 2019). Although the relative abundance of saprotrophs was lower in the vegetated soil samples, the absolute abundance of these microorganisms might still be higher than in unvegetated mine wastes. This issue could be avoided in further studies by using quantitative PCR to estimate the total populations in these environments (Rastogi *et al.*, 2010) or by spiking exogenous bacteria, fungi or synthetic DNA prior to sample processing (Tourlousse *et al.*, 2017).

Furthermore, many rhizobacteria of the orders *Rhizobiales*, *Sphingomonadales* and *Burkholderiales* and the phylum *Planctomycetes*, *Bacteroidetes*, *Actinobacteria* and *Acidobacteria* were found to be more abundant in vegetated soil samples than in unvegetated mine wastes; however, at lower taxonomic levels, the taxa detected differed between waste rock and tailings. These taxa have previously been associated with the rhizosphere microbiome (da Rocha *et al.*, 2013; McBride *et al.*, 2014; Madhaiyan *et al.*, 2015; Qiao *et al.*, 2017); plant growth promotion (e.g. through nitrogen fixation (Caballero-Mellado *et al.*, 2004; Dai *et al.*, 2014; Sun *et al.*, 2015; Jeanbille *et al.*, 2016); the production of IAA (Mehnaz *et al.*, 2010); or disease suppression (Xue *et al.*, 2015)), and nutrient cycling (Webb *et al.*, 2014; Santoyo *et al.*, 2016; Wu *et al.*, 2017). Similarly, vegetation increased the relative abundance of ectomycorrhizal taxa, such as *Meliniomyces*, which have previously been isolated from poplars growing in mine wastes (Gaster *et al.*, 2015; Katanić *et al.*, 2015).

For both mine sites, *Proteobacteria* were more abundant and the *Proteobacteria*-to-*Acidobacteria* ratio was higher in vegetated soils than in unvegetated mine wastes. This corroborates previous studies showing that this ratio is an indicator of soil trophic levels, and for which *Proteobacteria* were linked to nutrient-rich soils and *Acidobacteria* to nutrient-poor soils (Fierer *et al.*, 2007; Castro *et al.*, 2010). Gottel *et al.* (2011) found similar results in the rhizosphere of *Populus deltoides* in which *Proteobacteria* were slightly more prevalent than *Acidobacteria*.

### Vegetation increased bacterial richness and diversity

Mine site restoration aims to mitigate the negative impacts of mining on the environment and human health. However, land restoration is a long process since the affected ecosystems have lost their plant and microbial biodiversity and most of their functions and services (Prach and Tolvanen, 2016). In this study, vegetation increased bacterial richness and diversity in mine substrates, which suggests an improvement in ecosystem productivity and stability (Tilman *et al.*, 2006). On the other hand, a decrease in fungal richness and diversity was observed in the vegetated soils compared to the mine wastes. This might be due to the competitive exclusion of ectomycorrhizal fungi on other fungi, particularly plant pathogens, corroborating other studies that have shown that disturbed lands have a greater fungal diversity than forested lands (Ding *et al.*, 2011).

### Substrate type has a stronger effect on community composition than genotype

In the greenhouse experiment, substrate type was shown to be the main driver of bacterial and fungal community structure and diversity in the rhizosphere of balsam poplar. This is a consistent finding among studies about the *Populus* root microbiome (Gottel *et al.*, 2011; Bonito *et al.*, 2014; Veach *et al.*, 2019) and other plant species (Marschner *et al.*, 2004; Lebeis *et al.*, 2015; Wagner *et al.*, 2016; Colin *et al.*, 2017; Gallart *et al.*, 2018), indicating that larger-scale edaphic conditions primarily regulate overall rhizosphere microbiomes. Physicochemical properties, including granulometry and pH, and other unmeasured factors, like water holding capacity and variations in temperature and rain between seasons, contribute to those larger-scale edaphic conditions and likely play a role in the *Populus* root microbiome assembly (Chaparro *et al.*, 2012; Philippot *et al.*, 2013). Nevertheless, in this study, plant genotype also influenced physicochemical properties, but this indirect effect was too weak to be reflected on the community assembly. Furthermore, the effect of genotype varied between substrate types, as shown by the significant interaction between both factors for most bacterial and fungal taxa. Bonito *et al.* (2019) also found that soil origin and properties structured microbial communities to a greater degree than host genotype, and that OTUs enriched in genotype samples vary based on the soil properties in which the genotype was grown. This confirms our expectations of a weak genotype effect and indicates that OTUs that are enriched in a sample cannot be used to discriminate plant genotype.

### Tree genotype has low effect over fungal community structure, diversity and functional prevalence compared to bacterial community

Similarly to the results obtained for the field experiment, diversity and structure of fungal community are much more conserved between treatments compared to bacterial community. Tree genotype had a significant effect on the relative abundance of 17 of the 30 most abundant bacterial taxa in at least one substrate type but only 5 of the 29 most abundant fungi. Additionally, while all bacterial taxa were affected by substrate type, five fungal taxa were not. Fungal guild designations revealed that there was barely any effect of genotype on the relative abundance of fungal functions: the effect of genotype was only significant for the relative abundance of ericoid mycorrhizae in waste rock. This could be explained by the fact that plants recruit taxonomic groups in order to balance functions (Maherali and Klironomos, 2007). In other words, different genotypes may select different taxa but ultimately they select similar functions.

### Tree genotype and its associated microbiome could be linked to improvement of physicochemical properties of substrates

Overall, the genotype C29 was associated with higher nutrient content in both mine substrates when compared with the control substrate. This suggests that some genotypes could be selected to improve the physicochemical properties of a broad range of substrate types. Conversely, growth of genotypes W08 and N16 generally led to less favorable physicochemical properties (i.e. lower carbon and nutrient content, lower pH in waste rock) in both mine substrates, suggesting that they are both ill-adapted for revegetation purposes. Moreover, growth of genotype C21 led to the most significant improvement of the physicochemical properties in tailings, but not in waste rock. Interestingly, genotype C21 was collected at the La Corne Mine site, which may suggest a fine scale local adaptation of this genotype that could be further investigated (Boshier *et al.*, 2015).

Furthermore, trends in physicochemical properties could be associated with the rhizosphere bacterial richness. Indeed, in waste rock, genotype C29 was associated with higher nutrient content and pH as well as chao1 index of alpha diversity compared to C21, which was associated with the lowest nutrient content, the lowest pH and the lowest chao1 index. These results suggest that an increase in microbial diversity can lead to improved soil health (Garbeva *et al.*, 2004). Additionally, some dominant taxa could be associated with more favorable conditions of the substrates under poplars of particular genotypes. For example, *Rhodoplanes* was previously found in the rhizosphere of rice paddy soils irrigated by acid mine drainage contaminated water and it has been suggested that they may have a beneficial ecological function to enhance soil fertility (Sun *et al.*, 2015). Interestingly, in our study, they were more abundant in the rhizosphere of genotype C23 (8%) compared to genotype W08 (4%) in waste rock, and nutrient content in the pots of genotypes C23 was also higher compared to W08, supporting the idea that they may play a role in soil health.

### Further studies will need to assess the degree of standing genetic variation with harsher treatments

There was an overall effect of cutting origin on growth measurements, physicochemical properties of the substrates and abundance of some bacterial and fungal taxa in each substrate type. This effect could suggest that there may have been a fine scale adaptation of the genotypes in these novel environments due to high selective environmental pressures. Further research may better assess the effect of this fine scale adaptation on the composition of the microbiome of poplar trees in such novel environments.

Previous studies have shown that changes in the presence-absence or abundance of just a few microbial taxa can affect plant performance because of their broad functions (Zolla *et al.*, 2013; Henning *et al.*, 2016). In this study, it was shown that plant genotype had a significant effect on the abundance of some bacterial and fungal taxa, but there was no obvious correlation between the rhizosphere microbiome and tree growth. This may be because of the relatively lenient conditions of our treatments. Indeed, the mine substrates were amended with a peat mix to help plant growth and reduce stress caused by acidity, non-proper hydric conditions and low nutrient content. Nonetheless, tree growth measurements during the second season (shoot diameter, plant biomass and chlorophyll content) were also influenced by substrate type, showing that longer exposition to the mine substrates could amplify the effect of genotype on the rhizosphere microbiome. Further investigation involving harsher treatments, either directly on the field or with non-amended mine substrates in a greenhouse experiment, would be necessary to better assess the effect of genotype-by-environment interactions on plant performance and its associated microbiome.

## Conclusion

This study has shown that balsam poplars are able to improve soil health of mine wastes in field conditions. Moreover, in greenhouse experiments we demonstrated the effect of genotype-by-environment interactions on the structure and diversity of the rhizosphere microbiome as well as the physicochemical properties of the soil. Our results highlight the influence of balsam poplar genotype in contrasting substrate types. We provided evidence that (1) balsam poplars are suitable to initiate a community of microorganisms closer to a functional vegetated ecosystem on various types of mine wastes as well as increasing soil nutrient content and improving pH; (2) substrate type has a stronger effect on rhizosphere microbial community composition than genotype; (3) nevertheless, plant genotype can act as a selective pressure in structuring rhizosphere microbial communities, particularly bacterial taxa; and (4) genotype-by-environment interactions have an impact on the physicochemical properties of substrates and the composition of the rhizosphere microbiome.

From a practical point of view, the selection of tree genotypes together with associated microbiomes benefitting their growth in mine wastes is a strategy that could facilitate the ecological restoration of mine sites. This study also highlights the importance of microbial criteria to assess the success of revegetation, as shown by the major changes in microbial community structure and diversity. Future research efforts should take into consideration the interdependence between host identity and associated microbiomes in forest ecosystems, in order to better understand plant-soil feedbacks as well as incorporate microbiome community ecology into mining restoration strategies. Our results contribute to the understanding of the relationships between tree genetics and the associated microbial communities and highlight the potential for host genotype-by-environment interactions to shape the composition of host-associated microbial communities. This study confirms the importance of large-scale conditions and environmental heterogeneity on driving soil microbiome assembly, but additionally validates the contribution of plant host genotype in acting as a selective pressure in the surrounding rhizosphere soil.

## 5 Acknowledgements

We wish to acknowledge the contribution of Janet Condie for DNA sequencing at the Illumina Sequencing Platform, National Research Council Canada-Saskatoon. We would also like to thank Éric Dussault for technical assistance provided in the collection of field samples, as well as Serge Rousseau and David Paré for their assistance with soil analyses. We are also grateful to Marie-Claude Gros-Louis for her contribution to the poplar genetic analyses and Gervais Pelletier for support with the greenhouse experiment. The content of this article first appeared in the first author’s master’s thesis (Rheault, 2020).

## 6 Author Contributions

KR, DL, MG and AS contributed conception and design of the study; KR, DL and MJM contributed to acquisition of data; KR and CM performed the statistical analyses; KR, ET, CM and AS contributed to interpretation of data; KR wrote the first draft of the manuscript; CM and NI wrote sections of the manuscript. All authors contributed to manuscript revision, read and approved the submitted version.

## 7 Conflict of Interest

The authors declare that the research was conducted in the absence of any commercial or financial relationships that could be construed as a potential conflict of interest.

## 8 Contribution to the Field

Many studies have attempted to characterize the root microbiome of *Populus*. It has been shown that soil type is the main driver of microbial community assembly, since physicochemical properties influence microbial composition and functional group prevalence. However, genetic variations of the host plant are also associated with differential microbial colonization. Differentiating between the effects of soil properties and those of the host plant genotype has not been sufficiently addressed. Our results contribute to the understanding of the relationships between tree genetic background and the associated microbial communities and highlight the potential for host genotype-by-environment interactions to shape the composition of host-associated microbial communities. This study confirms the importance of large-scale conditions and environmental heterogeneity on driving soil microbiome assembly, but additionally validates the contribution of plant host genotype in acting as a selective pressure in the surrounding rhizosphere soil. Initiatives using *P. balsamifera* as a candidate for abandoned mine sites restoration may need to consider the interplay between genotype and the belowground microbiome. Further examination of microbial community dynamics over longer exposition of trees to mine substrates and other degraded lands may provide a clearer understanding of genotype-by-environment interactions. This knowledge will enable the development of more efficient and effective land reclamation strategies.

## 9 Funding

Financial support was provided by the Fonds de recherche du Québec - Nature et technologies (FRQNT) and the Natural Sciences and Engineering Research Council of Canada (NSERC) for student scholarships to KR. This work was also made possible through funding by the Genomics Research and Development Initiative of Canada and the Ecobiomics Project: Advancing Metagenomics Assessment of Soil Health and Freshwater Quality (https://doi.org/10.1016/j.scitotenv.2019.135906).

## 11 Data Availability Statement

The Illumina data generated in this study was deposited in the NCBI Sequence Read Archive and is available under the project number PRJNA615167.

## Footnotes

1 https://support.illumina.com/content/dam/illumina-support/documents/documentation/chemistry_documentation/16s/16s-metagenomic-library-prep-guide-15044223-b.pdf

## Notes

### Competing Interest Statement

The authors have declared no competing interest.

