## Supplementary Figure 1 for "Plant Genotype Influences Physicochemical Properties of Substrate as well as Bacterial and Fungal Assemblages in the Rhizosphere of Balsam Poplar"

**Supplementary Figure 1.** Pictures of the Westwood site. Sampling of waste rock **(A)**; poplars growing on waste rock **(B)**; screenshot from Google Map of the Westwood site **(C)**; vegetated compared to unvegetated mine waste **(D)**.

**A** **B** **C** **D**

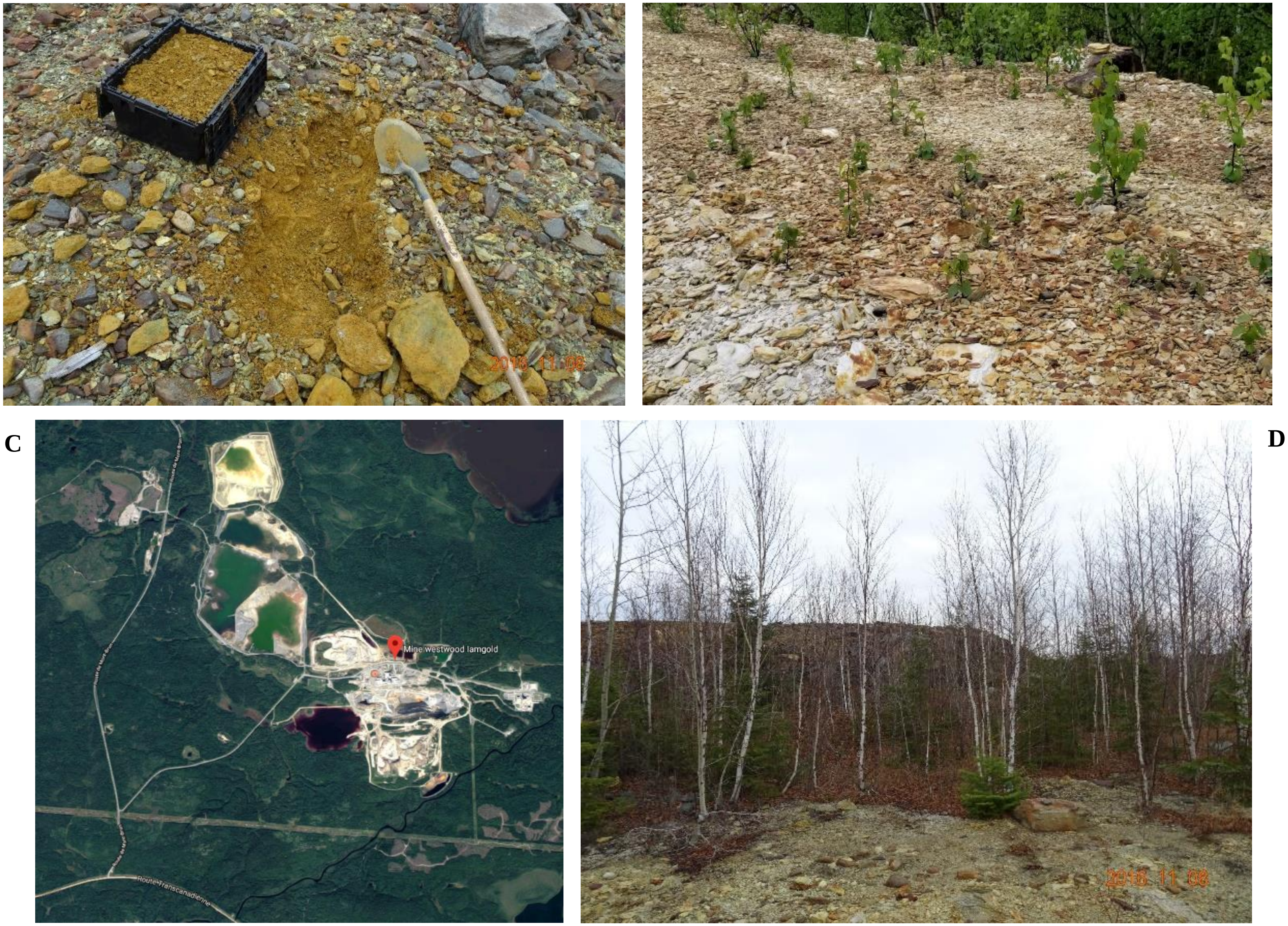
