## Supplementary Figure 2 for "Plant Genotype Influences Physicochemical Properties of Substrate as well as Bacterial and Fungal Assemblages in the Rhizosphere of Balsam Poplar"

**Supplementary Figure 2.** Pictures of the La Corne Mine site. Vegetated compared to unvegetated mine waste **(A)**; sampling of tailings **(B)**; screenshot from Google Map of the La Corne Mine site **(C)**; large view of the vegetation growing in tailings **(D)**.

A

B

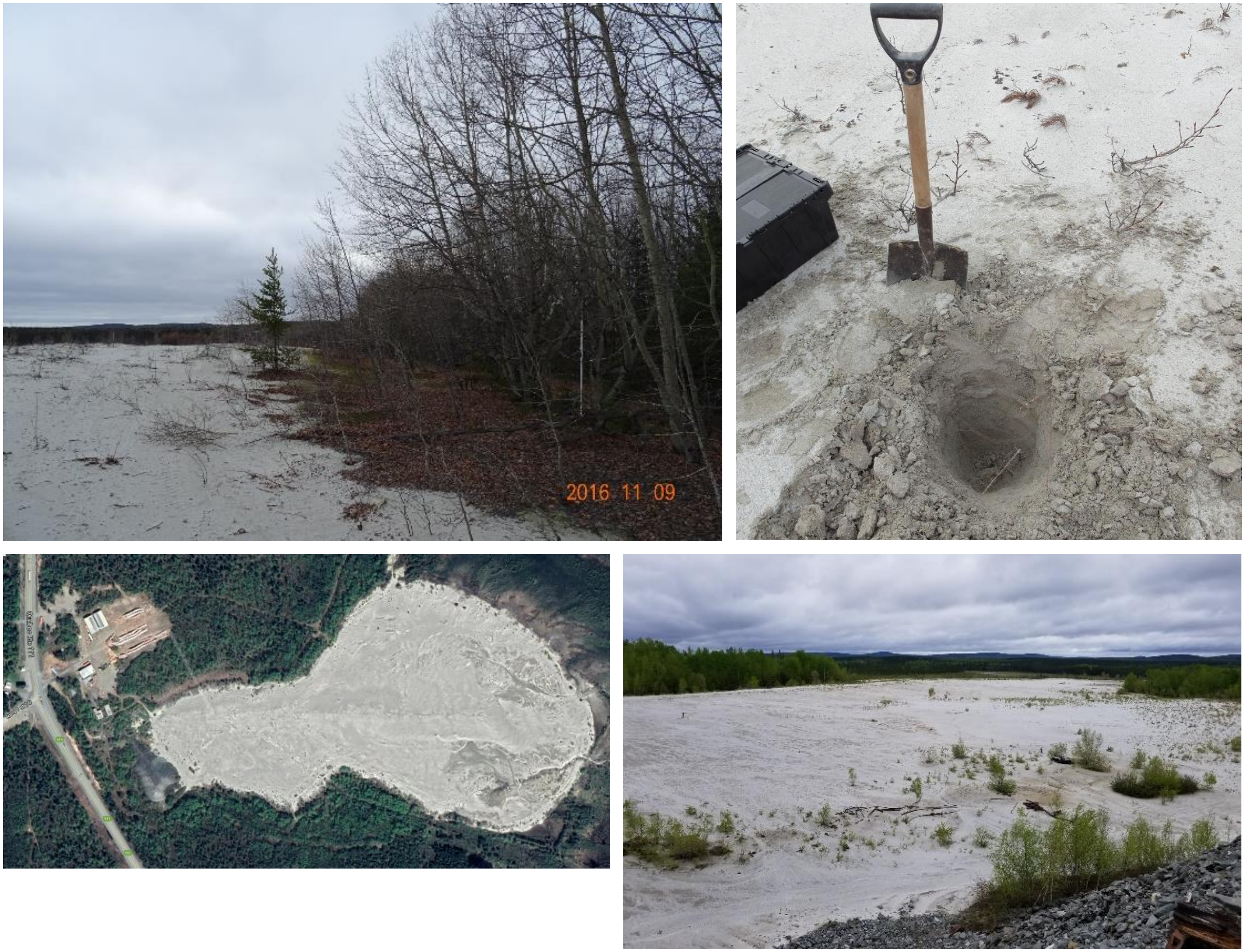


C

D
