## Supplementary Figure 3 for "Plant Genotype Influences Physicochemical Properties of Substrate as well as Bacterial and Fungal Assemblages in the Rhizosphere of Balsam Poplar"

### Field caracterisation experiment

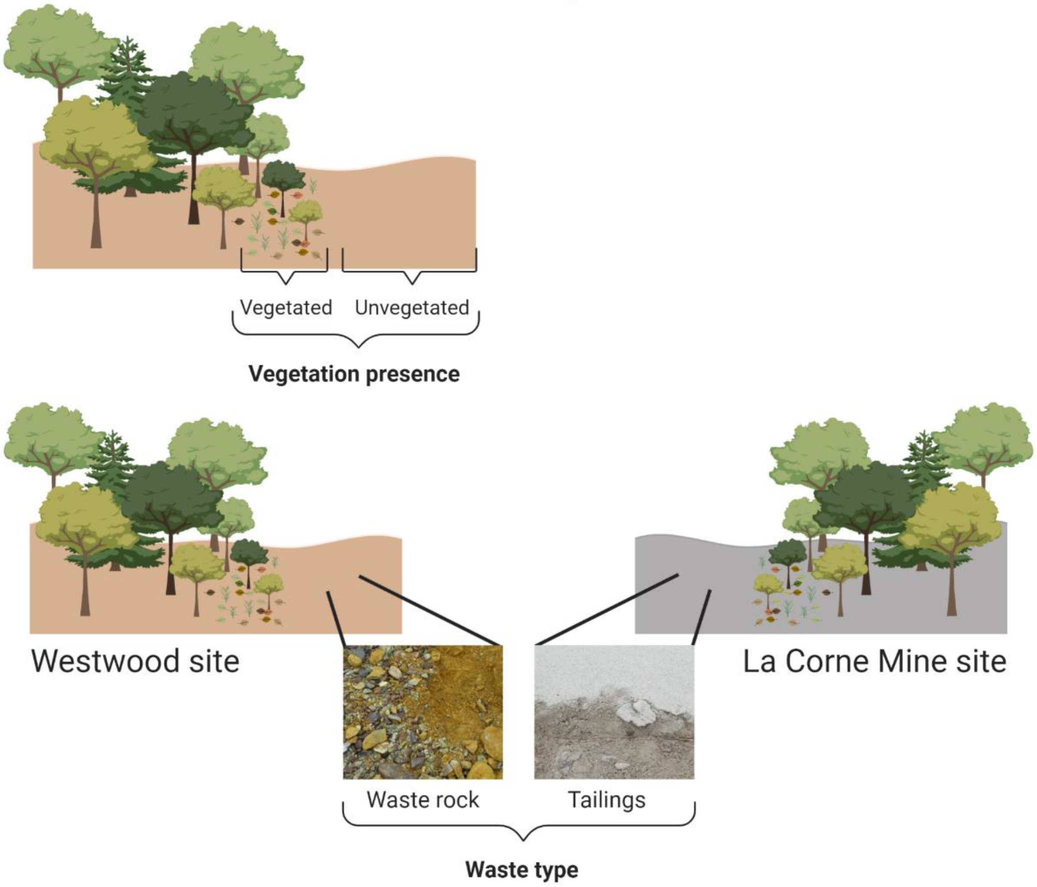


### Greenhouse experiment

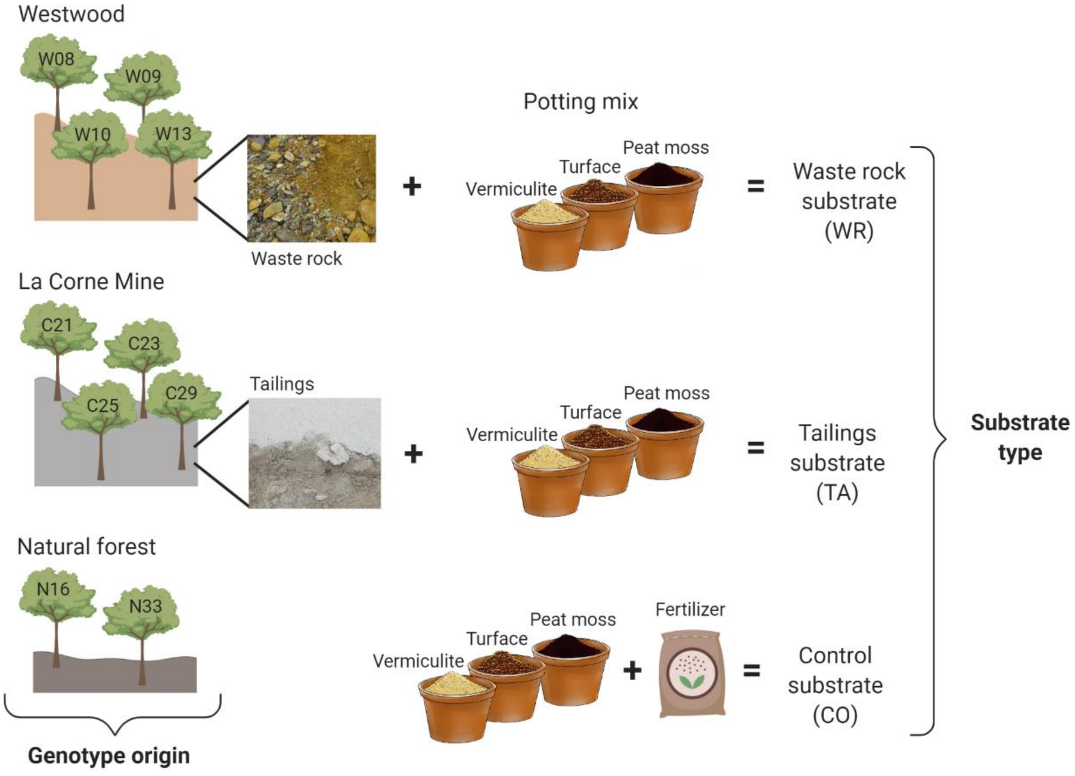

**Supplementary Figure 3.** Illustration of the factors in each experiment. Vegetation presence (unvegetated or vegetated) and waste type (waste rock or tailings) for the field experiment. The origin of the 10 genotypes (2 from a natural forest, 4 from Westwood site and 4 from La Corne Mine site) and substrate type (waste rock, tailings or a control substrate) for the greenhouse experiment.
