## Supplementary Figure 4 for "Plant Genotype Influences Physicochemical Properties of Substrate as well as Bacterial and Fungal Assemblages in the Rhizosphere of Balsam Poplar"

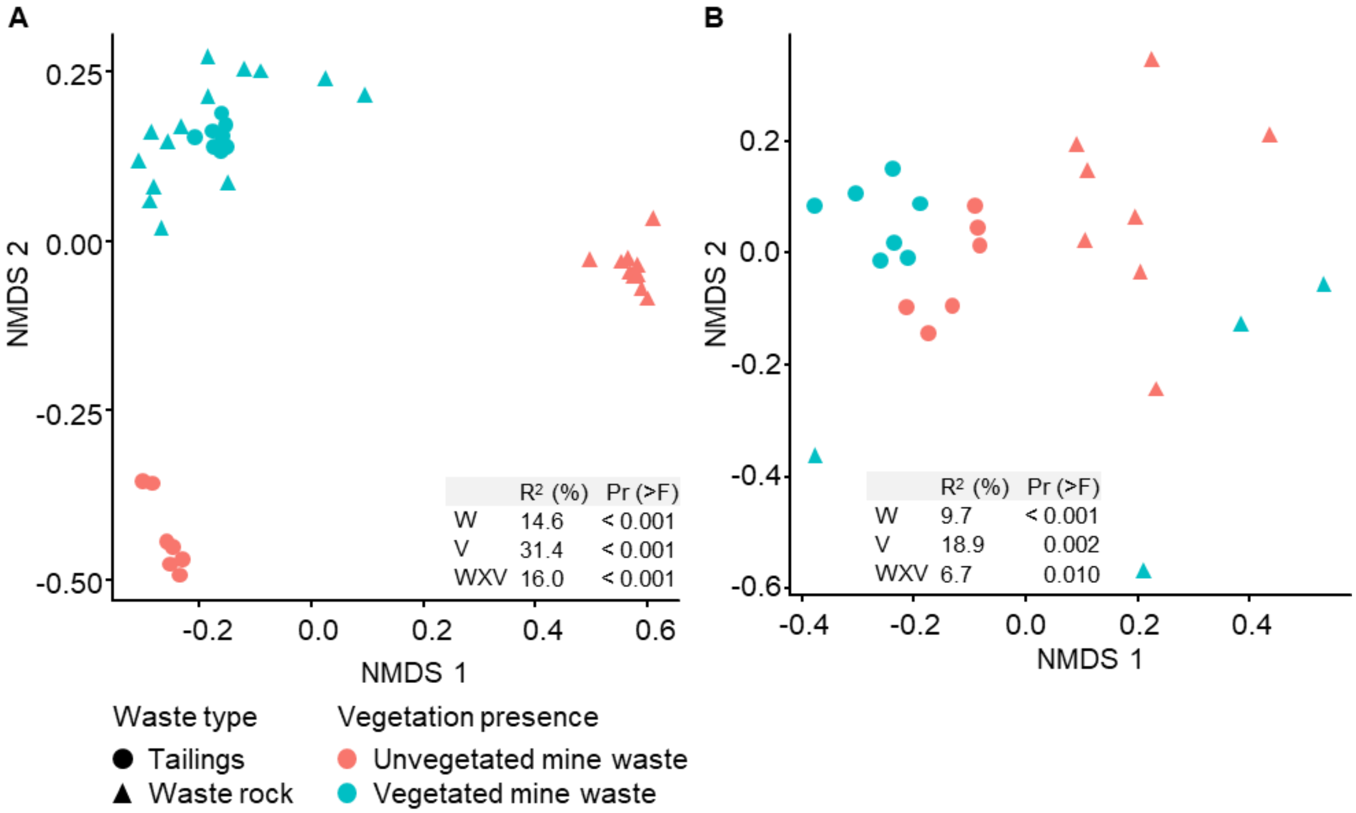


**Supplementary Figure 4.** Non-metric multidimensional scaling (NMDS) ordination of variation in bacterial and fungal community structure of mining substrates in the field experiment. Bacterial community **(A)** and fungal community **(B)**. A permutational multivariate analysis of variance model was implemented to discern the amount of variation attributed to each factor and their interaction ($n \geq 8$). In PERMANOVA tables, W is for waste type; V is for vegetation presence and WXV for the interaction between waste type and vegetation presence. A multilevel pairwise comparison test revealed that all treatments clustered separately for bacterial communities ($p <$ 0.001) and fungal communities ($p < 0.01$).
