## Supplementary Figure 5 for "Plant Genotype Influences Physicochemical Properties of Substrate as well as Bacterial and Fungal Assemblages in the Rhizosphere of Balsam Poplar"

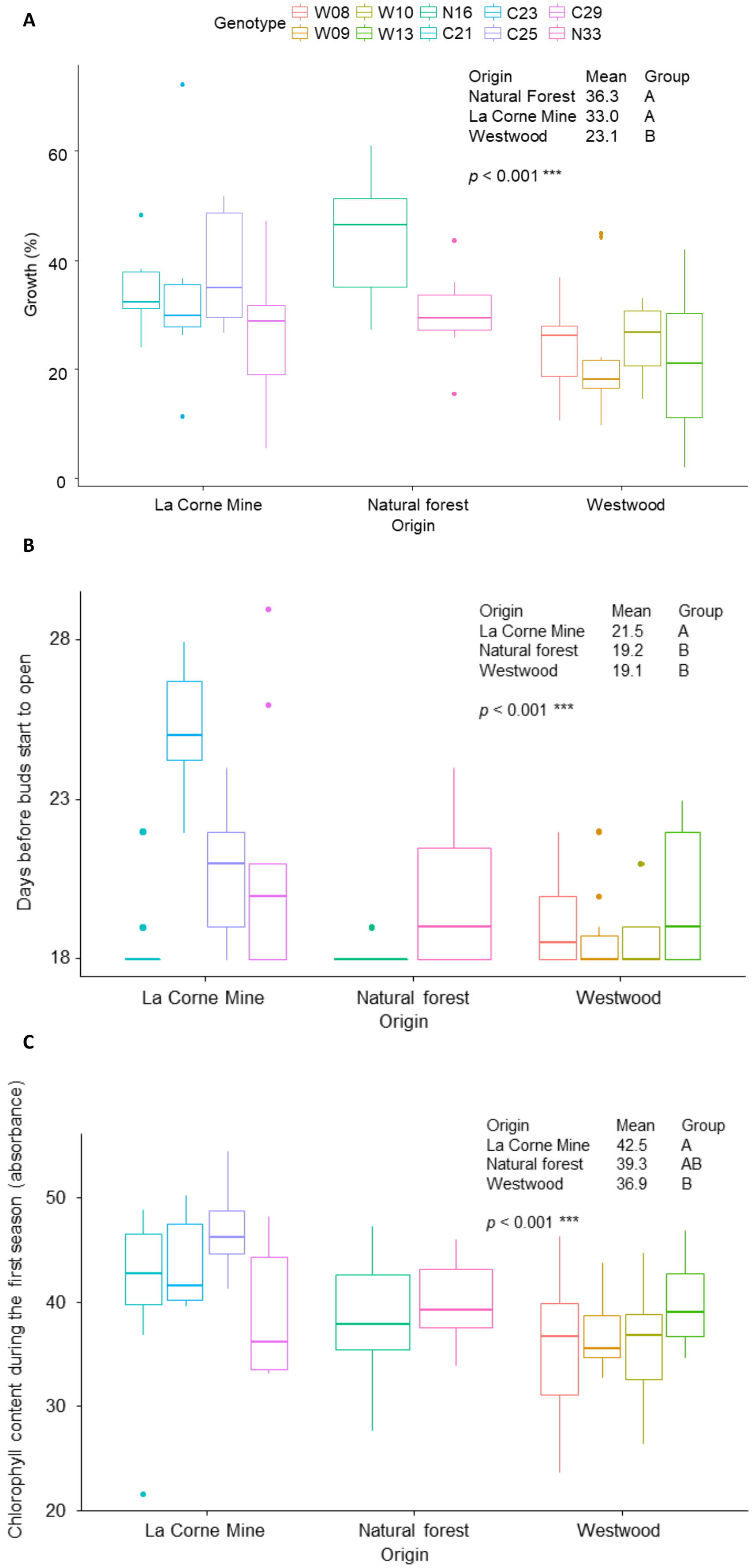
A
Genotype
W08
W09
W10
W13
N16
C21
C23
C25
C29
N33
Origin Mean Group
Natural Forest 36.3 A
La Corne Mine 33.0 A
Westwood 23.1 B
$p < 0.001$ ***
Growth (%)
0
20
40
60
La Corne Mine
Natural forest
Westwood
Origin
B
Origin Mean Group
La Corne Mine 21.5 A
Natural forest 19.2 B
Westwood 19.1 B
$p < 0.001$ ***
Days before buds start to open
18
23
28
La Corne Mine
Natural forest
Westwood
Origin
C
Origin Mean Group
La Corne Mine 42.5 A
Natural forest 39.3 AB
Westwood 36.9 B
$p < 0.001$ ***
Chlorophyll content during the first season (absorbance)
20
30
40
50
La Corne Mine
Natural forest
Westwood
Origin

**Supplementary Figure 5.** Effect of the origin of cuttings on their growth during the first season. Tukey HSD post-hoc pairwise comparison tests were used to discern how the origin of cuttings influenced their growth ($n \geq 3$). Measurements from the first season only are shown because the origin of cuttings did not have a significant effect on tree growth measurements for the second season.
