## Supplementary Figure 6 for "Plant Genotype Influences Physicochemical Properties of Substrate as well as Bacterial and Fungal Assemblages in the Rhizosphere of Balsam Poplar"

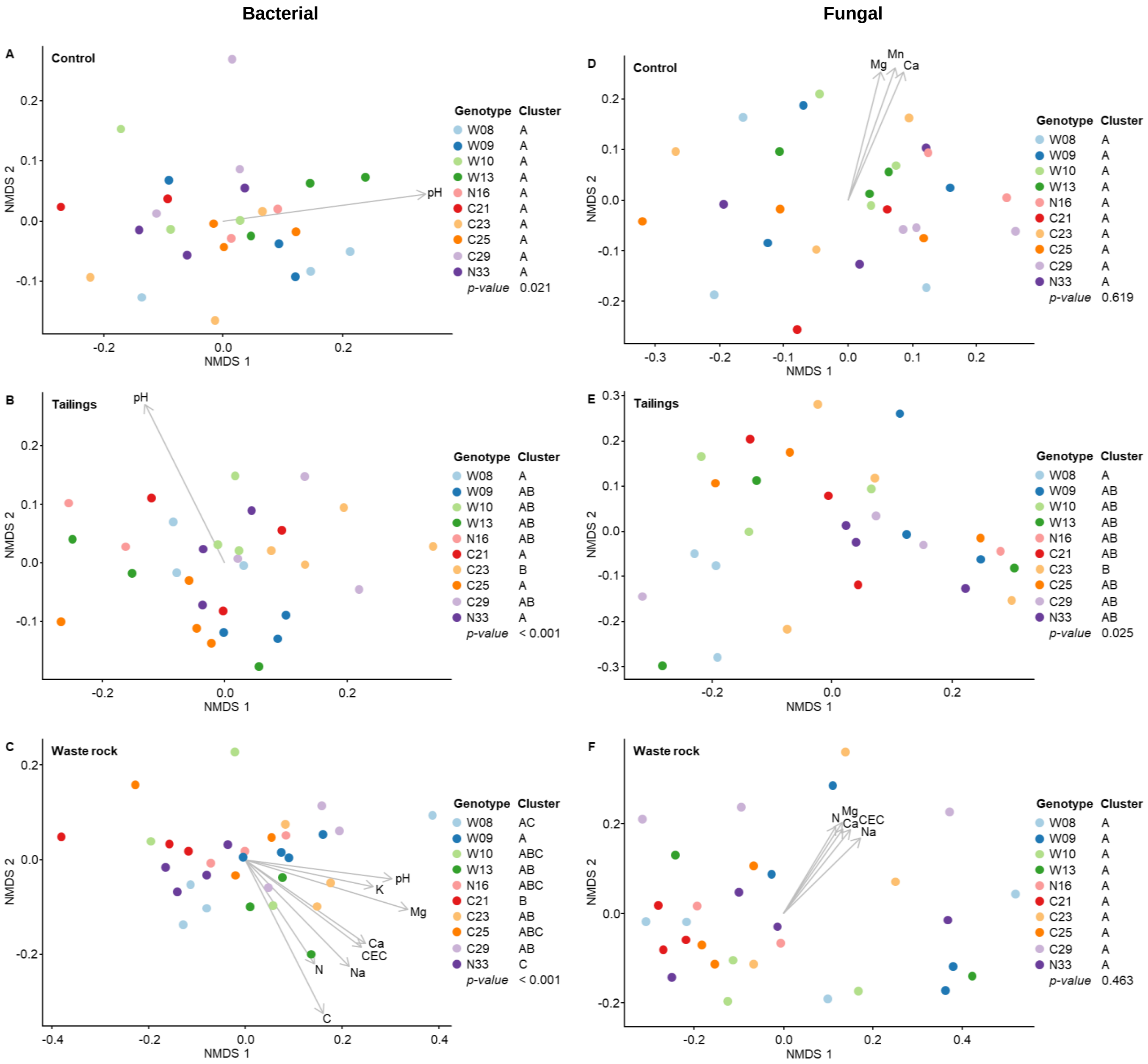


**Supplementary Figure 6.** Non-metric multidimensional scaling (NMDS) ordination of variation in bacterial and fungal community structure in substrates from the greenhouse experiment. Bacterial community structure in control substrate **(A)**, Tailings **(B)** and Waste rock **(C)** and fungal community structure in control substrate **(D)**, Tailings **(E)** and Waste rock **(F)**. Points represent samples and arrows represent the significant correlations between NMDS axes and the physicochemical properties of the substrates. Clusters between genotypes were determined by a multilevel pairwise comparison test (n ≥ 3).
