## Supplementary Table 1 for "Plant Genotype Influences Physicochemical Properties of Substrate as well as Bacterial and Fungal Assemblages in the Rhizosphere of Balsam Poplar"

**Supplementary Table 1.** Balsam poplar genotyping on a Sequenom intraspecific panel. Trees were sampled at the Westwood site (W), the La Corne Mine site (C) and in a natural forest nearby of the La Corne Mine site (N). Unique genotypes were verified using a 40 SNP-array designed to reveal *P. balsamifera* intraspecific variations.

| **Sample** | **004** | **006** | **010** | **011** | **012** | **018** | **020** | **021** | **022** | **024** | **029** | **031** | **032** | **036** | **038** | **040** | **043** | **047** | **049** | **053** | **057** | **061** | **063** | **064** | **068** | **071** | **074** | **075** |
|---|---|---|---|---|---|---|---|---|---|---|---|---|---|---|---|---|---|---|---|---|---|---|---|---|---|---|---|---|
| **W01** | CC | AA | CT | TT | GG | AA | TT | CC | GT | AA | TT | CC | CT | AA | TT | AA | AA | AG | CC | CC | GT | GG | CC | CC | AA | TT | TT | GG |
| **W03** | TT | AA | CT | TT | GG | AA | TT | CC | TT | AT | TT | CC | CC | GG | TT | AG | AA | AA | CC | CT | GT | GG | TT | CT | AA | CT | TT | GG |
| **W05** | CT | AA | TT | TT | AG | AA | TT | CC | GT | TT | TT | CC | CT | AA | CT | AG | AG | GG | CC | CC | TT | CG | CC | CC | AA | CC | CC | GG |
| **W08** | TT | AA | CT | AT | GG | AA | TT | CC | GT | AA | CT | CC | CT | 0 | CT | AG | AA | GG | CC | CC | TT | GG | CT | CC | AA | CT | CT | GG |
| **W09** | CT | AA | CC | TT | AG | AA | CT | CC | GT | AT | TT | CC | CT | AG | TT | AG | AA | GG | CC | CT | GT | GG | CT | CC | AA | CT | TT | GG |
| **W10** | TT | AA | CT | TT | AG | AA | CT | CC | GT | AA | CT | CC | CT | AG | TT | AA | AA | GG | CC | CC | GG | GG | CT | CT | AA | TT | TT | GT |
| **W12** | TT | AA | CC | AA | AA | AA | CT | TT | GT | AT | CT | CC | CT | GG | TT | AA | AA | AG | AC | CC | GG | CG | CT | CC | AA | CT | CT | GG |
| **W13** | CT | AA | CT | TT | AG | AA | TT | CT | GG | AA | CC | CC | TT | AA | CT | AA | AG | GG | CC | CC | GT | GG | TT | CC | AA | TT | CT | GG |
| **N16** | CT | AA | CT | AT | GG | AA | CT | CT | TT | AA | CT | CC | CC | AA | CT | AG | AA | AG | CC | CC | GG | CG | CT | CC | AA | TT | CC | GG |
| **C17** | TT | AA | CT | AT | GG | AA | TT | TT | GT | AA | CT | AC | CC | AG | CT | AG | AA | GG | AC | CT | TT | GG | TT | CC | AA | CT | CT | GG |
| **C19** | CC | AC | CT | AT | GG | AA | TT | CT | GT | AA | CT | CC | CT | AA | TT | AA | AA | GG | CC | CC | GT | GG | TT | 0 | AA | CT | CC | GG |
| **C21** | TT | AA | CT | TT | AG | AA | TT | TT | GT | AT | CT | CC | CC | AG | CT | AG | AA | GG | CC | CC | TT | CG | CT | CC | AT | TT | CT | GG |
| **C23** | TT | AA | CT | TT | GG | AA | CT | CC | GT | AT | CC | CC | CC | AA | TT | AG | AA | AG | AC | CC | GG | CG | CT | CC | AA | TT | TT | GG |
| **C25** | TT | AA | CC | AT | GG | AA | TT | TT | GT | AT | CT | CC | CT | GG | CT | AG | AA | GG | CC | CT | GT | CG | CT | CT | AA | TT | TT | GG |
| **C27** | TT | AC | CT | TT | AG | AA | TT | CC | GT | AA | CT | CC | CC | AG | TT | GG | AA | GG | CC | CC | GG | GG | CT | CT | AA | TT | TT | GG |
| **C29** | TT | AA | CT | TT | GG | AA | TT | CT | GT | AA | CT | CC | CT | AG | CT | AA | AA | AG | CC | CT | GT | GG | TT | TT | AA | TT | CT | GT |
| **C30** | TT | AA | CT | TT | AG | AA | CT | CT | GG | AT | TT | CC | CC | AA | TT | AA | AA | GG | AC | CC | GT | GG | CT | CC | AA | TT | TT | GG |
| **N33** | TT | CC | CT | TT | GG | AA | CT | CC | GG | AA | CT | CC | CC | AA | TT | AA | AA | GG | CC | TT | GT | GG | CT | CT | AA | CT | CT | GG |
