## Supplementary Table 2 for "Plant Genotype Influences Physicochemical Properties of Substrate as well as Bacterial and Fungal Assemblages in the Rhizosphere of Balsam Poplar"

**Supplementary Table 2.** Pairwise comparison between all treatments on the field for bacterial and fungal taxa abundances and fungal functions. Two-way ANOVAs were used to discern how waste type, vegetation presence and their interaction influenced taxa relative abundance and fungal functions relative abundance. When a factor was revealed as a statistically significant predictor, a Tukey HSD post-hoc pairwise comparison test was performed between all treatments.

| **Bacteria** | | *Acetobacteraceae_g* | | *Acidimicrobiales_f_g* | | *Acidiphilium* | | *Acidobacteriaceae_g* | | *Actinomycetales_f_g* | | *[Pedosphaerales] auto67-4W_g* | |
|---|---|---|---|---|---|---|---|---|---|---|---|---|---|
| | Waste type | < 0.001 | | 0.072 | | < 0.001 | | < 0.001 | | 0.061 | | 0.325 | |
| | Vegetation presence | < 0.001 | | < 0.001 | | < 0.001 | | < 0.001 | | < 0.001 | | < 0.001 | |
| | Interaction | < 0.001 | | 0.047 | | < 0.001 | | < 0.001 | | 0.038 | | 0.524 | |
| **Pairwise comparison** | | | | | | | | | | | | | |
| Tailings | Unvegetated | 1.9% | B | 16.0% | A | 0.0% | B | 0.0% | C | 0.0% | C | 0.1% | B |
| | Vegetated | 1.4% | B | 0.1% | B | 0.0% | B | 2.4% | B | 1.2% | A | 1.0% | A |
| Waste rock | Unvegetated | 5.3% | A | 6.7% | A | 3.2% | A | 13.6% | A | 0.1% | BC | 0.0% | B |
| | Vegetated | 2.0% | B | 0.2% | B | 0.0% | B | 3.7% | B | 0.7% | AB | 0.8% | A |
| | p-value | < 0.001 | | < 0.001 | | < 0.001 | | < 0.001 | | < 0.001 | | < 0.001 | |
| **Fungi** | | *Acidea* | | *Agaricales_f_g* | | *Alternaria* | | *Amphinema* | | *Apiotrichum* | | *Ascomycota _o_c_f_g* | |
| | Waste type | 0.188 | | 0.352 | | 0.697 | | 0.068 | | 0.664 | | 0.038 | |
| | Vegetation presence | 0.294 | | 0.418 | | 0.029 | | < 0.001 | | 0.473 | | 0.665 | |
| | Interaction | 0.944 | | 0.418 | | 0.028 | | 0.003 | | 0.257 | | 0.364 | |
| **Pairwise comparison** | | | | | | | | | | | | | |
| Tailings | Unvegetated | 0.4% | A | 0.0% | A | 5.5% | A | 1.4% | A | 0.7% | A | 1.2% | A |
| | Vegetated | 0.0% | A | 2.5% | A | 0.1% | B | 0.0% | C | 0.1% | A | 1.6% | A |
| Waste rock | Unvegetated | 2.5% | A | 0.0% | A | 0.5% | AB | 0.2% | B | 0.1% | A | 0.3% | A |
| | Vegetated | 0.2% | A | 0.0% | A | 0.3% | B | 0.0% | C | 1.7% | A | 0.0% | A |
| | p-value | 0.392 | | 0.497 | | 0.004 | | < 0.001 | | 0.531 | | 0.078 | |
| **Fungal functions** | | Ectomycorrhizal | | Saprotroph | | Ericoid | | Plant pathogen | | Lichenized | | White rot | |
| | Waste type | 0.047 | | 0.169 | | 0.133 | | 0.337 | | 0.056 | | 0.178 | |
| | Vegetation presence | 0.008 | | < 0.001 | | 0.002 | | < 0.001 | | 0.958 | | 0.067 | |
| | Interaction | 0.358 | | 0.773 | | 0.340 | | 0.078 | | 0.398 | | 0.738 | |
| **Pairwise comparisons** | | | | | | | | | | | | | |
| Tailings | Unvegetated | 22.2% | B | 37.5% | A | 2.2% | B | 7.9% | A | 3.8% | A | 1.7% | A |
| | Vegetated | 68.2% | A | 8.9% | B | 7.3% | A | 0.4% | C | 2.2% | A | 0.1% | A |
| Waste rock | Unvegetated | 11.8% | B | 43.2% | A | 10.5% | AB | 4.2% | AB | 0.7% | A | 2.3% | A |
| | Vegetated | 36.4% | AB | 19.4% | AB | 35.6% | AB | 0.8% | BC | 0.9% | A | 0.0% | A |
| | p-value | < 0.001 | | < 0.001 | | < 0.001 | | < 0.001 | | 0.142 | | 0.092 | |

**Supplementary Table 2.** Pairwise comparison between all treatments on the field for bacterial and fungal taxa abundances and fungal functions. Two-way ANOVAs were used to discern how waste type, vegetation presence and their interaction influenced taxa relative abundance and fungal functions relative abundance. When a factor was revealed as a statistically significant predictor, a Tukey HSD post-hoc pairwise comparison test was performed between all treatments.

| **Bacteria** | | *Bradyrhizobium* | | *Burkholderia* | | *Chloroflexi C0119_o_f_g* | | *Candidatus Koribacter* | | *Candidatus Nitrososphaera* | | *Chitinophagaceae_g* | |
|---|---|---|---|---|---|---|---|---|---|---|---|---|---|
| | Waste type | < 0.001 | | 0.333 | | < 0.001 | | < 0.001 | | 0.018 | | < 0.001 | |
| | Vegetation presence | < 0.001 | | < 0.001 | | 0.591 | | 0.012 | | 0.024 | | < 0.001 | |
| | Interaction | 0.008 | | 0.182 | | < 0.001 | | 0.005 | | 0.018 | | < 0.001 | |
| **Pairwise comparison** | | | | | | | | | | | | | |
| Tailings | Unvegetated | 1.2% | C | 0.0% | B | 1.7% | A | 4.1% | A | 1.6% | A | 3.9% | B |
| | Vegetated | 7.8% | A | 3.8% | A | 0.0% | B | 0.4% | B | 0.0% | B | 6.7% | AB |
| Waste rock | Unvegetated | 0.0% | D | 0.0% | B | 0.0% | B | 0.0% | B | 0.0% | B | 0.0% | C |
| | Vegetated | 4.7% | B | 6.7% | A | 0.1% | B | 0.6% | B | 0.0% | B | 7.1% | A |
| | p-value | < 0.001 | | < 0.001 | | < 0.001 | | < 0.001 | | < 0.001 | | < 0.001 | |

| **Fungi** | | *Cenococcum* | | *Chaetosphaeriaceae_g* | | *Chaetothyriales_g* | | *Cistella* | | *Cladophialophora* | | *Cladosporium* | |
|---|---|---|---|---|---|---|---|---|---|---|---|---|---|
| | Waste type | < 0.001 | | 0.318 | | 0.887 | | 0.977 | | 0.112 | | 0.937 | |
| | Vegetation presence | 0.144 | | 0.001 | | 0.002 | | 0.009 | | 0.485 | | 0.003 | |
| | Interaction | 0.102 | | 0.065 | | 0.671 | | 0.027 | | 0.186 | | 0.092 | |
| **Pairwise comparison** | | | | | | | | | | | | | |
| Tailings | Unvegetated | 11.8% | AB | 1.2% | A | 1.2% | A | 4.3% | A | 0.1% | A | 11.7% | A |
| | Vegetated | 15.8% | A | 0.0% | B | 0.0% | B | 0.2% | B | 0.0% | A | 0.3% | B |
| Waste rock | Unvegetated | 3.5% | BC | 0.3% | AB | 0.9% | AB | 0.9% | AB | 13.1% | A | 3.8% | AB |
| | Vegetated | 0.0% | C | 0.0% | B | 0.0% | AB | 0.3% | B | 0.1% | A | 0.8% | B |
| | p-value | < 0.001 | | < 0.001 | | < 0.001 | | < 0.001 | | 0.139 | | < 0.001 | |

| **Fungal functions** | | Mycoparasite | | Arbuscular | |
|---|---|---|---|---|---|
| | Waste type | 0.170 | | 0.270 | |
| | Vegetation presence | 0.021 | | 0.208 | |
| | Interaction | 0.220 | | 0.208 | |
| **Pairwise comparisons** | | | | | |
| Tailings | Unvegetated | 0.2% | A | 0.2% | A |
| | Vegetated | 0.0% | A | 0.0% | B |
| Waste rock | Unvegetated | 1.5% | A | 0.0% | B |
| | Vegetated | 0.0% | A | 0.0% | AB |
| | p-value | 0.025 | | 0.003 | |

**Supplementary Table 2.** Pairwise comparison between all treatments on the field for bacterial and fungal taxa abundances and fungal functions. Two-way ANOVAs were used to discern how waste type, vegetation presence and their interaction influenced taxa relative abundance and fungal functions relative abundance. When a factor was revealed as a statistically significant predictor, a Tukey HSD post-hoc pairwise comparison test was performed between all treatments.

| **Bacteria** | | *Cytophagaceae_g* | | *[Chthoniobacteraceae] DA101* | | *Deinococcus* | | *Acidobacteria iii1-8 DS-18_f_g* | | *Thermoplasmata E2_f_g* | | *Alphaproteobacteria Ellin329_f_g* | |
|---|---|---|---|---|---|---|---|---|---|---|---|---|---|
| | Waste type | 0.146 | | < 0.001 | | < 0.001 | | < 0.001 | | 0.010 | | 0.085 | |
| | Vegetation presence | 0.015 | | 0.281 | | < 0.001 | | 0.002 | | 0.014 | | < 0.001 | |
| | Interaction | 0.034 | | < 0.001 | | < 0.001 | | < 0.001 | | 0.010 | | 0.002 | |
| **Pairwise comparison** | | | | | | | | | | | | | |
| Tailings | Unvegetated | 0.2% | AB | 6.0% | A | 1.2% | A | 3.3% | A | 1.7% | A | 0.0% | C |
| | Vegetated | 0.2% | AB | 0.6% | B | 0.0% | B | 0.2% | B | 0.0% | B | 2.8% | A |
| Waste rock | Unvegetated | 0.0% | B | 0.0% | B | 0.0% | B | 0.0% | B | 0.0% | B | 0.0% | C |
| | Vegetated | 0.8% | A | 0.1% | B | 0.0% | B | 0.1% | B | 0.0% | B | 1.7% | B |
| | p-value | 0.001 | | < 0.001 | | < 0.001 | | < 0.001 | | < 0.001 | | < 0.001 | |
| **Fungi** | | *Clavulinopsis* | | *Cortinarius* | | *Cryptococcus* | | *Exophiala* | | *Ganoderma* | | *Geomyces* | |
| | Waste type | < 0.001 | | 0.259 | | 0.302 | | 0.006 | | 0.267 | | 0.119 | |
| | Vegetation presence | 0.015 | | 0.002 | | 0.209 | | < 0.001 | | 0.019 | | 0.011 | |
| | Interaction | 0.016 | | 0.298 | | 0.098 | | < 0.001 | | 0.047 | | 0.681 | |
| **Pairwise comparison** | | | | | | | | | | | | | |
| Tailings | Unvegetated | 0.2% | B | 0.8% | AB | 0.9% | A | 1.4% | A | 1.1% | A | 1.8% | A |
| | Vegetated | 2.6% | A | 14.6% | A | 0.7% | A | 0.0% | B | 0.0% | B | 0.2% | A |
| Waste rock | Unvegetated | 0.0% | B | 0.3% | B | 0.4% | A | 0.0% | B | 0.2% | AB | 2.7% | A |
| | Vegetated | 0.0% | B | 31.2% | A | 2.2% | A | 0.1% | B | 0.0% | B | 0.0% | A |
| | p-value | < 0.001 | | < 0.001 | | 0.097 | | < 0.001 | | 0.003 | | 0.075 | |

**Supplementary Table 2.** Pairwise comparison between all treatments on the field for bacterial and fungal taxa abundances and fungal functions. Two-way ANOVAs were used to discern how waste type, vegetation presence and their interaction influenced taxa relative abundance and fungal functions relative abundance. When a factor was revealed as a statistically significant predictor, a Tukey HSD post-hoc pairwise comparison test was performed between all treatments.

| **Bacteria** | | *Betaproteobacteria Ellin6067_f_g* | | *Acidobacteria DA052 Ellin6513_f_g* | | *Flavobacterium* | | *Gaiellaceae_g* | | *Gammaproteobacteria _o_f_g* | | *Gemmata* | |
|---|---|---|---|---|---|---|---|---|---|---|---|---|---|
| | Waste type | < 0.001 | | < 0.001 | | 0.004 | | 0.100 | | < 0.001 | | 0.157 | |
| | Vegetation presence | 0.393 | | < 0.001 | | < 0.001 | | < 0.001 | | 0.065 | | 0.002 | |
| | Interaction | 0.004 | | < 0.001 | | 0.003 | | 0.224 | | < 0.001 | | 0.005 | |
| **Pairwise comparison** | | | | | | | | | | | | | |
| Tailings | Unvegetated | 1.2% | A | 0.0% | C | 0.0% | B | 1.2% | A | 0.3% | B | 2.0% | A |
| | Vegetated | 0.8% | AB | 7.2% | A | 0.4% | A | 0.8% | AB | 0.0% | B | 0.6% | B |
| Waste rock | Unvegetated | 0.0% | C | 0.0% | C | 0.0% | B | 0.0% | C | 14.2% | A | 0.0% | C |
| | Vegetated | 0.6% | B | 2.5% | B | 1.2% | A | 0.6% | B | 0.2% | B | 0.8% | B |
| | p-value | < 0.001 | | < 0.001 | | < 0.001 | | < 0.001 | | < 0.001 | | < 0.001 | |

| **Fungi** | | *Hebeloma* | | *Inocybe* | | *Knufia* | | *Lecanoromycetes_g* | | *Leotiomycetes_c_f_g* | | *Leptodontidium* | |
|---|---|---|---|---|---|---|---|---|---|---|---|---|---|
| | Waste type | 0.291 | | 0.288 | | 0.214 | | < 0.001 | | 0.355 | | 0.854 | |
| | Vegetation presence | 0.227 | | 0.827 | | 0.160 | | < 0.001 | | 0.592 | | 0.357 | |
| | Interaction | 0.227 | | 0.441 | | 0.530 | | < 0.001 | | 0.394 | | 0.037 | |
| **Pairwise comparison** | | | | | | | | | | | | | |
| Tailings | Unvegetated | 0.0% | A | 0.5% | A | 0.0% | A | 0.0% | B | 1.6% | A | 0.0% | A |
| | Vegetated | 0.0% | A | 8.4% | A | 0.1% | A | 1.3% | A | 1.5% | A | 0.2% | A |
| Waste rock | Unvegetated | 0.0% | A | 1.2% | A | 0.2% | A | 0.0% | B | 3.0% | A | 3.0% | A |
| | Vegetated | 4.8% | A | 0.2% | A | 1.2% | A | 0.0% | B | 0.1% | A | 0.0% | A |
| | p-value | 0.207 | | 0.582 | | 0.214 | | < 0.001 | | 0.565 | | 0.081 | |

**Supplementary Table 2.** Pairwise comparison between all treatments on the field for bacterial and fungal taxa abundances and fungal functions. Two-way ANOVAs were used to discern how waste type, vegetation presence and their interaction influenced taxa relative abundance and fungal functions relative abundance. When a factor was revealed as a statistically significant predictor, a Tukey HSD post-hoc pairwise comparison test was performed between all treatments.

| **Bacteria** | | *Gemmataceae_g* | | *Acidobacteria-6 iii1-15_f_g* | | *Isosphaeraceae_g* | | *AD3 JG37-AG-4 _o_f_g* | | *Kaistobacter* | | *Koribacteraceae_g* | | *Leptospirillum* | |
|---|---|---|---|---|---|---|---|---|---|---|---|---|---|---|---|
| | Waste type | < 0.001 | | 0.293 | | < 0.001 | | 0.604 | | < 0.001 | | 0.005 | | 0.360 | |
| | Vegetation presence | 0.285 | | < 0.001 | | < 0.001 | | 0.927 | | < 0.001 | | < 0.001 | | < 0.001 | |
| | Interaction | < 0.001 | | 0.044 | | 0.477 | | 0.001 | | < 0.001 | | < 0.001 | | < 0.001 | |
| **Pairwise comparison** | | | | | | | | | | | | | | | |
| Tailings | Unvegetated | 4.7% | AB | 1.1% | B | 0.2% | B | 0.0% | C | 4.8% | A | 0.0% | C | 0.0% | B |
| | Vegetated | 2.6% | B | 2.0% | A | 1.6% | A | 0.1% | B | 0.0% | C | 7.9% | A | 0.0% | B |
| Waste rock | Unvegetated | 7.1% | A | 0.0% | C | 1.9% | A | 8.4% | A | 0.0% | C | 0.0% | C | 7.2% | A |
| | Vegetated | 1.6% | B | 1.3% | AB | 0.5% | AB | 0.6% | BC | 1.2% | B | 2.7% | B | 0.0% | B |
| | p-value | < 0.001 | | < 0.001 | | < 0.001 | | < 0.001 | | < 0.001 | | < 0.001 | | < 0.001 | |

| **Fungi** | | *Lipomyces* | | *Meliniomyces* | | *Oidiodendron* | | *Parmelia* | | *Penicillium* | | *Pezoloma* | | *Phialocephala* | |
|---|---|---|---|---|---|---|---|---|---|---|---|---|---|---|---|
| | Waste type | 0.134 | | 0.114 | | 0.412 | | 0.022 | | 0.155 | | 0.258 | | 0.011 | |
| | Vegetation presence | 0.188 | | 0.006 | | 0.575 | | 0.035 | | 0.035 | | 0.101 | | 0.074 | |
| | Interaction | 0.188 | | 0.237 | | 0.788 | | 0.634 | | 0.706 | | 0.657 | | 0.694 | |
| **Pairwise comparison** | | | | | | | | | | | | | | | |
| Tailings | Unvegetated | 0.0% | A | 1.0% | B | 1.2% | A | 3.8% | A | 0.0% | B | 1.8% | A | 6.2% | AB |
| | Vegetated | 0.0% | A | 5.8% | A | 1.5% | A | 2.2% | AB | 0.7% | AB | 0.1% | A | 1.8% | A |
| Waste rock | Unvegetated | 1.2% | A | 7.0% | AB | 3.5% | A | 0.7% | AB | 0.5% | AB | 6.0% | A | 3.3% | AB |
| | Vegetated | 0.0% | A | 32.8% | AB | 2.7% | A | 0.0% | B | 1.6% | A | 1.2% | A | 0.4% | B |
| | p-value | 0.083 | | < 0.001 | | 0.766 | | 0.003 | | 0.007 | | 0.178 | | 0.003 | |

**Supplementary Table 2.** Pairwise comparison between all treatments on the field for bacterial and fungal taxa abundances and fungal functions. Two-way ANOVAs were used to discern how waste type, vegetation presence and their interaction influenced taxa relative abundance and fungal functions relative abundance. When a factor was revealed as a statistically significant predictor, a Tukey HSD post-hoc pairwise comparison test was performed between all treatments.

| **Bacteria** | | *[Methanomassiliicoccaceae]_g* | | *Methylobacterium* | | *Methylocystaceae_g* | | *Mycobacterium* | | *Oxalobacteraceae_g* | |
|---|---|---|---|---|---|---|---|---|---|---|---|
| | Waste type | 0.376 | | 0.039 | | < 0.001 | | 0.011 | | 0.003 | |
| | Vegetation presence | 0.007 | | < 0.001 | | < 0.001 | | < 0.001 | | < 0.001 | |
| | Interaction | 0.007 | | 0.105 | | < 0.001 | | 0.276 | | 0.489 | |
| **Pairwise comparison** | | | | | | | | | | | |
| Tailings | Unvegetated | 0.2% | B | 1.4% | A | 0.3% | B | 0.0% | B | 1.4% | A |
| | Vegetated | 0.0% | B | 0.0% | B | 1.3% | A | 0.6% | A | 0.4% | AB |
| Waste rock | Unvegetated | 2.8% | A | 0.0% | B | 0.0% | B | 0.1% | B | 0.0% | C |
| | Vegetated | 0.0% | B | 0.0% | B | 0.7% | AB | 1.4% | A | 0.3% | B |
| | p-value | < 0.001 | | < 0.001 | | < 0.001 | | < 0.001 | | < 0.001 | |
| **Fungi** | | *Piloderma* | | *Pleosporales_ fam_Incertae_sedis_g* | | *Pyrenopeziza* | | *Rozellomycota_ unidentified* | | *Sagenomella* | |
| | Waste type | < 0.001 | | 0.476 | | 0.133 | | 0.084 | | 0.375 | |
| | Vegetation presence | 0.214 | | 0.077 | | 0.061 | | < 0.001 | | 0.888 | |
| | Interaction | 0.002 | | 0.204 | | 0.464 | | 0.379 | | 0.435 | |
| **Pairwise comparison** | | | | | | | | | | | |
| Tailings | Unvegetated | 4.7% | B | 4.6% | A | 0.4% | A | 0.0% | A | 0.1% | A |
| | Vegetated | 19.7% | A | 0.1% | A | 0.0% | A | 3.0% | A | 0.1% | A |
| Waste rock | Unvegetated | 2.0% | BC | 0.7% | A | 1.8% | A | 0.0% | A | 1.2% | A |
| | Vegetated | 0.0% | C | 0.4% | A | 0.2% | A | 0.6% | A | 0.1% | A |
| | p-value | < 0.001 | | 0.099 | | 0.053 | | 0.057 | | 0.672 | |

**Supplementary Table 2.** Pairwise comparison between all treatments on the field for bacterial and fungal taxa abundances and fungal functions. Two-way ANOVAs were used to discern how waste type, vegetation presence and their interaction influenced taxa relative abundance and fungal functions relative abundance. When a factor was revealed as a statistically significant predictor, a Tukey HSD post-hoc pairwise comparison test was performed between all treatments.

| **Bacteria** | | *Planctomyces* | | *Proteobacteria _c_o_f_g* | | *Rhodoplanes* | | *Rhodospirillaceae_g* | | *Rubrivivax* | | *Sinobacteraceae_g* | |
|---|---|---|---|---|---|---|---|---|---|---|---|---|---|
| | Waste type | < 0.001 | | 0.397 | | 0.148 | | 0.021 | | 0.210 | | < 0.001 | |
| | Vegetation presence | 0.021 | | 0.008 | | < 0.001 | | < 0.001 | | < 0.001 | | 0.032 | |
| | Interaction | < 0.001 | | 0.006 | | 0.001 | | 0.087 | | 0.558 | | < 0.001 | |
| **Pairwise comparison** | | | | | | | | | | | | | |
| Tailings | Unvegetated | 0.9% | A | 0.1% | B | 0.4% | B | 0.2% | B | 1.2% | A | 0.3% | C |
| | Vegetated | 0.9% | A | 0.0% | B | 1.2% | A | 2.8% | A | 0.2% | B | 10.6% | A |
| Waste rock | Unvegetated | 0.0% | B | 4.0% | A | 0.0% | C | 0.0% | B | 0.0% | C | 5.3% | B |
| | Vegetated | 1.3% | A | 0.0% | B | 1.0% | AB | 2.3% | A | 0.1% | B | 4.1% | B |
| | p-value | < 0.001 | | < 0.001 | | < 0.001 | | < 0.001 | | < 0.001 | | < 0.001 | |

| **Fungi** | | *Setophoma* | | *Sistotrema* | | *Sordariales_g* | | *Talaromyces* | | *Teratosphaeriaceae_g* | | *Tetracladium* | |
|---|---|---|---|---|---|---|---|---|---|---|---|---|---|
| | Waste type | 0.217 | | 0.078 | | 0.145 | | 0.133 | | 0.025 | | 0.455 | |
| | Vegetation presence | 0.021 | | 0.047 | | 0.477 | | 0.188 | | 0.045 | | 0.004 | |
| | Interaction | 0.717 | | 0.047 | | 0.179 | | 0.188 | | 0.045 | | 0.231 | |
| **Pairwise comparison** | | | | | | | | | | | | | |
| Tailings | Unvegetated | 1.7% | A | 0.0% | A | 4.9% | A | 0.0% | A | 0.0% | B | 2.0% | A |
| | Vegetated | 0.1% | A | 0.0% | A | 0.4% | A | 0.0% | A | 0.0% | B | 0.1% | B |
| Waste rock | Unvegetated | 2.3% | A | 0.0% | A | 0.0% | A | 1.2% | A | 4.4% | A | 1.0% | AB |
| | Vegetated | 0.0% | A | 11.4% | A | 0.0% | A | 0.0% | A | 0.0% | B | 0.1% | B |
| | p-value | 0.025 | | 0.262 | | 0.162 | | 0.083 | | < 0.001 | | 0.001 | |

**Supplementary Table 2.** Pairwise comparison between all treatments on the field for bacterial and fungal taxa abundances and fungal functions. Two-way ANOVAs were used to discern how waste type, vegetation presence and their interaction influenced taxa relative abundance and fungal functions relative abundance. When a factor was revealed as a statistically significant predictor, a Tukey HSD post-hoc pairwise comparison test was performed between all treatments.

| **Bacteria** | | *Solibacterales_f_g* | | *Solirubrobacterales_f_g* | | *Sphingobacteriaceae_g* | | *Sphingobacteriales_f_g* | | *Sphingomonas* | |
|---|---|---|---|---|---|---|---|---|---|---|---|
| | Waste type | 0.321 | | 0.004 | | 0.349 | | 0.188 | | 0.882 | |
| | Vegetation presence | < 0.001 | | < 0.001 | | < 0.001 | | < 0.001 | | < 0.001 | |
| | Interaction | < 0.001 | | 0.427 | | < 0.001 | | 0.010 | | 0.008 | |
| **Pairwise comparison** | | | | | | | | | | | |
| Tailings | Unvegetated | 0.8% | B | 3.0% | A | 0.1% | C | 0.4% | BC | 0.3% | B |
| | Vegetated | 2.4% | A | 0.2% | B | 2.7% | B | 0.8% | AB | 0.5% | AB |
| Waste rock | Unvegetated | 0.0% | C | 0.3% | B | 0.0% | C | 0.0% | C | 0.0% | C |
| | Vegetated | 2.0% | A | 0.3% | B | 5.4% | A | 1.1% | A | 4.1% | A |
| | p-value | < 0.001 | | < 0.001 | | < 0.001 | | < 0.001 | | < 0.001 | |
| **Fungi** | | *Tomentella* | | *Tremella* | | *Tricholoma* | | *Umbelopsis* | | *Venturia* | |
| | Waste type | 0.448 | | 0.144 | | 0.032 | | 0.626 | | 0.048 | |
| | Vegetation presence | 0.285 | | 0.043 | | 0.054 | | 0.209 | | 0.051 | |
| | Interaction | 0.693 | | 0.248 | | 0.159 | | 0.391 | | 0.086 | |
| **Pairwise comparison** | | | | | | | | | | | |
| Tailings | Unvegetated | 0.0% | A | 0.2% | A | 2.3% | A | 1.3% | A | 0.2% | AB |
| | Vegetated | 0.0% | A | 0.0% | A | 8.9% | A | 1.4% | A | 0.2% | B |
| Waste rock | Unvegetated | 2.6% | A | 1.5% | A | 1.0% | A | 2.7% | A | 2.2% | A |
| | Vegetated | 0.0% | A | 0.0% | A | 0.0% | A | 0.5% | A | 0.5% | AB |
| | p-value | 0.559 | | 0.025 | | 0.004 | | 0.414 | | 0.003 | |

**Supplementary Table 2.** Pairwise comparison between all treatments on the field for bacterial and fungal taxa abundances and fungal functions. Two-way ANOVAs were used to discern how waste type, vegetation presence and their interaction influenced taxa relative abundance and fungal functions relative abundance. When a factor was revealed as a statistically significant predictor, a Tukey HSD post-hoc pairwise comparison test was performed between all treatments.

| **Bacteria** | | *Sulfobacillaceae _g* | | *Syntrophobacteraceae_g* | | *Thermogemmatisporaceae_g* | | *Xanthomonadaceae_g* | |
|---|---|---|---|---|---|---|---|---|---|
| | Waste type | < 0.001 | | < 0.001 | | 0.033 | | 0.018 | |
| | Vegetation presence | < 0.001 | | < 0.001 | | 0.031 | | 0.154 | |
| | Interaction | < 0.001 | | < 0.001 | | 0.008 | | 0.690 | |
| **Pairwise comparison** | | | | | | | | | |
| Tailings | Unvegetated | 0.0% | B | 0.1% | B | 0.0% | A | 0.0% | C |
| | Vegetated | 0.0% | B | 0.6% | A | 0.1% | A | 0.3% | B |
| Waste rock | Unvegetated | 6.3% | A | 1.3% | A | 1.1% | A | 0.1% | BC |
| | Vegetated | 0.0% | B | 0.5% | AB | 1.6% | A | 1.6% | A |
| | p-value | < 0.001 | | < 0.001 | | 0.020 | | < 0.001 | |
