## Supplementary Table 3 for "Plant Genotype Influences Physicochemical Properties of Substrate as well as Bacterial and Fungal Assemblages in the Rhizosphere of Balsam Poplar"

**Supplementary Table 3.** Spearman linear correlation analyses between bacterial and fungal taxa abundances and physicochemical properties of substrates in field samples. Weak correlations (>|0.3|) are highlighted in red; moderate correlations (>|0.5|) are highlighted in yellow; strong correlations (>|0.7|) are highlighted in green. CEC: Cation exchange capacity; BCSR: Base cation saturation ratio.

| **Bacteria** | **C total** | **N total** | **S total** | **pH** | **P** | **K** | **Ca** | **Mg** | **Mn** | **Fe** | **Na** | **CEC** | **BCSR** |
|---|---|---|---|---|---|---|---|---|---|---|---|---|---|
| *Acetobacteraceae_g* | -0.38 | -0.54 | 0.74 | -0.55 | 0.21 | -0.55 | 0.20 | -0.52 | -0.68 | 0.56 | -0.49 | 0.18 | 0.06 |
| *Acidimicrobiales_f_g* | -0.95 | -0.91 | 0.14 | 0.15 | -0.36 | -0.75 | -0.50 | -0.89 | -0.75 | -0.08 | -0.44 | -0.37 | -0.54 |
| *Acidiphilium* | -0.26 | -0.42 | 0.80 | -0.80 | 0.30 | -0.54 | 0.33 | -0.53 | -0.68 | 0.70 | -0.36 | 0.46 | 0.15 |
| *Acidobacteriaceae_g* | 0.22 | -0.01 | 0.89 | -0.88 | 0.56 | -0.30 | 0.67 | -0.19 | -0.46 | 0.77 | -0.35 | 0.58 | 0.51 |
| *Actinomycetales_f_g* | 0.83 | 0.82 | 0.15 | -0.38 | 0.48 | 0.54 | 0.37 | 0.57 | 0.44 | 0.32 | 0.37 | 0.24 | 0.38 |
| *[Pedosphaerales] auto67-4W_g* | 0.81 | 0.85 | -0.22 | 0.09 | 0.37 | 0.75 | 0.23 | 0.74 | 0.71 | -0.22 | 0.27 | 0.20 | 0.32 |
| *Bradyrhizobium* | 0.82 | 0.93 | -0.47 | 0.25 | 0.11 | 0.86 | 0.12 | 0.85 | 0.90 | -0.26 | 0.55 | 0.03 | 0.23 |
| *Burkholderia* | 0.80 | 0.79 | -0.02 | 0.00 | 0.60 | 0.73 | 0.25 | 0.69 | 0.56 | -0.20 | 0.09 | 0.20 | 0.27 |
| *Chloroflexi C0119_o_f_g* | -0.37 | -0.21 | -0.61 | 0.71 | -0.36 | 0.28 | -0.75 | 0.07 | 0.24 | -0.61 | 0.03 | -0.56 | -0.60 |
| *Candidatus Koribacter* | -0.26 | -0.04 | -0.67 | 0.62 | -0.48 | 0.22 | -0.71 | 0.01 | 0.31 | -0.52 | 0.20 | -0.62 | -0.57 |
| *Candidatus Nitrososphaera* | -0.69 | -0.53 | -0.53 | 0.73 | -0.70 | -0.29 | -0.74 | -0.40 | -0.17 | -0.54 | -0.05 | -0.73 | -0.63 |
| *Chitinophagaceae_g* | 0.58 | 0.64 | -0.54 | 0.45 | 0.29 | 0.76 | -0.01 | 0.69 | 0.71 | -0.49 | 0.19 | -0.05 | 0.10 |
| *Cytophagaceae_g* | 0.54 | 0.53 | -0.36 | 0.22 | 0.42 | 0.76 | -0.02 | 0.64 | 0.58 | -0.36 | 0.00 | 0.10 | 0.07 |
| *[Chthoniobacteraceae] DA101* | -0.14 | 0.09 | -0.82 | 0.80 | -0.62 | 0.31 | -0.64 | 0.21 | 0.49 | -0.65 | 0.38 | -0.64 | -0.46 |
| *Deinococcus* | -0.66 | -0.53 | -0.57 | 0.73 | -0.71 | -0.27 | -0.72 | -0.38 | -0.14 | -0.51 | -0.13 | -0.72 | -0.61 |
| *Acidobacteria iii1-8 DS-18_f_g* | -0.24 | -0.01 | -0.69 | 0.70 | -0.61 | 0.13 | -0.69 | 0.00 | 0.28 | -0.53 | 0.32 | -0.65 | -0.52 |
| *Thermoplasmata E2_f_g* | -0.69 | -0.53 | -0.53 | 0.73 | -0.70 | -0.29 | -0.74 | -0.40 | -0.17 | -0.54 | -0.05 | -0.73 | -0.63 |
| *Alphaproteobacteria Ellin329_f_g* | 0.89 | 0.96 | -0.19 | -0.05 | 0.30 | 0.75 | 0.41 | 0.81 | 0.77 | -0.02 | 0.56 | 0.29 | 0.47 |
| *Betaproteobacteria Ellin6067_f_g* | -0.08 | 0.11 | -0.96 | 0.84 | -0.52 | 0.41 | -0.55 | 0.31 | 0.53 | -0.67 | 0.39 | -0.42 | -0.41 |
| *Acidobacteria DA052 Ellin6513_f_g* | 0.90 | 0.97 | -0.20 | -0.08 | 0.31 | 0.78 | 0.29 | 0.80 | 0.79 | 0.00 | 0.54 | 0.20 | 0.36 |
| *Flavobacterium* | 0.71 | 0.74 | -0.23 | 0.11 | 0.53 | 0.73 | 0.22 | 0.71 | 0.63 | -0.35 | 0.20 | 0.20 | 0.28 |
| *Gaiellaceae_g* | -0.19 | 0.03 | -0.84 | 0.78 | -0.54 | 0.33 | -0.72 | 0.15 | 0.45 | -0.59 | 0.31 | -0.59 | -0.56 |
| *Gammaproteobacteria_o_f_g* | -0.27 | -0.40 | 0.67 | -0.59 | 0.18 | -0.64 | 0.34 | -0.54 | -0.70 | 0.57 | -0.31 | 0.29 | 0.14 |
| *Gemmata* | -0.16 | 0.03 | -0.76 | 0.86 | -0.43 | 0.38 | -0.62 | 0.24 | 0.48 | -0.73 | 0.17 | -0.47 | -0.44 |
| *Gemmataceae_g* | -0.54 | -0.63 | 0.27 | -0.36 | -0.30 | -0.76 | -0.02 | -0.71 | -0.64 | 0.48 | -0.24 | -0.08 | -0.13 |
| *Acidobacteria-6 iii1-15_f_g* | 0.54 | 0.70 | -0.74 | 0.46 | -0.11 | 0.75 | -0.07 | 0.75 | 0.85 | -0.39 | 0.68 | -0.02 | 0.04 |
| *Isosphaeraceae_g* | 0.40 | 0.26 | 0.51 | -0.80 | 0.14 | -0.08 | 0.61 | 0.03 | -0.05 | 0.81 | 0.05 | 0.44 | 0.51 |
| *AD3 JG37-AG-4_o_f_g* | 0.11 | -0.05 | 0.82 | -0.92 | 0.47 | -0.36 | 0.57 | -0.29 | -0.47 | 0.79 | -0.17 | 0.61 | 0.40 |
| *Kaistobacter* | -0.52 | -0.35 | -0.46 | 0.77 | -0.18 | 0.12 | -0.78 | -0.13 | 0.02 | -0.73 | -0.17 | -0.55 | -0.65 |
| *Koribacteraceae_g* | 0.87 | 0.94 | -0.16 | -0.12 | 0.33 | 0.72 | 0.35 | 0.75 | 0.70 | 0.02 | 0.54 | 0.24 | 0.41 |
| *Leptospirillum* | -0.26 | -0.46 | 0.78 | -0.78 | 0.22 | -0.63 | 0.33 | -0.58 | -0.74 | 0.68 | -0.44 | 0.31 | 0.12 |
| *Methanomassiliicoccaceae_g* | -0.40 | -0.61 | 0.48 | -0.51 | -0.01 | -0.62 | 0.05 | -0.63 | -0.71 | 0.54 | -0.57 | 0.04 | -0.14 |

| Bacteria | C total | N total | S total | pH | P | K | Ca | Mg | Mn | Fe | Na | CEC | BCSR |
|---|---|---|---|---|---|---|---|---|---|---|---|---|---|
| *Methylobacterium* | -0.64 | -0.48 | -0.63 | 0.77 | -0.53 | 0.00 | -0.78 | -0.18 | 0.02 | -0.63 | 0.01 | -0.49 | -0.68 |
| *Methylocystaceae_g* | 0.68 | 0.84 | -0.25 | 0.12 | 0.16 | 0.70 | -0.02 | 0.62 | 0.63 | -0.17 | 0.50 | -0.16 | 0.10 |
| *Mycobacterium* | 0.74 | 0.73 | 0.13 | -0.21 | 0.78 | 0.77 | 0.26 | 0.68 | 0.48 | -0.02 | 0.13 | 0.34 | 0.26 |
| *Oxalobacteraceae_g* | -0.27 | -0.07 | -0.74 | 0.88 | -0.46 | 0.24 | -0.65 | 0.11 | 0.37 | -0.85 | 0.14 | -0.55 | -0.48 |
| *Planctomyces* | 0.22 | 0.30 | -0.61 | 0.53 | 0.11 | 0.61 | -0.37 | 0.42 | 0.52 | -0.55 | 0.01 | -0.17 | -0.27 |
| *Proteobacteria_c_o_f_g* | -0.55 | -0.66 | 0.56 | -0.51 | -0.06 | -0.75 | 0.06 | -0.73 | -0.86 | 0.51 | -0.31 | 0.10 | -0.13 |
| *Rhodoplanes* | 0.62 | 0.74 | -0.52 | 0.34 | 0.08 | 0.81 | -0.10 | 0.74 | 0.87 | -0.42 | 0.48 | 0.02 | 0.03 |
| *Rhodospirillaceae_g* | 0.79 | 0.84 | -0.38 | 0.25 | 0.25 | 0.83 | 0.19 | 0.83 | 0.75 | -0.32 | 0.36 | 0.12 | 0.28 |
| *Rubrivivax* | -0.14 | 0.07 | -0.77 | 0.83 | -0.55 | 0.27 | -0.61 | 0.18 | 0.37 | -0.75 | 0.24 | -0.75 | -0.42 |
| *Sinobacteraceae_g* | 0.81 | 0.74 | 0.27 | -0.59 | 0.26 | 0.33 | 0.76 | 0.58 | 0.42 | 0.60 | 0.47 | 0.57 | 0.74 |
| *Solibacterales_f_g* | 0.68 | 0.79 | -0.34 | 0.22 | 0.32 | 0.81 | 0.00 | 0.73 | 0.72 | -0.27 | 0.42 | 0.00 | 0.12 |
| *Solirubrobacterales_f_g* | -0.62 | -0.52 | -0.28 | 0.52 | -0.36 | -0.29 | -0.64 | -0.42 | -0.28 | -0.34 | -0.17 | -0.52 | -0.64 |
| *Sphingobacteriaceae_g* | 0.65 | 0.71 | -0.22 | 0.18 | 0.54 | 0.86 | 0.06 | 0.74 | 0.60 | -0.34 | 0.15 | 0.09 | 0.14 |
| *Sphingobacteriales_f_g* | 0.50 | 0.61 | -0.40 | 0.38 | 0.38 | 0.87 | -0.11 | 0.74 | 0.65 | -0.47 | 0.31 | 0.05 | 0.01 |
| *Sphingomonas* | 0.49 | 0.54 | -0.45 | 0.41 | 0.42 | 0.76 | -0.12 | 0.62 | 0.63 | -0.50 | 0.05 | -0.02 | -0.01 |
| *Sulfobacillaceae_g* | -0.27 | -0.46 | 0.79 | -0.81 | 0.24 | -0.64 | 0.43 | -0.52 | -0.72 | 0.71 | -0.36 | 0.41 | 0.25 |
| *Syntrophobacteraceae_g* | 0.29 | 0.12 | 0.49 | -0.67 | 0.45 | -0.09 | 0.37 | -0.03 | -0.24 | 0.54 | -0.06 | 0.32 | 0.19 |
| *Thermogemmatisporaceae_g* | 0.37 | 0.23 | 0.47 | -0.52 | 0.71 | 0.10 | 0.14 | -0.03 | -0.17 | 0.40 | -0.34 | 0.16 | -0.01 |
| *Xanthomonadaceae_g* | 0.64 | 0.59 | 0.13 | -0.12 | 0.78 | 0.59 | 0.16 | 0.46 | 0.28 | -0.13 | -0.14 | 0.14 | 0.13 |

**Supplementary Table 3.** Spearman linear correlation analyses between bacterial and fungal taxa abundances and physicochemical properties of substrates in field samples. Weak correlations (>|0.3|) are highlighted in red; moderate correlations (>|0.5|) are highlighted in yellow; strong correlations (>|0.7|) are highlighted in green. CEC: Cation exchange capacity; BCSR: Base cation saturation ratio.

| **Fungi** | **C total** | **N total** | **S total** | **pH** | **P** | **K** | **Ca** | **Mg** | **Mn** | **Fe** | **Na** | **CEC** | **BCSR** |
|---|---|---|---|---|---|---|---|---|---|---|---|---|---|
| *Acidea* | -0.31 | -0.39 | 0.42 | -0.23 | 0.02 | -0.39 | -0.08 | -0.50 | -0.37 | 0.01 | -0.66 | -0.21 | -0.09 |
| *Agaricales_f_g* | 0.38 | 0.40 | -0.31 | 0.24 | -0.31 | 0.03 | 0.31 | 0.31 | 0.31 | -0.31 | 0.17 | -0.03 | 0.38 |
| *Alternaria* | -0.56 | -0.48 | -0.29 | 0.47 | -0.45 | -0.30 | -0.61 | -0.48 | -0.32 | -0.52 | -0.43 | -0.73 | -0.60 |
| *Amphinema* | -0.91 | -0.86 | 0.02 | 0.20 | -0.57 | -0.72 | -0.43 | -0.79 | -0.67 | -0.16 | -0.28 | -0.39 | -0.45 |
| *Apiotrichum* | 0.32 | 0.29 | 0.04 | -0.02 | 0.04 | 0.16 | 0.41 | 0.34 | 0.19 | 0.02 | -0.14 | 0.05 | 0.47 |
| *Ascomycota_o_c_f_g* | 0.28 | 0.48 | -0.63 | 0.40 | -0.45 | 0.30 | -0.01 | 0.41 | 0.40 | -0.23 | 0.71 | -0.16 | 0.09 |
| *Cenococcum* | 0.14 | 0.34 | -0.63 | 0.44 | -0.66 | 0.18 | -0.29 | 0.22 | 0.50 | -0.30 | 0.59 | -0.39 | -0.14 |
| *Chaetosphaeriaceae_g* | -0.63 | -0.53 | -0.19 | 0.40 | -0.68 | -0.34 | -0.59 | -0.47 | -0.29 | -0.37 | -0.15 | -0.57 | -0.53 |
| *Chaetothyriales_g* | -0.77 | -0.76 | 0.09 | 0.06 | -0.43 | -0.69 | -0.39 | -0.84 | -0.75 | -0.01 | -0.34 | -0.32 | -0.50 |
| *Cistella* | -0.68 | -0.59 | -0.18 | 0.34 | -0.62 | -0.39 | -0.59 | -0.57 | -0.34 | -0.28 | -0.07 | -0.35 | -0.58 |
| *Cladophialophora* | -0.42 | -0.59 | 0.65 | -0.60 | 0.23 | -0.75 | 0.14 | -0.80 | -0.70 | 0.49 | -0.59 | 0.18 | -0.04 |
| *Cladosporium* | -0.72 | -0.62 | -0.14 | 0.23 | -0.49 | -0.47 | -0.56 | -0.71 | -0.55 | -0.32 | -0.29 | -0.51 | -0.62 |
| *Clavulinopsis* | 0.54 | 0.70 | -0.67 | 0.33 | -0.44 | 0.43 | 0.07 | 0.63 | 0.65 | -0.21 | 0.80 | -0.13 | 0.22 |
| *Cortinarius* | 0.68 | 0.78 | -0.44 | 0.29 | 0.22 | 0.93 | -0.16 | 0.80 | 0.81 | -0.24 | 0.46 | -0.05 | -0.04 |
| *Cryptococcus* | 0.09 | 0.17 | -0.27 | 0.32 | 0.09 | 0.35 | -0.15 | 0.33 | 0.25 | -0.53 | 0.01 | -0.15 | -0.07 |
| *Exophiala* | -0.69 | -0.49 | -0.29 | 0.52 | -0.39 | -0.06 | -0.66 | -0.31 | -0.19 | -0.60 | -0.13 | -0.51 | -0.55 |
| *Ganoderma* | -0.27 | -0.20 | -0.46 | 0.65 | -0.52 | -0.08 | -0.59 | -0.17 | 0.01 | -0.56 | -0.14 | -0.68 | -0.57 |
| *Geomyces* | -0.55 | -0.52 | -0.09 | 0.25 | -0.56 | -0.57 | -0.37 | -0.64 | -0.50 | -0.14 | -0.24 | -0.49 | -0.41 |
| *Hebeloma* | 0.21 | 0.17 | 0.13 | 0.16 | 0.51 | 0.39 | 0.07 | 0.31 | 0.20 | -0.10 | -0.17 | 0.24 | 0.07 |
| *Inocybe* | 0.02 | 0.08 | -0.37 | 0.15 | -0.24 | 0.01 | 0.11 | 0.25 | 0.11 | -0.03 | 0.55 | 0.13 | 0.14 |
| *Knufia* | 0.34 | 0.24 | 0.03 | -0.13 | 0.35 | 0.39 | -0.18 | 0.21 | 0.11 | 0.01 | -0.13 | -0.07 | -0.24 |
| *Lecanoromycetes_g* | 0.70 | 0.79 | -0.54 | 0.13 | -0.20 | 0.47 | 0.20 | 0.66 | 0.76 | 0.01 | 0.72 | 0.07 | 0.31 |
| *Leotiomycetes_c_f_g* | 0.01 | 0.07 | -0.27 | 0.06 | -0.54 | -0.02 | -0.14 | -0.02 | 0.17 | 0.06 | 0.21 | -0.21 | -0.07 |
| *Leptodontidium* | 0.18 | 0.05 | 0.18 | -0.39 | -0.26 | -0.28 | 0.53 | 0.04 | 0.05 | 0.46 | 0.18 | 0.51 | 0.49 |
| *Lipomyces* | -0.10 | -0.38 | 0.46 | -0.52 | 0.26 | -0.41 | 0.13 | -0.28 | -0.45 | 0.52 | -0.28 | 0.23 | -0.06 |
| *Meliniomyces* | 0.59 | 0.51 | 0.13 | -0.17 | 0.39 | 0.22 | 0.52 | 0.48 | 0.34 | 0.06 | -0.10 | 0.08 | 0.57 |
| *Oidiodendron* | 0.29 | 0.31 | -0.18 | 0.17 | 0.16 | 0.25 | 0.07 | 0.42 | 0.41 | -0.16 | 0.19 | -0.04 | 0.16 |
| *Parmelia* | -0.24 | -0.02 | -0.53 | 0.36 | -0.58 | 0.03 | -0.49 | -0.09 | 0.17 | -0.21 | 0.45 | -0.42 | -0.38 |
| *Penicillium* | 0.67 | 0.51 | 0.19 | -0.22 | 0.59 | 0.56 | 0.26 | 0.59 | 0.42 | 0.18 | 0.07 | 0.42 | 0.22 |
| *Pezoloma* | -0.44 | -0.54 | 0.48 | -0.44 | 0.10 | -0.60 | 0.11 | -0.56 | -0.54 | 0.27 | -0.46 | -0.01 | 0.03 |
| *Phialocephala* | -0.43 | -0.41 | 0.16 | -0.12 | -0.44 | -0.67 | 0.02 | -0.59 | -0.41 | 0.10 | -0.09 | -0.09 | 0.02 |
| *Piloderma* | 0.27 | 0.45 | -0.68 | 0.36 | -0.58 | 0.23 | -0.23 | 0.27 | 0.51 | -0.22 | 0.64 | -0.34 | -0.11 |

| Fungi | C total | N total | S total | pH | P | K | Ca | Mg | Mn | Fe | Na | CEC | BCSR |
|---|---|---|---|---|---|---|---|---|---|---|---|---|---|
| *Pleosporales_fam_Incertae_sedis_g* | -0.54 | -0.56 | 0.03 | -0.09 | -0.16 | -0.33 | -0.39 | -0.54 | -0.64 | 0.05 | -0.27 | -0.26 | -0.52 |
| *Pyrenopeziza* | -0.47 | -0.58 | 0.62 | -0.45 | 0.02 | -0.59 | 0.04 | -0.68 | -0.61 | 0.24 | -0.63 | -0.02 | -0.03 |
| *Rozellomycota_g* | 0.81 | 0.88 | -0.39 | 0.18 | 0.23 | 0.75 | 0.19 | 0.78 | 0.84 | -0.22 | 0.42 | 0.18 | 0.29 |
| *Sagenomella* | -0.07 | -0.20 | 0.17 | -0.28 | 0.15 | -0.18 | 0.12 | -0.05 | -0.26 | 0.35 | -0.13 | 0.05 | 0.04 |
| *Setophoma* | -0.56 | -0.50 | -0.03 | 0.11 | -0.35 | -0.67 | -0.21 | -0.72 | -0.49 | -0.08 | -0.18 | -0.22 | -0.27 |
| *Sistotrema* | 0.10 | 0.11 | 0.13 | -0.11 | 0.56 | 0.45 | -0.20 | 0.14 | 0.09 | -0.16 | -0.23 | 0.02 | -0.20 |
| *Sordariales_g* | -0.29 | -0.11 | -0.56 | 0.43 | -0.28 | 0.09 | -0.51 | -0.08 | 0.05 | -0.42 | 0.12 | -0.45 | -0.47 |
| *Talaromyces* | -0.14 | -0.38 | 0.49 | -0.45 | -0.04 | -0.49 | 0.12 | -0.51 | -0.51 | 0.45 | -0.52 | 0.12 | -0.05 |
| *Teratosphaeriaceae_g* | -0.29 | -0.48 | 0.64 | -0.59 | 0.04 | -0.68 | 0.30 | -0.70 | -0.70 | 0.59 | -0.47 | 0.33 | 0.10 |
| *Tetracladium* | -0.69 | -0.59 | -0.09 | 0.30 | -0.48 | -0.52 | -0.47 | -0.70 | -0.42 | -0.21 | -0.19 | -0.33 | -0.49 |
| *Tomentella* | -0.16 | -0.25 | 0.20 | -0.08 | 0.22 | -0.11 | -0.44 | -0.38 | -0.27 | -0.17 | -0.65 | -0.51 | -0.51 |
| *Tremella* | -0.61 | -0.67 | 0.33 | -0.15 | -0.17 | -0.71 | -0.22 | -0.85 | -0.70 | 0.14 | -0.47 | -0.18 | -0.35 |
| *Tricholoma* | -0.03 | 0.13 | -0.39 | 0.14 | -0.42 | 0.06 | -0.25 | 0.01 | 0.21 | 0.07 | 0.54 | -0.20 | -0.17 |
| *Umbelopsis* | -0.26 | -0.24 | -0.06 | 0.08 | -0.05 | -0.41 | 0.05 | -0.24 | -0.12 | 0.08 | 0.12 | 0.10 | 0.02 |
| *Venturia* | -0.15 | -0.39 | 0.67 | -0.50 | 0.30 | -0.52 | 0.32 | -0.44 | -0.46 | 0.38 | -0.54 | 0.41 | 0.18 |
