## Supplementary Table 4 for "Plant Genotype Influences Physicochemical Properties of Substrate as well as Bacterial and Fungal Assemblages in the Rhizosphere of Balsam Poplar"

**Supplementary Table 4.** Factorial analysis of physicochemical properties of substrates in the greenhouse experiment. Two-way ANOVAs were used to discern how substrate type, genotype and their interaction influenced physicochemical properties of substrates. Additional analyses were performed for each substrate type separately to better assess the effect of genotype on physicochemical properties (see Table 2 and Supplementary Table 5).

| | C total | N total | S total | pH | P | K | Ca | Mg | Mn | Fe | Na | CEC | BCSR |
|---|---|---|---|---|---|---|---|---|---|---|---|---|---|
| **Substrate type** | < 0.001 | < 0.001 | < 0.001 | < 0.001 | < 0.001 | < 0.001 | < 0.001 | < 0.001 | < 0.001 | < 0.001 | < 0.001 | < 0.001 | < 0.001 |
| **Genotype** | 0.007 | < 0.001 | < 0.001 | 0.003 | < 0.001 | 0.386 | < 0.001 | < 0.001 | 0.076 | < 0.001 | < 0.001 | 0.002 | < 0.001 |
| **Interaction** | < 0.001 | < 0.001 | < 0.001 | 0.070 | 0.075 | < 0.001 | < 0.001 | < 0.001 | 0.005 | 0.121 | 0.001 | < 0.001 | 0.005 |
