## Supplementary Table 5 for "Plant Genotype Influences Physicochemical Properties of Substrate as well as Bacterial and Fungal Assemblages in the Rhizosphere of Balsam Poplar"

**Supplementary Table 5.** Other physicochemical properties of substrates after the greenhouse experiment. See Table 2 for the other parameters. CEC: Cation exchange capacity; BCSR: Base cation saturation ratio.

| | Mn (mmol/kg) | | | Na (mmol/kg) | | | CEC | | | BCSR | | |
|---|---|---|---|---|---|---|---|---|---|---|---|---|
| | **Control** | **Tailings** | **Waste rock** | **Control** | **Tailings** | **Waste rock** | **Control** | **Tailings** | **Waste rock** | **Control** | **Tailings** | **Waste rock** |
| **W08** | 0.84 b | 0.256 | 0.144 | 5.70 b | 0.59 b | 1.98 bc | 16.4 c | 4.0 b | 12.5 | 15.7 abcd | 55.0 d | 38.9 cd |
| **W09** | 2.79 a | 0.432 | 0.177 | 16.10 a | 2.47 a | 3.90 a | 43.9 a | 7.6 a | 16.0 | 18.0 abc | 57.5 bcd | 48.6 ab |
| **W10** | 2.25 ab | 0.495 | 0.214 | 14.30 ab | 2.54 a | 2.90 abc | 42.4 a | 8.8 a | 15.4 | 9.0 bcd | 61.6 bcd | 45.9 abc |
| **W13** | 2.45 a | 0.514 | 0.185 | 12.70 ab | 2.50 a | 4.00 ab | 38.7 abc | 8.3 a | 15.7 | 25.7 a | 62.8 abcd | 53.3 a |
| **N16** | 1.94 ab | 0.605 | 0.116 | 11.30 ab | 2.42 ab | 2.28 abc | 34.3 abc | 9.5 a | 13.7 | 23.5 a | 71.1 a | 45.7 abc |
| **C21** | 1.73 ab | 0.702 | 0.163 | 12.80 ab | 2.19 ab | 1.59 c | 39.1 abc | 9.2 a | 11.4 | 8.7 cd | 64.6 ab | 35.4 d |
| **C23** | 1.77 ab | 0.456 | 0.241 | 9.20 b | 1.57 ab | 3.66 abc | 31.2 bc | 8.2 a | 15.4 | 20.0 ab | 59.9 bcd | 50.6 a |
| **C25** | 2.23 ab | 0.623 | 0.162 | 11.80 ab | 2.91 a | 2.64 abc | 41.8 a | 8.8 a | 13.5 | 6.3 d | 63.9 abc | 42.0 bcd |
| **C29** | 1.98 ab | 0.597 | 0.185 | 11.50 ab | 1.99 ab | 3.13 abc | 35.5 abc | 9.3 a | 16.4 | 18.0 abc | 60.9 bcd | 51.9 a |
| **N33** | 2.33 ab | 0.563 | 0.202 | 16.50 a | 2.20 ab | 2.44 abc | 41.2 ab | 8.6 a | 13.6 | 8.0 cd | 56.5 cd | 46.6 ab |
| **p-value** | 0.023 | 0.086 | 0.326 | 0.007 | 0.007 | 0.005 | < 0.001 | < 0.001 | 0.021 | < 0.001 | 0.007 | < 0.001 |
