## Supplementary Table 6 for "Plant Genotype Influences Physicochemical Properties of Substrate as well as Bacterial and Fungal Assemblages in the Rhizosphere of Balsam Poplar"

**Supplementary Table 6.** Spearman linear correlation analyses between physicochemical properties of substrates and tree growth measurements in the greenhouse experiment. Weak correlations (>|0.3|) are highlighted in red; moderate correlations (>|0.5|) are highlighted in yellow; strong correlations (>|0.7|) are highlighted in green. CEC: Cation exchange capacity; BCSR: Base cation saturation ratio.

| | | Chlorophyll content | | | | |
|---|---|---|---|---|---|---|
| | **Growth** | **Season 1** | **Season 2** | **Shoot diameter** | **Biomass** | **Blooming** |
| **C total** | 0.04 | 0.07 | -0.01 | 0.14 | 0.22 | -0.04 |
| **N total** | 0.00 | 0.05 | 0.05 | 0.17 | 0.32 | -0.04 |
| **S total** | -0.17 | -0.03 | 0.21 | -0.07 | 0.00 | -0.06 |
| **pH** | 0.13 | -0.18 | -0.29 | -0.05 | -0.18 | -0.06 |
| **P** | 0.00 | 0.05 | 0.00 | 0.05 | 0.13 | 0.03 |
| **K** | 0.19 | 0.08 | -0.13 | 0.08 | 0.10 | -0.06 |
| **Ca** | 0.05 | 0.05 | -0.06 | 0.03 | 0.24 | -0.04 |
| **Mg** | 0.18 | 0.11 | -0.29 | 0.09 | 0.15 | -0.11 |
| **Mn** | 0.14 | 0.12 | -0.24 | 0.23 | 0.19 | 0.01 |
| **Fe** | -0.15 | 0.00 | 0.27 | -0.23 | -0.01 | 0.13 |
| **Na** | 0.01 | 0.10 | -0.08 | 0.16 | 0.39 | -0.07 |
| **CEC** | 0.01 | 0.07 | 0.01 | 0.06 | 0.28 | -0.05 |
| **BCSR** | 0.16 | 0.08 | -0.27 | 0.15 | 0.09 | -0.06 |
