## Supplementary Table 7 for "Plant Genotype Influences Physicochemical Properties of Substrate as well as Bacterial and Fungal Assemblages in the Rhizosphere of Balsam Poplar"

**Supplementary Table 7.** Pairwise comparison between genotypes in each substrate type in the greenhouse experiment for fungal functions relative abundance. Two-way ANOVAs were used to discern how substrate type, genotype and their interaction influenced fungal functions relative abundance. Additional analyses were performed for each substrate type separately and between substrate types to better assess the indivudial effects of genotype and substrate type.

| **Anova** | | Ectomycorrhizae | Saprotroph | Plant pathogen | Ericoid mycorrhizae | Brown rot |
|---|---|---|---|---|---|---|
| | Genotype | 0.718 | 0.896 | 0.171 | 0.144 | 0.071 |
| | Substrate type | < 0.001 | < 0.001 | 0.029 | < 0.001 | 0.310 |
| | Interaction | 0.093 | 0.450 | 0.594 | 0.099 | 0.558 |
| **Pairwise comparison between substrate types** | | | | | | |
| | Control | 41.3% A | 24.3% A | 6.7% A | 0.1% B | 0.4% A |
| | Tailings | 28.4% B | 19.1% B | 4.7% B | 0.1% B | 0.3% A |
| | Waste rock | 49.6% A | 23.6% A | 5.0% AB | 2.8% A | 0.5% A |
| | p-value | < 0.001 | < 0.001 | 0.019 | 0.001 | 0.245 |
| **Pairwise comparison by substrate type** | | | | | | |
| Control | W08 | 36.8% A | 19.4% A | 5.4% A | 0.2% A | 0.5% A |
| | W09 | 37.8% A | 23.3% A | 7.2% A | 0.2% A | 0.4% A |
| | W10 | 37.0% A | 26.0% A | 7.0% A | 0.3% A | 0.2% A |
| | W13 | 36.8% A | 26.2% A | 8.4% A | 0.2% A | 0.1% A |
| | N16 | 55.9% A | 24.6% A | 6.7% A | 0.1% A | 0.0% A |
| | C21 | 49.8% A | 24.1% A | 3.3% A | 0.0% A | 0.5% A |
| | C23 | 33.9% A | 23.9% A | 7.4% A | 0.2% A | 0.5% A |
| | C25 | 35.3% A | 27.9% A | 7.8% A | 0.1% A | 0.4% A |
| | C29 | 58.7% A | 19.8% A | 4.7% A | 0.1% A | 0.0% A |
| | N33 | 38.5% A | 27.5% A | 8.1% A | 0.1% A | 0.9% A |
| | p-value | 0.390 | 0.686 | 0.207 | 0.362 | 0.063 |
| Tailngs | W08 | 24.0% A | 22.3% A | 3.5% A | 0.1% A | 0.7% A |
| | W09 | 34.8% A | 17.2% A | 6.7% A | 0.0% A | 0.1% A |
| | W10 | 8.4% A | 21.9% A | 7.3% A | 0.7% A | 0.2% A |
| | W13 | 38.9% A | 17.0% A | 4.5% A | 0.0% A | 0.2% A |
| | N16 | 26.8% A | 17.3% A | 9.9% A | 0.1% A | 0.1% A |
| | C21 | 22.0% A | 26.2% A | 3.3% A | 0.0% A | 0.3% A |
| | C23 | 32.2% A | 17.6% A | 4.6% A | 0.0% A | 0.2% A |
| | C25 | 28.7% A | 14.5% A | 3.4% A | 0.0% A | 0.1% A |
| | C29 | 40.6% A | 19.5% A | 2.6% A | 0.0% A | 0.2% A |
| | N33 | 25.6% A | 17.2% A | 2.9% A | 0.0% A | 0.7% A |
| | p-value | 0.148 | 0.229 | 0.516 | 0.414 | 0.730 |
| Waste rock | W08 | 53.3% A | 18.8% A | 4.5% A | 2.7% AB | 0.5% A |
| | W09 | 45.1% A | 27.1% A | 6.8% A | 0.8% B | 0.3% A |
| | W10 | 63.4% A | 21.8% A | 2.4% A | 3.4% AB | 0.1% A |
| | W13 | 52.7% A | 25.7% A | 6.6% A | 0.3% AB | 0.9% A |
| | N16 | 57.5% A | 25.3% A | 2.4% A | 2.7% AB | 0.2% A |
| | C21 | 57.5% A | 21.1% A | 1.3% A | 2.6% AB | 0.2% A |
| | C23 | 40.4% A | 27.9% A | 5.9% A | 2.5% AB | 0.8% A |
| | C25 | 55.3% A | 23.5% A | 6.0% A | 0.5% B | 1.1% A |
| | C29 | 35.2% A | 21.1% A | 8.7% A | 0.5% B | 0.7% A |
| | N33 | 41.7% A | 24.8% A | 4.7% A | 9.7% A | 0.4% A |
| | p-value | 0.380 | 0.907 | 0.484 | 0.005 | 0.348 |

**Supplementary Table 7.** Pairwise comparison between genotypes in each substrate type in the greenhouse experiment for fungal functions relative abundance. Two-way ANOVAs were used to discern how substrate type, genotype and their interaction influenced fungal functions relative abundance. Additional analyses were performed for each substrate type separately and between substrate types to better assess the indivudial effects of genotype and substrate type.

| **Anova** | | White rot | | Lichenized mycorrhizae | | Arbuscular mycorrhizae | |
|---|---|---|---|---|---|---|---|
| | Genotype | 0.472 | | 0.063 | | 0.696 | |
| | Substrate type | < 0.001 | | 0.997 | | 0.384 | |
| | Interaction | 0.885 | | 0.270 | | 0.931 | |
| **Pairwise comparison between substrate types** | | | | | | | |
| | Control | 0.2% | A | 0.0% | A | 0.0% | A |
| | Tailings | 0.0% | B | 0.0% | A | 0.0% | A |
| | Waste rock | 0.1% | B | 0.0% | A | 0.0% | A |
| | p-value | < 0.001 | | 0.998 | | 0.281 | |
| **Pairwise comparison by substrate type** | | | | | | | |
| Control | W08 | 0.2% | A | 0.0% | A | 0.0% | A |
| | W09 | 0.3% | A | 0.0% | A | 0.0% | A |
| | W10 | 0.2% | A | 0.0% | A | 0.0% | A |
| | W13 | 0.1% | A | 0.0% | A | 0.0% | A |
| | N16 | 0.2% | A | 0.0% | A | 0.0% | A |
| | C21 | 0.1% | A | 0.0% | A | 0.0% | A |
| | C23 | 0.2% | A | 0.0% | A | 0.0% | A |
| | C25 | 0.2% | A | 0.0% | A | 0.0% | A |
| | C29 | 0.1% | A | 0.0% | A | 0.0% | A |
| | N33 | 0.2% | A | 0.0% | A | 0.0% | A |
| | p-value | 0.717 | | 0.622 | | 0.999 | |
| Tailngs | W08 | 0.0% | A | 0.0% | A | 0.0% | A |
| | W09 | 0.0% | A | 0.0% | A | 0.0% | A |
| | W10 | 0.1% | A | 0.0% | A | 0.0% | A |
| | W13 | 0.1% | A | 0.0% | A | 0.0% | A |
| | N16 | 0.0% | A | 0.0% | A | 0.0% | A |
| | C21 | 0.0% | A | 0.0% | A | 0.0% | A |
| | C23 | 0.1% | A | 0.0% | A | 0.0% | A |
| | C25 | 0.0% | A | 0.0% | A | 0.0% | A |
| | C29 | 0.0% | A | 0.0% | A | 0.0% | A |
| | N33 | 0.1% | A | 0.0% | A | 0.0% | A |
| | p-value | 0.668 | | 0.049 | | 0.781 | |
| Waste rock | W08 | 0.1% | A | 0.0% | A | 0.0% | A |
| | W09 | 0.2% | A | 0.0% | A | 0.0% | A |
| | W10 | 0.1% | A | 0.0% | A | 0.0% | A |
| | W13 | 0.1% | A | 0.0% | A | 0.0% | A |
| | N16 | 0.1% | A | 0.0% | A | 0.0% | A |
| | C21 | 0.0% | A | 0.0% | A | 0.0% | A |
| | C23 | 0.1% | A | 0.0% | A | 0.0% | A |
| | C25 | 0.0% | A | 0.0% | A | 0.0% | A |
| | C29 | 0.1% | A | 0.0% | A | 0.0% | A |
| | N33 | 0.1% | A | 0.0% | A | 0.0% | A |
| | p-value | 0.645 | | 0.150 | | 0.999 | |
