## Supplementary Table 8 for "Plant Genotype Influences Physicochemical Properties of Substrate as well as Bacterial and Fungal Assemblages in the Rhizosphere of Balsam Poplar"

**Supplementary Table 8.** Pairwise comparison between genotypes in each substrate type in the greenhouse experiment for bacterial and fungal taxa relative abundance. Two-way ANOVAs were used to discern how substrate type, genotype and their interaction influenced taxa relative abundance. Additional analyses were performed for each substrate type separately and between substrate types to better assess the indivudial effects of genotype and substrate type.

| **Bacteria** | | *Anaerolineae SBR1031 A4b_g* | *Acidobacteriaceae_g* | *Bradyrhizobium* | *Caulobacteraceae_g* |
|---|---|---|---|---|---|
| | Genotype | 0.004 | 0.694 | 0.013 | 0.168 |
| | Substrate type | < 0.001 | < 0.001 | < 0.001 | < 0.001 |
| | Interaction | 0.307 | 0.707 | 0.011 | 0.124 |
| **Pairwise comparison between substrate types** | | | | | |
| | Control | 0.8% B | 0.1% B | 0.4% C | 0.5% B |
| | Tailings | 1.4% A | 0.0% C | 0.9% B | 0.5% B |
| | Waste rock | 0.7% B | 1.9% A | 1.2% A | 1.8% A |
| | p-value | < 0.001 | < 0.001 | < 0.001 | < 0.001 |
| **Pairwise comparison by substrate type** | | | | | |
| Control | W08 | 0.9% A | 0.2% A | 0.4% AB | 0.5% A |
| | W09 | 0.6% A | 0.1% A | 0.5% AB | 0.5% A |
| | W10 | 0.6% A | 0.1% A | 0.3% B | 0.5% A |
| | W13 | 0.7% A | 0.1% A | 0.4% AB | 0.4% A |
| | N16 | 0.9% A | 0.0% A | 0.3% AB | 0.6% A |
| | C21 | 0.7% A | 0.1% A | 0.7% A | 0.8% A |
| | C23 | 1.1% A | 0.2% A | 0.4% AB | 0.5% A |
| | C25 | 0.7% A | 0.1% A | 0.4% AB | 0.6% A |
| | C29 | 0.7% A | 0.1% A | 0.5% AB | 0.6% A |
| | N33 | 0.8% A | 0.1% A | 0.4% AB | 0.5% A |
| | p-value | 0.321 | 0.557 | 0.011 | 0.503 |
| Tailngs | W08 | 1.4% A | 0.0% A | 0.6% A | 0.4% A |
| | W09 | 1.6% A | 0.0% A | 1.3% A | 0.6% A |
| | W10 | 1.3% A | 0.1% A | 0.9% A | 0.5% A |
| | W13 | 1.4% A | 0.0% A | 0.9% A | 0.5% A |
| | N16 | 1.3% A | 0.0% A | 0.6% A | 0.3% A |
| | C21 | 1.0% A | 0.0% A | 0.8% A | 0.4% A |
| | C23 | 1.5% A | 0.0% A | 1.5% A | 0.6% A |
| | C25 | 1.3% A | 0.0% A | 0.5% A | 0.8% A |
| | C29 | 1.5% A | 0.0% A | 1.0% A | 0.5% A |
| | N33 | 1.5% A | 0.1% A | 1.0% A | 0.5% A |
| | p-value | 0.904 | 0.667 | 0.018 | 0.047 |
| Waste rock | W08 | 0.4% B | 2.5% A | 1.5% A | 1.7% A |
| | W09 | 0.9% AB | 1.4% A | 1.3% A | 1.5% A |
| | W10 | 0.5% AB | 1.6% A | 0.9% A | 2.2% A |
| | W13 | 0.6% AB | 1.5% A | 1.3% A | 1.4% A |
| | N16 | 0.8% AB | 2.2% A | 1.5% A | 2.4% A |
| | C21 | 0.3% B | 2.5% A | 1.3% A | 2.4% A |
| | C23 | 1.4% A | 1.3% A | 1.2% A | 1.3% A |
| | C25 | 0.4% B | 2.6% A | 0.8% A | 1.8% A |
| | C29 | 1.1% AB | 1.3% A | 1.0% A | 1.4% A |
| | N33 | 0.5% B | 1.8% A | 1.1% A | 1.7% A |
| | p-value | < 0.001 | 0.711 | 0.123 | 0.188 |

**Supplementary Table 8.** Pairwise comparison between genotypes in each substrate type in the greenhouse experiment for bacterial and fungal taxa relative abundance. Two-way ANOVAs were used to discern how substrate type, genotype and their interaction influenced taxa relative abundance. Additional analyses were performed for each substrate type separately and between substrate types to better assess the indivudial effects of genotype and substrate type.

| **Bacteria** | | *Chitinophagaceae_g* | *Acidimicrobiales EB1017_g* | *Alphaproteobacteria Ellin329_f_g* | *Betaproteobacteria Ellin6067_f_g* |
|---|---|---|---|---|---|
| | Genotype | 0.011 | 0.690 | 0.022 | 0.203 |
| | Substrate type | < 0.001 | < 0.001 | < 0.001 | < 0.001 |
| | Interaction | 0.261 | 0.245 | 0.451 | 0.026 |
| **Pairwise comparison between substrate types** | | | | | |
| | Control | 22.4% A | 0.9% B | 6.1% B | 2.0% A |
| | Tailings | 11.9% B | 1.1% A | 3.6% C | 1.6% B |
| | Waste rock | 8.2% C | 0.4% C | 8.7% A | 2.1% A |
| | p-value | < 0.001 | < 0.001 | < 0.001 | < 0.001 |
| **Pairwise comparison by substrate type** | | | | | |
| Control | W08 | 23.9% A | 0.8% A | 5.2% A | 2.3% A |
| | W09 | 25.9% A | 0.9% A | 6.2% A | 2.2% A |
| | W10 | 20.8% A | 0.6% A | 7.2% A | 2.5% A |
| | W13 | 26.0% A | 0.6% A | 4.6% A | 1.4% A |
| | N16 | 25.2% A | 1.1% A | 6.3% A | 1.7% A |
| | C21 | 17.3% A | 0.9% A | 7.0% A | 2.1% A |
| | C23 | 19.0% A | 0.8% A | 6.3% A | 2.3% A |
| | C25 | 24.1% A | 0.9% A | 6.3% A | 2.3% A |
| | C29 | 18.2% A | 0.9% A | 5.6% A | 1.8% A |
| | N33 | 22.7% A | 1.0% A | 6.8% A | 1.8% A |
| | p-value | 0.069 | 0.710 | 0.823 | 0.217 |
| Tailngs | W08 | 13.9% A | 1.1% A | 2.7% AB | 1.5% A |
| | W09 | 13.4% A | 1.3% A | 2.9% AB | 1.9% A |
| | W10 | 11.1% A | 1.4% A | 4.1% AB | 1.4% A |
| | W13 | 12.5% A | 1.5% A | 3.8% AB | 1.3% A |
| | N16 | 8.9% A | 1.3% A | 3.3% AB | 2.1% A |
| | C21 | 10.9% A | 0.7% A | 4.6% AB | 2.0% A |
| | C23 | 11.2% A | 1.1% A | 2.3% B | 1.3% A |
| | C25 | 14.4% A | 0.9% A | 5.3% A | 1.5% A |
| | C29 | 9.0% A | 0.9% A | 2.6% AB | 1.6% A |
| | N33 | 12.6% A | 1.1% A | 4.4% AB | 1.8% A |
| | p-value | 0.350 | 0.128 | 0.002 | 0.030 |
| Waste rock | W08 | 9.3% A | 0.5% A | 11.2% A | 1.8% A |
| | W09 | 11.1% A | 0.5% A | 8.1% A | 2.3% A |
| | W10 | 6.7% A | 0.5% A | 9.4% A | 1.7% A |
| | W13 | 9.4% A | 0.4% A | 8.5% A | 2.2% A |
| | N16 | 7.0% A | 0.3% A | 7.6% A | 2.6% A |
| | C21 | 3.2% A | 0.2% A | 7.6% A | 1.5% A |
| | C23 | 8.6% A | 0.4% A | 9.4% A | 2.6% A |
| | C25 | 7.3% A | 0.2% A | 7.5% A | 2.3% A |
| | C29 | 11.9% A | 0.5% A | 8.1% A | 2.0% A |
| | N33 | 6.2% A | 0.2% A | 8.8% A | 1.8% A |
| | p-value | 0.051 | 0.527 | 0.719 | 0.155 |

**Supplementary Table 8.** Pairwise comparison between genotypes in each substrate type in the greenhouse experiment for bacterial and fungal taxa relative abundance. Two-way ANOVAs were used to discern how substrate type, genotype and their interaction influenced taxa relative abundance. Additional analyses were performed for each substrate type separately and between substrate types to better assess the indivudial effects of genotype and substrate type.

| **Bacteria** | | *Frankiaceae_g* | *Gaiellaceae_g* | *Gemmataceae_g* | *Geobacter* | *Acidobacteria-6 iii1_15_f_g* |
|---|---|---|---|---|---|---|
| | Genotype | 0.008 | 0.339 | < 0.001 | < 0.001 | 0.608 |
| | Substrate type | < 0.001 | < 0.001 | < 0.001 | < 0.001 | < 0.001 |
| | Interaction | 0.081 | 0.001 | 0.002 | 0.006 | 0.057 |
| **Pairwise comparison between substrate types** | | | | | | |
| | Control | 1.4% A | 0.8% B | 0.8% B | 0.2% B | 0.9% B |
| | Tailings | 0.8% C | 0.6% C | 3.7% A | 5.6% A | 2.2% A |
| | Waste rock | 1.0% B | 1.2% A | 1.5% C | 0.2% B | 0.6% C |
| | p-value | < 0.001 | < 0.001 | < 0.001 | < 0.001 | < 0.001 |
| **Pairwise comparison by substrate type** | | | | | | |
| Control | W08 | 1.4% A | 1.0% A | 1.4% A | 0.0% A | 1.0% A |
| | W09 | 1.2% A | 0.7% A | 0.7% A | 0,0% A | 0.8% A |
| | W10 | 1.7% A | 0.7% A | 0.6% A | 0.1% A | 1.1% A |
| | W13 | 1.4% A | 0.6% A | 0.9% A | 0.0% A | 1.1% A |
| | N16 | 1.3% A | 0.6% A | 1.1% A | 0,0% A | 0.9% A |
| | C21 | 1.6% A | 1.0% A | 0.6% A | 0,0% A | 0.9% A |
| | C23 | 1.4% A | 0.9% A | 0.8% A | 0.6% A | 0.7% A |
| | C25 | 1.4% A | 0.9% A | 0.8% A | 0.7% A | 0.9% A |
| | C29 | 1.4% A | 0.6% A | 0.6% A | 0.1% A | 0.9% A |
| | N33 | 1.4% A | 0.9% A | 0.8% A | 0.2% A | 1.0% A |
| | p-value | 0.844 | 0.495 | 0.205 | 0.669 | 0.805 |
| Tailngs | W08 | 0.9% AB | 0.6% A | 4.8% A | 4.5% ABC | 2.5% AB |
| | W09 | 0.8% AB | 0.8% A | 2.7% CD | 1.9% ABC | 1.8% AB |
| | W10 | 0.7% AB | 0.7% A | 4.2% ABC | 10.8% A | 2.2% AB |
| | W13 | 0.5% B | 0.6% A | 4.3% AB | 2.6% ABC | 3.0% A |
| | N16 | 1.5% A | 0.3% A | 5.0% A | 0.0% C | 2.8% AB |
| | C21 | 0.8% AB | 0.4% A | 3.4% ABCD | 6.7% ABC | 2.4% AB |
| | C23 | 0.4% B | 0.5% A | 3.1% BCD | 10.8% A | 1.4% B |
| | C25 | 1.1% AB | 0.5% A | 4.0% ABCD | 1.3% BC | 2.0% AB |
| | C29 | 0.4% B | 0.4% A | 2.4% D | 9.0% AB | 2.1% AB |
| | N33 | 0.9% AB | 0.7% A | 3.6% ABCD | 4.7% ABC | 2.3% AB |
| | p-value | < 0.001 | 0.004 | < 0.001 | < 0.001 | 0.020 |
| Waste rock | W08 | 1.2% A | 1.1% AB | 1.2% A | 0.0% B | 0.5% A |
| | W09 | 1.2% A | 1.2% AB | 1.9% A | 0.1% B | 0.8% A |
| | W10 | 0.9% A | 1.0% AB | 1.4% A | 0.1% B | 0.7% A |
| | W13 | 1.1% A | 1.2% AB | 1.5% A | 0.4% B | 0.4% A |
| | N16 | 1.2% A | 1.6% A | 1.7% A | 0.0% B | 0.9% A |
| | C21 | 1.0% A | 1.4% AB | 1.1% A | 0.0% B | 0.3% A |
| | C23 | 0.8% A | 1.2% AB | 1.8% A | 1.5% A | 0.8% A |
| | C25 | 0.8% A | 1.3% AB | 1.3% A | 0.0% B | 0.7% A |
| | C29 | 0.9% A | 1.1% AB | 1.3% A | 0.3% B | 0.9% A |
| | N33 | 0.9% A | 0.9% B | 1.5% A | 0.1% B | 0.6% A |
| | p-value | 0.781 | 0.017 | 0.955 | < 0.001 | 0.303 |

**Supplementary Table 8.** Pairwise comparison between genotypes in each substrate type in the greenhouse experiment for bacterial and fungal taxa relative abundance. Two-way ANOVAs were used to discern how substrate type, genotype and their interaction influenced taxa relative abundance. Additional analyses were performed for each substrate type separately and between substrate types to better assess the indivudial effects of genotype and substrate type.

| **Bacteria** | | *Isosphaeraceae_g* | *Myxococcales_f_g* | *Opitutaceae_g* | *Opitutus* | *Pedosphaerales_f_g* |
|---|---|---|---|---|---|---|
| | Genotype | 0.002 | 0.003 | 0.281 | 0.002 | 0.001 |
| | Substrate type | < 0.001 | < 0.001 | < 0.001 | < 0.001 | < 0.001 |
| | Interaction | 0.001 | 0.567 | 0.328 | < 0.001 | < 0.001 |
| **Pairwise comparison between substrate types** | | | | | | |
| | Control | 0.4% B | 3.3% A | 0.4% B | 0.8% B | 0.7% B |
| | Tailings | 0.3% B | 2.4% B | 2.8% A | 1.2% A | 0.7% B |
| | Waste rock | 1.8% A | 1.5% C | 0.4% B | 0.6% B | 1.3% A |
| | p-value | < 0.001 | < 0.001 | < 0.001 | < 0.001 | < 0.001 |
| **Pairwise comparison by substrate type** | | | | | | |
| Control | W08 | 0.6% A | 3.8% A | 0.4% A | 0.7% A | 0.7% AB |
| | W09 | 0.4% AB | 2.8% A | 0.3% A | 0.7% A | 0.4% B |
| | W10 | 0.3% AB | 3.3% A | 0.3% A | 0.8% A | 0.6% B |
| | W13 | 0.2% B | 4.7% A | 0.5% A | 0.5% A | 0.4% B |
| | N16 | 0.4% AB | 3.3% A | 0.5% A | 1.0% A | 0.6% AB |
| | C21 | 0.4% AB | 1.9% A | 0.3% A | 0.5% A | 1.4% A |
| | C23 | 0.5% AB | 3.0% A | 0.8% A | 1.0% A | 1.1% AB |
| | C25 | 0.3% B | 3.1% A | 0.4% A | 0.7% A | 0.8% AB |
| | C29 | 0.2% B | 3.5% A | 0.2% A | 0.7% A | 0.8% AB |
| | N33 | 0.4% AB | 2.9% A | 0.2% A | 0.9% A | 0.8% AB |
| | p-value | < 0.001 | 0.342 | 0.419 | 0.475 | < 0.001 |
| Tailngs | W08 | 0.3% A | 2.7% A | 3.9% A | 1.0% B | 0.7% A |
| | W09 | 0.6% A | 2.4% A | 1.3% A | 0.9% B | 0.6% A |
| | W10 | 0.4% A | 2.4% A | 3.4% A | 1.2% B | 0.6% A |
| | W13 | 0.3% A | 2.7% A | 4.5% A | 1.1% B | 0.7% A |
| | N16 | 0.4% A | 3.7% A | 4.5% A | 2.7% A | 0.5% A |
| | C21 | 0.5% A | 2.1% A | 1.9% A | 1.2% B | 0.7% A |
| | C23 | 0.2% A | 1.7% A | 1.6% A | 0.9% B | 1.0% A |
| | C25 | 0.3% A | 2.3% A | 1.6% A | 0.7% B | 0.5% A |
| | C29 | 0.2% A | 2.1% A | 2.8% A | 1.2% B | 1.1% A |
| | N33 | 0.3% A | 2.8% A | 3.1% A | 1.2% B | 1.0% A |
| | p-value | 0.055 | 0.098 | 0.107 | < 0.001 | 0.051 |
| Waste rock | W08 | 2.5% A | 1.4% A | 0.4% A | 0.7% A | 2.4% AB |
| | W09 | 1.4% A | 1.4% A | 0.4% A | 0.5% A | 1.1% ABC |
| | W10 | 1.1% A | 1.1% A | 0.2% A | 0.6% A | 1.2% ABC |
| | W13 | 1.0% A | 1.7% A | 0.3% A | 0.4% A | 2.7% A |
| | N16 | 3.0% A | 2.7% A | 0.3% A | 0.5% A | 1.0% ABC |
| | C21 | 2.7% A | 0.9% A | 0.2% A | 0.7% A | 0.7% C |
| | C23 | 1.2% A | 1.6% A | 0.6% A | 1.0% A | 0.9% BC |
| | C25 | 2.2% A | 1.6% A | 0.8% A | 0.7% A | 0.7% C |
| | C29 | 0.9% A | 2.1% A | 0.2% A | 0.4% A | 1.2% ABC |
| | N33 | 1.9% A | 1.0% A | 0.4% A | 0.5% A | 1.2% ABC |
| | p-value | 0.009 | 0.063 | 0.600 | 0.753 | < 0.001 |

**Supplementary Table 8.** Pairwise comparison between genotypes in each substrate type in the greenhouse experiment for bacterial and fungal taxa relative abundance. Two-way ANOVAs were used to discern how substrate type, genotype and their interaction influenced taxa relative abundance. Additional analyses were performed for each substrate type separately and between substrate types to better assess the indivudial effects of genotype and substrate type.

| **Bacteria** | | *Pirellulaceae_g* | *Planctomyces* | *Rhizobiales_f_g* | *Rhodoplanes* | *Rhodospirillaceae_g* |
|---|---|---|---|---|---|---|
| | Genotype | 0.962 | 0.596 | 0.088 | < 0.001 | < 0.001 |
| | Substrate type | < 0.001 | < 0.001 | 0.006 | < 0.001 | < 0.001 |
| | Interaction | 0.068 | 0.001 | 0.186 | < 0.001 | < 0.001 |
| **Pairwise comparison between substrate types** | | | | | | |
| | Control | 1.2% A | 2.5% B | 2.3% A | 3.1% C | 10.2% A |
| | Tailings | 1.2% A | 2.5% B | 1.9% B | 3.9% B | 4.8% C |
| | Waste rock | 0.5% B | 3.7% A | 1.8% B | 5.7% A | 6.7% B |
| | p-value | < 0.001 | 0.004 | 0.015 | < 0.001 | < 0.001 |
| **Pairwise comparison by substrate type** | | | | | | |
| Control | W08 | 0.8% A | 2.2% A | 2.2% A | 2.8% A | 7.8% D |
| | W09 | 1.1% A | 2.4% A | 2.6% A | 2.5% A | 9.8% BCD |
| | W10 | 1.1% A | 2.4% A | 2.0% A | 3.7% A | 11.3% ABC |
| | W13 | 0.8% A | 2.0% A | 1.9% A | 2.7% A | 9.1% CD |
| | N16 | 1.7% A | 2.2% A | 2.4% A | 2.8% A | 8.7% CD |
| | C21 | 1.7% A | 3.1% A | 3.1% A | 3.7% A | 14.8% A |
| | C23 | 1.4% A | 2.6% A | 2.2% A | 3.3% A | 10.3% BCD |
| | C25 | 1.1% A | 2.2% A | 1.7% A | 3.3% A | 9.9% BCD |
| | C29 | 1.2% A | 2.9% A | 2.0% A | 2.9% A | 12.3% AB |
| | N33 | 1.2% A | 2.8% A | 3.0% A | 3.1% A | 9.3% BCD |
| | p-value | 0.201 | 0.697 | 0.238 | 0.098 | < 0.001 |
| Tailngs | W08 | 1.0% A | 2.5% A | 1.9% A | 3.1% B | 4.6% ABC |
| | W09 | 1.3% A | 3.2% A | 1.8% A | 4.4% AB | 4.5% BC |
| | W10 | 1.1% A | 2.2% A | 2.0% A | 3.6% B | 4.2% BC |
| | W13 | 1.9% A | 2.5% A | 2.4% A | 4.0% AB | 5.3% ABC |
| | N16 | 1.5% A | 2.9% A | 1.9% A | 3.6% AB | 6.5% AB |
| | C21 | 1.3% A | 2.5% A | 2.2% A | 4.1% AB | 5.0% ABC |
| | C23 | 1.0% A | 2.5% A | 1.5% A | 3.0% B | 3.7% C |
| | C25 | 1.5% A | 2.9% A | 2.2% A | 5.7% A | 6.6% A |
| | C29 | 0.7% A | 2.5% A | 1.7% A | 3.3% B | 3.9% C |
| | N33 | 1.1% A | 1.8% A | 1.7% A | 4.2% AB | 4.6% ABC |
| | p-value | 0.076 | 0.437 | 0.294 | < 0.001 | < 0.001 |
| Waste rock | W08 | 0.7% A | 2.5% B | 2.3% A | 3.9% B | 6.6% A |
| | W09 | 0.5% A | 2.7% B | 1.8% A | 5.6% AB | 6.9% A |
| | W10 | 0.5% A | 5.9% AB | 1.4% A | 7.0% AB | 7.1% A |
| | W13 | 0.6% A | 2.4% B | 2.1% A | 5.2% AB | 5.8% A |
| | N16 | 0.5% A | 2.7% B | 2.0% A | 6.3% AB | 6.3% A |
| | C21 | 0.5% A | 7.3% A | 2.6% A | 5.4% AB | 5.0% A |
| | C23 | 0.4% A | 2.6% B | 1.7% A | 7.9% A | 8.5% A |
| | C25 | 0.5% A | 4.0% AB | 1.4% A | 6.5% AB | 7.5% A |
| | C29 | 0.6% A | 2.9% B | 1.5% A | 4.9% AB | 7.6% A |
| | N33 | 0.4% A | 4.3% AB | 1.1% A | 5.2% AB | 6.3% A |
| | p-value | 0.969 | < 0.001 | 0.068 | 0.002 | 0.598 |

**Supplementary Table 8.** Pairwise comparison between genotypes in each substrate type in the greenhouse experiment for bacterial and fungal taxa relative abundance. Two-way ANOVAs were used to discern how substrate type, genotype and their interaction influenced taxa relative abundance. Additional analyses were performed for each substrate type separately and between substrate types to better assess the indivudial effects of genotype and substrate type.

| **Bacteria** | | *Rubrivivax* | *Sinobacteraceae_g* | *Solibacterales_f_g* | *Solirubrobacterales_f_g* |
|---|---|---|---|---|---|
| | Genotype | 0.191 | 0.003 | 0.056 | 0.031 |
| | Substrate type | < 0.001 | < 0.001 | < 0.001 | < 0.001 |
| | Interaction | 0.129 | < 0.001 | 0.278 | 0.043 |
| **Pairwise comparison between substrate types** | | | | | |
| | Control | 0.2% B | 1.6% B | 1.4% B | 1.4% A |
| | Tailings | 3.8% A | 0.9% C | 2.1% A | 0.5% C |
| | Waste rock | 0.2% B | 2.8% A | 1.3% B | 0.8% B |
| | p-value | < 0.001 | < 0.001 | < 0.001 | < 0.001 |
| **Pairwise comparison by substrate type** | | | | | |
| Control | W08 | 0.0% A | 1.1% A | 1.9% A | 1.7% A |
| | W09 | 0.0% A | 1.9% A | 1.2% A | 1.7% A |
| | W10 | 0.1% A | 1.4% A | 1.1% A | 1.2% A |
| | W13 | 0.4% A | 2.1% A | 1.4% A | 1.4% A |
| | N16 | 0.1% A | 1.9% A | 1.4% A | 1.2% A |
| | C21 | 0.0% A | 1.6% A | 1.5% A | 1.1% A |
| | C23 | 0.3% A | 1.2% A | 1.8% A | 1.6% A |
| | C25 | 0.1% A | 1.6% A | 1.2% A | 1.0% A |
| | C29 | 0.5% A | 1.6% A | 1.6% A | 1.6% A |
| | N33 | 0.1% A | 1.6% A | 1.2% A | 1.2% A |
| | p-value | 0.717 | 0.142 | 0.361 | 0.014 |
| Tailngs | W08 | 3.2% A | 0.8% A | 1.8% A | 0.5% AB |
| | W09 | 4.7% A | 1.4% A | 2.3% A | 0.9% A |
| | W10 | 2.1% A | 0.7% A | 2.0% A | 0.5% AB |
| | W13 | 2.1% A | 1.0% A | 2.3% A | 0.3% B |
| | N16 | 2.2% A | 1.0% A | 1.6% A | 0.6% AB |
| | C21 | 3.1% A | 0.8% A | 1.9% A | 0.4% B |
| | C23 | 6.7% A | 0.7% A | 2.3% A | 0.6% AB |
| | C25 | 2.8% A | 1.3% A | 1.6% A | 0.6% AB |
| | C29 | 7.8% A | 0.6% A | 2.7% A | 0.5% AB |
| | N33 | 2.4% A | 0.9% A | 2.5% A | 0.4% AB |
| | p-value | 0.009 | 0.009 | 0.026 | 0.011 |
| Waste rock | W08 | 0.0% A | 3.4% AB | 1.2% A | 0.8% A |
| | W09 | 0.1% A | 2.3% B | 1.2% A | 0.7% A |
| | W10 | 0.1% A | 2.8% AB | 1.7% A | 0.7% A |
| | W13 | 1.0% A | 2.5% B | 1.4% A | 0.8% A |
| | N16 | 0.0% A | 3.2% AB | 1.2% A | 0.8% A |
| | C21 | 0.0% A | 4.7% A | 0.8% A | 0.7% A |
| | C23 | 0.2% A | 1.7% B | 1.9% A | 0.6% A |
| | C25 | 0.3% A | 2.5% B | 1.4% A | 0.8% A |
| | C29 | 0.2% A | 2.3% B | 1.5% A | 0.8% A |
| | N33 | 0.0% A | 3.1% AB | 1.3% A | 1.2% A |
| | p-value | 0.423 | < 0.001 | 0.451 | 0.760 |

**Supplementary Table 8.** Pairwise comparison between genotypes in each substrate type in the greenhouse experiment for bacterial and fungal taxa relative abundance. Two-way ANOVAs were used to discern how substrate type, genotype and their interaction influenced taxa relative abundance. Additional analyses were performed for each substrate type separately and between substrate types to better assess the indivudial effects of genotype and substrate type.

| **Bacteria** | | *Sphingobacteriaceae_g* | *Phycisphaerae WD2101_f_g* | *Xanthomonadaceae_g* |
|---|---|---|---|---|
| | Genotype | 0.017 | 0.007 | 0.837 |
| | Substrate type | < 0.001 | < 0.001 | < 0.001 |
| | Interaction | 0.674 | 0.114 | 0.063 |
| **Pairwise comparison between substrate types** | | | | |
| | Control | 0.9% B | 3.2% A | 0.1% B |
| | Tailings | 0.2% C | 0.9% C | 0.0% C |
| | Waste rock | 2.1% A | 2.0% B | 6.4% A |
| | p-value | < 0.001 | < 0.001 | < 0.001 |
| **Pairwise comparison by substrate type** | | | | |
| Control | W08 | 0.9% A | 4.0% A | 0.1% A |
| | W09 | 0.7% A | 4.9% A | 0.1% A |
| | W10 | 1.1% A | 2.9% A | 0.1% A |
| | W13 | 0.7% A | 2.8% A | 0.1% A |
| | N16 | 0.6% A | 2.2% A | 0.0% A |
| | C21 | 1.4% A | 4.1% A | 0.1% A |
| | C23 | 0.5% A | 2.8% A | 0.0% A |
| | C25 | 1.2% A | 2.5% A | 0.1% A |
| | C29 | 0.6% A | 2.7% A | 0.1% A |
| | N33 | 1.3% A | 2.7% A | 0.1% A |
| | p-value | 0.749 | 0.040 | 0.873 |
| Tailngs | W08 | 0.1% A | 1.1% AB | 0.0% A |
| | W09 | 0.1% A | 0.9% AB | 0.0% A |
| | W10 | 0.2% A | 0.6% AB | 0.0% A |
| | W13 | 0.4% A | 1.1% AB | 0.0% A |
| | N16 | 0.2% A | 0.8% AB | 0.0% A |
| | C21 | 0.2% A | 1.3% A | 0.0% A |
| | C23 | 0.2% A | 0.6% B | 0.1% A |
| | C25 | 0.5% A | 1.4% A | 0.0% A |
| | C29 | 0.1% A | 0.8% AB | 0.0% A |
| | N33 | 0.3% A | 0.9% AB | 0.0% A |
| | p-value | 0.057 | 0.002 | 0.300 |
| Waste rock | W08 | 1.8% A | 2.9% A | 5.4% A |
| | W09 | 1.4% A | 2.6% A | 5.4% A |
| | W10 | 2.5% A | 1.7% A | 6.6% A |
| | W13 | 1.8% A | 2.5% A | 7.5% A |
| | N16 | 2.3% A | 1.5% A | 6.1% A |
| | C21 | 3.3% A | 1.0% A | 9.5% A |
| | C23 | 1.7% A | 2.5% A | 4.2% A |
| | C25 | 2.2% A | 1.4% A | 8.4% A |
| | C29 | 1.4% A | 2.0% A | 3.4% A |
| | N33 | 2.7% A | 1.3% A | 8.1% A |
| | p-value | 0.182 | 0.474 | 0.039 |

**Supplementary Table 8.** Pairwise comparison between genotypes in each substrate type in the greenhouse experiment for bacterial and fungal taxa relative abundance. Two-way ANOVAs were used to discern how substrate type, genotype and their interaction influenced taxa relative abundance. Additional analyses were performed for each substrate type separately and between substrate types to better assess the indivudial effects of genotype and substrate type.

| **Fungi** | | *Acidea* | *Alternaria* | *Articulospora* | *Cadophora* | *Cephalothecaceae_g* | *Chrysosporium* |
|---|---|---|---|---|---|---|---|
| | Genotype | 0.293 | 0.306 | 0.634 | 0.005 | 0.753 | 0.904 |
| | Substrate type | < 0.001 | < 0.001 | < 0.001 | 0.081 | 0.089 | < 0.001 |
| | Interaction | 0.626 | 0.790 | 0.809 | 0.329 | 0.107 | 0.133 |
| **Pairwise comparison between substrate types** | | | | | | | |
| | Control | 0.0% B | 2.1% A | 0.8% A | 0.2% A | 1.7% A | 18.1% A |
| | Tailings | 0.0% B | 0.7% B | 0.2% B | 0.5% A | 1.2% A | 14.3% B |
| | Waste rock | 3.3% A | 0.7% B | 0.3% B | 0.1% A | 1.5% A | 11.6% B |
| | p-value | < 0.001 | < 0.001 | < 0.001 | 0.101 | 0.079 | < 0.001 |
| **Pairwise comparison by substrate type** | | | | | | | |
| Control | W08 | 0.0% A | 1.6% A | 0.6% A | 0.2% A | 1.5% A | 14.0% A |
| | W09 | 0.0% A | 2.5% A | 1.1% A | 0.1% A | 1.2% A | 17.8% A |
| | W10 | 0.0% A | 2.2% A | 0.8% A | 0.1% A | 1.9% A | 18.9% A |
| | W13 | 0.0% A | 2.6% A | 1.0% A | 0.1% A | 1.5% A | 19.9% A |
| | N16 | 0.0% A | 2.1% A | 0.9% A | 0.0% A | 1.4% A | 18.7% A |
| | C21 | 0.0% A | 1.1% A | 0.7% A | 0.1% A | 1.2% A | 20.9% A |
| | C23 | 0.0% A | 2.2% A | 0.8% A | 0.8% A | 2.1% A | 16.6% A |
| | C25 | 0.0% A | 2.5% A | 0.9% A | 0.1% A | 2.2% A | 21.0% A |
| | C29 | 0.0% A | 1.6% A | 0.5% A | 0.1% A | 1.2% A | 15.2% A |
| | N33 | 0.0% A | 2.7% A | 1.0% A | 0.2% A | 2.3% A | 19.0% A |
| | p-value | 0.630 | 0.344 | 0.557 | 0.144 | 0.761 | 0.496 |
| Tailngs | W08 | 0.0% A | 1.1% A | 0.3% A | 0.3% A | 2.0% A | 17.7% A |
| | W09 | 0.0% A | 0.3% A | 0.1% A | 2.1% A | 0.9% A | 10.3% A |
| | W10 | 0.0% A | 0.9% A | 0.3% A | 0.4% A | 1.7% A | 16.4% A |
| | W13 | 0.0% A | 1.2% A | 0.4% A | 0.2% A | 0.7% A | 11.1% A |
| | N16 | 0.0% A | 0.9% A | 0.3% A | 0.1% A | 1.7% A | 12.6% A |
| | C21 | 0.0% A | 0.3% A | 0.2% A | 0.3% A | 1.4% A | 20.7% A |
| | C23 | 0.0% A | 0.9% A | 0.3% A | 0.7% A | 0.7% A | 12.0% A |
| | C25 | 0.0% A | 0.6% A | 0.3% A | 0.3% A | 1.5% A | 11.6% A |
| | C29 | 0.0% A | 0.3% A | 0.2% A | 0.1% A | 0.8% A | 16.8% A |
| | N33 | 0.0% A | 0.5% A | 0.2% A | 0.3% A | 1.4% A | 13.6% A |
| | p-value | 0.261 | 0.893 | 0.968 | 0.091 | 0.045 | 0.107 |
| Waste rock | W08 | 3.4% A | 1.0% A | 0.5% A | 0.2% A | 1.3% A | 9.0% A |
| | W09 | 3.7% A | 0.9% A | 0.4% A | 0.1% A | 2.1% A | 14.8% A |
| | W10 | 2.6% A | 0.6% A | 0.1% A | 0.0% A | 0.8% A | 9.9% A |
| | W13 | 2.0% A | 0.8% A | 0.3% A | 0.4% A | 1.9% A | 15.5% A |
| | N16 | 2.1% A | 0.6% A | 0.2% A | 0.0% A | 1.5% A | 10.5% A |
| | C21 | 2.9% A | 0.2% A | 0.1% A | 0.1% A | 1.1% A | 8.8% A |
| | C23 | 3.9% A | 1.4% A | 0.7% A | 0.4% A | 1.3% A | 12.3% A |
| | C25 | 3.2% A | 0.2% A | 0.1% A | 0.0% A | 1.3% A | 11.0% A |
| | C29 | 1.9% A | 0.9% A | 0.2% A | 0.1% A | 1.7% A | 13.4% A |
| | N33 | 5.5% A | 0.5% A | 0.2% A | 0.1% A | 1.6% A | 11.6% A |
| | p-value | 0.540 | 0.413 | 0.431 | 0.079 | 0.480 | 0.743 |

**Supplementary Table 8.** Pairwise comparison between genotypes in each substrate type in the greenhouse experiment for bacterial and fungal taxa relative abundance. Two-way ANOVAs were used to discern how substrate type, genotype and their interaction influenced taxa relative abundance. Additional analyses were performed for each substrate type separately and between substrate types to better assess the indivudial effects of genotype and substrate type.

| **Fungi** | | *Ciliophora* | *Cladosporium* | *Eurotiomycetes_o_f_g* | *Fusarium* | *Gibberella* |
|---|---|---|---|---|---|---|
| | Genotype | < 0.001 | 0.385 | 0.008 | 0.282 | 0.145 |
| | Substrate type | < 0.001 | < 0.001 | < 0.001 | 0.681 | < 0.001 |
| | Interaction | 0.136 | 0.917 | 0.008 | 0.842 | 0.732 |
| **Pairwise comparison between substrate types** | | | | | | |
| | Control | 15.0% A | 0.8% A | 0.4% B | 2.1% A | 0.9% A |
| | Tailings | 4.0% B | 0.3% B | 0.7% A | 2.1% A | 0.4% B |
| | Waste rock | 2.7% B | 0.3% B | 0.8% A | 2.6% A | 0.3% B |
| | p-value | < 0.001 | < 0.001 | < 0.001 | 0.676 | < 0.001 |
| **Pairwise comparison by substrate type** | | | | | | |
| Control | W08 | 26.9% A | 0.7% A | 0.8% A | 1.8% A | 0.6% A |
| | W09 | 17.1% AB | 0.7% A | 0.4% A | 2.4% A | 1.1% A |
| | W10 | 16.8% AB | 0.9% A | 0.1% A | 2.0% A | 1.1% A |
| | W13 | 14.5% AB | 1.1% A | 0.7% A | 2.4% A | 1.2% A |
| | N16 | 1.7% B | 0.9% A | 0.3% A | 2.1% A | 0.9% A |
| | C21 | 13.6% AB | 0.2% A | 0.2% A | 1.2% A | 0.4% A |
| | C23 | 21.6% AB | 0.9% A | 0.6% A | 2.1% A | 1.0% A |
| | C25 | 14.8% AB | 0.8% A | 0.3% A | 2.2% A | 1.2% A |
| | C29 | 7.4% AB | 0.5% A | 0.2% A | 1.5% A | 0.7% A |
| | N33 | 10.5% AB | 0.8% A | 0.3% A | 2.6% A | 1.0% A |
| | p-value | 0.015 | 0.521 | 0.044 | 0.343 | 0.486 |
| Tailngs | W08 | 7.3% AB | 0.3% A | 1.1% A | 0.9% A | 0.3% A |
| | W09 | 3.6% AB | 0.6% A | 0.7% A | 4.9% A | 0.2% A |
| | W10 | 2.9% AB | 0.3% A | 1.0% A | 3.3% A | 0.4% A |
| | W13 | 2.1% AB | 0.6% A | 0.9% A | 1.8% A | 0.6% A |
| | N16 | 2.3% AB | 0.3% A | 0.4% A | 2.7% A | 0.7% A |
| | C21 | 1.3% B | 0.2% A | 0.7% A | 2.0% A | 0.1% A |
| | C23 | 6.5% AB | 0.3% A | 0.9% A | 1.5% A | 0.5% A |
| | C25 | 0.5% B | 0.4% A | 0.3% A | 1.6% A | 0.5% A |
| | C29 | 0.8% B | 0.1% A | 0.7% A | 1.4% A | 0.3% A |
| | N33 | 11.2% A | 0.2% A | 0.5% A | 1.1% A | 0.2% A |
| | p-value | 0.001 | 0.762 | 0.110 | 0.656 | 0.720 |
| Waste rock | W08 | 5.5% A | 0.4% A | 0.5% A | 2.1% A | 0.4% A |
| | W09 | 4.6% A | 0.4% A | 1.2% A | 3.7% A | 0.5% A |
| | W10 | 0.4% A | 0.2% A | 0.4% A | 0.8% A | 0.3% A |
| | W13 | 1.7% A | 0.3% A | 1.3% A | 4.0% A | 0.4% A |
| | N16 | 1.0% A | 0.3% A | 1.1% A | 0.7% A | 0.2% A |
| | C21 | 1.3% A | 0.1% A | 0.3% A | 0.6% A | 0.1% A |
| | C23 | 1.9% A | 0.4% A | 1.0% A | 2.5% A | 0.7% A |
| | C25 | 0.9% A | 0.2% A | 1.3% A | 2.5% A | 0.2% A |
| | C29 | 2.7% A | 0.4% A | 1.0% A | 6.1% A | 0.3% A |
| | N33 | 4.0% A | 0.2% A | 0.6% A | 2.5% A | 0.3% A |
| | p-value | 0.435 | 0.730 | 0.002 | 0.729 | 0.176 |

**Supplementary Table 8.** Pairwise comparison between genotypes in each substrate type in the greenhouse experiment for bacterial and fungal taxa relative abundance. Two-way ANOVAs were used to discern how substrate type, genotype and their interaction influenced taxa relative abundance. Additional analyses were performed for each substrate type separately and between substrate types to better assess the indivudial effects of genotype and substrate type.

| **Fungi** | | *Lecythophora* | *Leptosphaeria* | *Lindtneria* | *Meliniomyces* | *Mortierella* | *Pezoloma* |
|---|---|---|---|---|---|---|---|
| | Genotype | 0.039 | 0.753 | 0.901 | 0.196 | 0.386 | 0.209 |
| | Substrate type | 0.253 | 0.004 | 0.994 | < 0.001 | < 0.001 | < 0.001 |
| | Interaction | 0.519 | 0.203 | 0.527 | 0.046 | 0.478 | 0.029 |
| **Pairwise comparison between substrate types** | | | | | | | |
| | Control | 0.3% A | 0.0% B | 0.7% A | 0.0% B | 0.6% A | 0.0% B |
| | Tailings | 0.3% A | 0.9% A | 0.6% A | 0.1% B | 0.2% B | 0.0% B |
| | Waste rock | 0.5% A | 0.0% B | 0.6% A | 2.3% A | 0.2% B | 6.6% A |
| | p-value | 0.194 | < 0.001 | 0.986 | 0.006 | < 0.001 | < 0.001 |
| **Pairwise comparison by substrate type** | | | | | | | |
| Control | W08 | 0.5% A | 0.0% A | 0.8% A | 0.1% A | 0.3% A | 0.0% A |
| | W09 | 0.3% A | 0.0% A | 0.0% A | 0.0% A | 0.7% A | 0.0% A |
| | W10 | 0.1% A | 0.0% A | 0.6% A | 0.0% A | 0.7% A | 0.0% A |
| | W13 | 0.0% A | 0.0% A | 0.2% A | 0.0% A | 0.6% A | 0.0% A |
| | N16 | 0.0% A | 0.0% A | 0.2% A | 0.1% A | 0.7% A | 0.0% A |
| | C21 | 0.5% A | 0.0% A | 0.0% A | 0.0% A | 0.3% A | 0.1% A |
| | C23 | 0.4% A | 0.0% A | 0.6% A | 0.1% A | 0.6% A | 0.0% A |
| | C25 | 0.3% A | 0.0% A | 0.6% A | 0.0% A | 0.7% A | 0.0% A |
| | C29 | 0.0% A | 0.0% A | 1.0% A | 0.0% A | 0.4% A | 0.0% A |
| | N33 | 0.8% A | 0.0% A | 2.4% A | 0.0% A | 0.7% A | 0.0% A |
| | p-value | 0.034 | 0.508 | 0.770 | 0.202 | 0.153 | 0.169 |
| Tailngs | W08 | 0.7% A | 0.4% A | 0.7% A | 0.1% A | 0.2% A | 0.0% A |
| | W09 | 0.1% A | 1.1% A | 0.1% A | 0.0% A | 0.1% A | 0.0% A |
| | W10 | 0.1% A | 2.1% A | 1.5% A | 0.6% A | 0.2% A | 0.0% A |
| | W13 | 0.2% A | 0.1% A | 1.9% A | 0.0% A | 0.2% A | 0.0% A |
| | N16 | 0.0% A | 5.2% A | 0.0% A | 0.0% A | 0.2% A | 0.0% A |
| | C21 | 0.3% A | 0.3% A | 1.5% A | 0.0% A | 0.1% A | 0.0% A |
| | C23 | 0.2% A | 0.8% A | 0.0% A | 0.0% A | 0.3% A | 0.0% A |
| | C25 | 0.1% A | 0.1% A | 0.0% A | 0.0% A | 0.2% A | 0.0% A |
| | C29 | 0.2% A | 0.1% A | 0.0% A | 0.0% A | 0.1% A | 0.0% A |
| | N33 | 0.7% A | 0.1% A | 0.6% A | 0.0% A | 0.1% A | 0.1% A |
| | p-value | 0.690 | 0.116 | 0.572 | 0.486 | 0.947 | 0.030 |
| Waste rock | W08 | 0.4% A | 0.0% A | 0.4% A | 2.3% AB | 0.2% A | 5.7% A |
| | W09 | 0.3% A | 0.0% A | 0.9% A | 0.3% B | 0.4% A | 5.4% A |
| | W10 | 0.1% A | 0.0% A | 0.2% A | 2.9% AB | 0.2% A | 9.6% A |
| | W13 | 0.9% A | 0.0% A | 1.4% A | 0.0% B | 0.2% A | 1.4% A |
| | N16 | 0.2% A | 0.0% A | 1.7% A | 2.1% AB | 0.2% A | 8.4% A |
| | C21 | 0.2% A | 0.0% A | 0.4% A | 1.4% AB | 0.0% A | 9.3% A |
| | C23 | 0.8% A | 0.0% A | 0.5% A | 2.4% AB | 0.3% A | 7.9% A |
| | C25 | 1.1% A | 0.0% A | 0.8% A | 0.1% B | 0.1% A | 8.5% A |
| | C29 | 0.7% A | 0.0% A | 0.4% A | 0.2% B | 0.4% A | 3.1% A |
| | N33 | 0.4% A | 0.0% A | 0.4% A | 9.0% A | 0.1% A | 6.5% A |
| | p-value | 0.340 | 0.730 | 0.554 | 0.002 | 0.124 | 0.261 |

**Supplementary Table 8.** Pairwise comparison between genotypes in each substrate type in the greenhouse experiment for bacterial and fungal taxa relative abundance. Two-way ANOVAs were used to discern how substrate type, genotype and their interaction influenced taxa relative abundance. Additional analyses were performed for each substrate type separately and between substrate types to better assess the indivudial effects of genotype and substrate type.

| **Fungi** | | *Phaeosphaeriaceae_g* | *Pleosporale_f_g* | *Pleosporales_ fam_Incertae_sedis_g* | *Pyrenopeziza* |
|---|---|---|---|---|---|
| | Genotype | 0.422 | 0.411 | 0.265 | 0.384 |
| | Substrate type | < 0.001 | < 0.001 | < 0.001 | < 0.001 |
| | Interaction | 0.755 | 0.522 | 0.933 | 0.622 |
| **Pairwise comparison between substrate types** | | | | | |
| | Control | 0.7% A | 0.5% A | 3.1% A | 0.0% B |
| | Tailings | 0.2% B | 0.2% B | 0.9% B | 0.0% B |
| | Waste rock | 0.2% B | 0.2% B | 1.0% B | 0.9% A |
| | p-value | < 0.001 | < 0.001 | < 0.001 | < 0.001 |
| **Pairwise comparison by substrate type** | | | | | |
| Control | W08 | 0.6% A | 0.4% A | 2.3% A | 0.0% A |
| | W09 | 1.1% A | 0.5% A | 3.6% A | 0.0% A |
| | W10 | 0.6% A | 0.5% A | 3.5% A | 0.0% A |
| | W13 | 0.9% A | 0.6% A | 3.6% A | 0.0% A |
| | N16 | 0.6% A | 0.6% A | 3.1% A | 0.0% A |
| | C21 | 0.4% A | 0.3% A | 1.5% A | 0.0% A |
| | C23 | 0.7% A | 0.6% A | 3.4% A | 0.0% A |
| | C25 | 0.9% A | 0.4% A | 3.3% A | 0.0% A |
| | C29 | 0.5% A | 0.4% A | 2.6% A | 0.0% A |
| | N33 | 0.8% A | 0.5% A | 3.5% A | 0.0% A |
| | p-value | 0.190 | 0.511 | 0.425 | 0.987 |
| Tailngs | W08 | 0.3% A | 0.4% A | 1.2% A | 0.0% A |
| | W09 | 0.1% A | 0.0% A | 0.4% A | 0.0% A |
| | W10 | 0.3% A | 0.4% A | 1.0% A | 0.0% A |
| | W13 | 0.3% A | 0.2% A | 1.5% A | 0.0% A |
| | N16 | 0.2% A | 0.2% A | 0.8% A | 0.0% A |
| | C21 | 0.1% A | 0.2% A | 0.6% A | 0.0% A |
| | C23 | 0.2% A | 0.2% A | 1.2% A | 0.0% A |
| | C25 | 0.2% A | 0.1% A | 1.2% A | 0.0% A |
| | C29 | 0.1% A | 0.1% A | 0.4% A | 0.0% A |
| | N33 | 0.2% A | 0.1% A | 0.7% A | 0.0% A |
| | p-value | 0.892 | 0.282 | 0.952 | 0.437 |
| Waste rock | W08 | 0.3% A | 0.2% A | 1.2% A | 0.6% A |
| | W09 | 0.3% A | 0.2% A | 1.7% A | 1.2% A |
| | W10 | 0.1% A | 0.1% A | 1.0% A | 0.7% A |
| | W13 | 0.2% A | 0.3% A | 1.0% A | 0.8% A |
| | N16 | 0.2% A | 0.2% A | 0.5% A | 0.7% A |
| | C21 | 0.1% A | 0.1% A | 0.3% A | 1.0% A |
| | C23 | 0.3% A | 0.3% A | 1.6% A | 1.0% A |
| | C25 | 0.2% A | 0.1% A | 0.6% A | 0.6% A |
| | C29 | 0.3% A | 0.3% A | 1.0% A | 0.9% A |
| | N33 | 0.2% A | 0.1% A | 0.8% A | 1.5% A |
| | p-value | 0.580 | 0.681 | 0.300 | 0.334 |

**Supplementary Table 8.** Pairwise comparison between genotypes in each substrate type in the greenhouse experiment for bacterial and fungal taxa relative abundance. Two-way ANOVAs were used to discern how substrate type, genotype and their interaction influenced taxa relative abundance. Additional analyses were performed for each substrate type separately and between substrate types to better assess the indivudial effects of genotype and substrate type.

| **Fungi** | | *Pyrenophora* | *Russula* | *Sebacinales_f_g* | *Sordariales_f_g* | *Sphaerosporella* | *Tomentella* |
|---|---|---|---|---|---|---|---|
| | Genotype | 0.514 | 0.892 | 0.008 | 0.639 | 0.031 | 0.395 |
| | Substrate type | < 0.001 | < 0.001 | < 0.001 | < 0.001 | < 0.001 | < 0.001 |
| | Interaction | 0.649 | 0.005 | 0.039 | 0.252 | 0.671 | 0.527 |
| **Pairwise comparison between substrate types** | | | | | | | |
| | Control | 0.9% A | 0.0% B | 0.3% B | 0.1% B | 9.5% A | 31.6% A |
| | Tailings | 0.2% B | 0.1% B | 1.7% A | 33.0% A | 7.7% A | 20.1% B |
| | Waste rock | 0.3% B | 6.0% A | 0.4% B | 0.5% B | 3.0% B | 40.5% A |
| | p-value | < 0.001 | < 0.001 | < 0.001 | < 0.001 | < 0.001 | < 0.001 |
| **Pairwise comparison by substrate type** | | | | | | | |
| Control | W08 | 0.6% A | 0.0% A | 0.3% A | 0.0% A | 13.6% A | 23.0% A |
| | W09 | 0.7% A | 0.0% A | 0.4% A | 0.1% A | 6.5% A | 31.2% A |
| | W10 | 0.9% A | 0.0% A | 0.1% A | 0.1% A | 3.8% A | 33.0% A |
| | W13 | 1.2% A | 0.0% A | 0.2% A | 0.0% A | 3.7% A | 33.1% A |
| | N16 | 1.0% A | 0.0% A | 0.3% A | 0.1% A | 4.0% A | 51.8% A |
| | C21 | 0.3% A | 0.0% A | 0.7% A | 0.0% A | 19.3% A | 30.3% A |
| | C23 | 0.7% A | 0.0% A | 0.3% A | 0.1% A | 12.0% A | 21.8% A |
| | C25 | 1.2% A | 0.0% A | 0.2% A | 0.1% A | 11.2% A | 24.0% A |
| | C29 | 0.6% A | 0.0% A | 0.1% A | 0.0% A | 14.5% A | 44.1% A |
| | N33 | 1.1% A | 0.0% A | 0.4% A | 0.1% A | 8.2% A | 30.2% A |
| | p-value | 0.279 | 0.910 | 0.048 | 0.634 | 0.246 | 0.211 |
| Tailngs | W08 | 0.2% A | 0.0% A | 1.7% AB | 28.1% A | 3.0% A | 21.0% A |
| | W09 | 0.1% A | 0.0% A | 2.2% AB | 27.7% A | 6.2% A | 28.4% A |
| | W10 | 0.3% A | 1.1% A | 1.9% AB | 46.6% A | 1.5% A | 5.8% A |
| | W13 | 0.5% A | 0.0% A | 1.2% B | 27.7% A | 10.2% A | 28.6% A |
| | N16 | 0.1% A | 0.0% A | 0.4% B | 34.7% A | 7.5% A | 19.2% A |
| | C21 | 0.2% A | 0.0% A | 5.7% A | 31.2% A | 9.9% A | 9.0% A |
| | C23 | 0.3% A | 0.0% A | 1.3% B | 27.3% A | 14.7% A | 17.4% A |
| | C25 | 0.2% A | 0.0% A | 1.3% B | 44.2% A | 11.1% A | 17.5% A |
| | C29 | 0.2% A | 0.0% A | 0.2% B | 30.9% A | 8.6% A | 31.2% A |
| | N33 | 0.1% A | 0.0% A | 1.0% B | 33.8% A | 2.3% A | 23.3% A |
| | p-value | 0.950 | 0.410 | < 0.001 | 0.211 | 0.753 | 0.695 |
| Waste rock | W08 | 0.4% A | 7.6% AB | 1.5% A | 0.0% A | 2.1% A | 43.5% A |
| | W09 | 0.4% A | 1.4% B | 0.4% A | 0.0% A | 6.1% A | 37.5% A |
| | W10 | 0.2% A | 16.2% A | 0.2% A | 0.0% A | 1.2% A | 45.9% A |
| | W13 | 0.3% A | 0.0% AB | 0.4% A | 0.0% A | 2.5% A | 50.1% A |
| | N16 | 0.2% A | 3.1% AB | 0.2% A | 0.1% A | 1.0% A | 53.4% A |
| | C21 | 0.0% A | 5.3% AB | 0.2% A | 5.1% A | 0.6% A | 51.7% A |
| | C23 | 0.5% A | 2.3% AB | 0.4% A | 0.0% A | 5.6% A | 32.5% A |
| | C25 | 0.2% A | 14.1% AB | 0.2% A | 0.0% A | 1.0% A | 40.1% A |
| | C29 | 0.3% A | 0.6% AB | 0.2% A | 0.0% A | 8.4% A | 26.2% A |
| | N33 | 0.2% A | 6.7% AB | 0.4% A | 0.2% A | 0.9% A | 34.1% A |
| | p-value | 0.152 | 0.001 | 0.839 | 0.390 | 0.069 | 0.542 |

**Supplementary Table 8.** Pairwise comparison between genotypes in each substrate type in the greenhouse experiment for bacterial and fungal taxa relative abundance. Two-way ANOVAs were used to discern how substrate type, genotype and their interaction influenced taxa relative abundance. Additional analyses were performed for each substrate type separately and between substrate types to better assess the indivudial effects of genotype and substrate type.

| **Fungi** | | *Trichoderma* | | *Vibrisseaceae_g* | |
|---|---|---|---|---|---|
| | Genotype | 0.815 | | 0.002 | |
| | Substrate type | 0.012 | | < 0.001 | |
| | Interaction | 0.599 | | 0.227 | |
| **Pairwise comparison between substrate types** | | | | | |
| | Control | 0.4% | B | 0.2% | B |
| | Tailings | 0.7% | AB | 0.0% | C |
| | Waste rock | 1.8% | A | 0.6% | A |
| | p-value | 0.009 | | < 0.001 | |
| **Pairwise comparison by substrate type** | | | | | |
| Control | W08 | 0.9% | A | 0.1% | A |
| | W09 | 0.1% | A | 0.4% | A |
| | W10 | 0.2% | A | 0.0% | A |
| | W13 | 0.1% | A | 0.0% | A |
| | N16 | 0.1% | A | 0.2% | A |
| | C21 | 0.1% | A | 0.5% | A |
| | C23 | 0.9% | A | 0.3% | A |
| | C25 | 0.5% | A | 0.1% | A |
| | C29 | 0.1% | A | 0.1% | A |
| | N33 | 0.6% | A | 0.4% | A |
| | p-value | 0.754 | | 0.052 | |
| Tailngs | W08 | 0.5% | A | 0.0% | B |
| | W09 | 1.0% | A | 0.0% | B |
| | W10 | 0.8% | A | 0.0% | B |
| | W13 | 0.4% | A | 0.0% | B |
| | N16 | 1.5% | A | 0.0% | AB |
| | C21 | 0.4% | A | 0.1% | A |
| | C23 | 1.0% | A | 0.0% | B |
| | C25 | 0.1% | A | 0.0% | AB |
| | C29 | 0.9% | A | 0.0% | B |
| | N33 | 0.3% | A | 0.0% | AB |
| | p-value | 0.601 | | 0.005 | |
| Waste rock | W08 | 0.3% | A | 1.3% | A |
| | W09 | 1.5% | A | 0.4% | A |
| | W10 | 0.3% | A | 0.4% | A |
| | W13 | 4.5% | A | 0.2% | A |
| | N16 | 2.7% | A | 0.2% | A |
| | C21 | 1.0% | A | 0.7% | A |
| | C23 | 2.7% | A | 1.3% | A |
| | C25 | 1.4% | A | 0.3% | A |
| | C29 | 0.8% | A | 0.1% | A |
| | N33 | 4.0% | A | 0.4% | A |
| | p-value | 0.542 | | 0.541 | |
