## Supplementary Table 9 for "Plant Genotype Influences Physicochemical Properties of Substrate as well as Bacterial and Fungal Assemblages in the Rhizosphere of Balsam Poplar"

**Supplementary Table 9.** Spearman linear correlation analyses between bacterial and fungal taxa relative abundance and chemical properties of substrates in greenhouse samples. Weak correlations (>|0.3|) are highlighted in red; moderate correlations (>|0.5|) are highlighted in yellow; strong correlations (>|0.7|) are highlighted in green. CEC: Cation exchange capacity; BCSR: Base cation saturation ratio.

| **Bacteria** | **C total** | **N total** | **S total** | **pH** | **P** | **K** | **Ca** | **Mg** | **Mn** | **Fe** | **Na** | **CEC** | **BCSR** |
|---|---|---|---|---|---|---|---|---|---|---|---|---|---|
| *Anaerolineae SBR1031 A4b_g* | -0.37 | -0.38 | -0.49 | 0.48 | -0.40 | -0.17 | -0.29 | 0.04 | 0.05 | -0.09 | -0.23 | -0.45 | 0.19 |
| *Acidobacteriaceae_g* | 0.19 | 0.24 | 0.76 | -0.73 | 0.28 | -0.17 | 0.16 | -0.29 | -0.51 | 0.46 | 0.04 | 0.32 | -0.56 |
| *Bradyrhizobium* | -0.46 | -0.48 | 0.36 | -0.52 | -0.42 | -0.52 | -0.47 | -0.59 | -0.65 | 0.65 | -0.50 | -0.41 | -0.65 |
| *Caulobacteraceae_g* | -0.09 | -0.07 | 0.72 | -0.84 | 0.00 | -0.35 | -0.12 | -0.46 | -0.62 | 0.63 | -0.15 | 0.03 | -0.68 |
| *Chitinophagaceae_g* | 0.55 | 0.51 | -0.43 | 0.55 | 0.40 | 0.66 | 0.49 | 0.70 | 0.68 | -0.71 | 0.60 | 0.43 | 0.71 |
| *Acidimicrobiales EB1017_g* | -0.16 | -0.21 | -0.66 | 0.63 | -0.26 | 0.16 | -0.14 | 0.24 | 0.47 | -0.42 | 0.02 | -0.26 | 0.47 |
| *Alphaproteobacteria Ellin329_f_g* | 0.38 | 0.38 | 0.63 | -0.72 | 0.36 | 0.10 | 0.31 | -0.07 | -0.25 | 0.34 | 0.30 | 0.47 | -0.35 |
| *Betaproteobacteria Ellin6067_f_g* | 0.28 | 0.30 | 0.25 | -0.21 | 0.25 | 0.09 | 0.27 | 0.11 | -0.10 | 0.06 | 0.23 | 0.32 | -0.10 |
| *Frankiaceae_g* | 0.47 | 0.46 | 0.07 | 0.09 | 0.40 | 0.42 | 0.46 | 0.37 | 0.23 | -0.37 | 0.42 | 0.48 | 0.29 |
| *Gaiellaceae_g* | 0.16 | 0.19 | 0.64 | -0.60 | 0.27 | -0.13 | 0.12 | -0.24 | -0.43 | 0.38 | 0.05 | 0.27 | -0.46 |
| *Gemmataceae_g* | -0.71 | -0.75 | -0.30 | 0.25 | -0.76 | -0.44 | -0.67 | -0.35 | -0.29 | 0.15 | -0.63 | -0.78 | -0.20 |
| *Geobacter* | -0.54 | -0.51 | -0.41 | 0.29 | -0.51 | -0.29 | -0.47 | -0.21 | -0.01 | 0.15 | -0.43 | -0.56 | -0.06 |
| *Acidobacteria-6 iii1-15_f_g* | -0.43 | -0.43 | -0.68 | 0.62 | -0.45 | -0.03 | -0.30 | 0.15 | 0.29 | -0.30 | -0.22 | -0.47 | 0.35 |
| *Isosphaeraceae_g* | -0.04 | -0.03 | 0.68 | -0.73 | -0.01 | -0.37 | -0.11 | -0.46 | -0.65 | 0.55 | -0.16 | 0.03 | -0.64 |
| *Myxococcales_f_g* | 0.25 | 0.25 | -0.45 | 0.64 | 0.16 | 0.32 | 0.23 | 0.48 | 0.46 | -0.62 | 0.24 | 0.12 | 0.59 |
| *Opitutaceae_g* | -0.56 | -0.56 | -0.48 | 0.49 | -0.58 | -0.20 | -0.52 | -0.14 | 0.04 | -0.12 | -0.47 | -0.64 | 0.09 |
| *Opitutus* | -0.27 | -0.29 | -0.41 | 0.35 | -0.27 | -0.06 | -0.24 | -0.02 | 0.22 | -0.14 | -0.23 | -0.31 | 0.17 |
| *Pedosphaerales_f_g* | 0.07 | 0.08 | 0.38 | -0.42 | 0.04 | -0.16 | 0.01 | -0.26 | -0.37 | 0.33 | -0.10 | 0.07 | -0.33 |
| *Pirellulaceae_g* | 0.09 | 0.01 | -0.62 | 0.48 | -0.01 | 0.41 | 0.08 | 0.46 | 0.60 | -0.51 | 0.23 | 0.01 | 0.59 |
| *Planctomyces* | -0.10 | -0.10 | 0.16 | -0.37 | 0.06 | -0.19 | -0.08 | -0.28 | -0.23 | 0.32 | -0.12 | -0.01 | -0.28 |
| *Rhizobiales_f_g* | 0.27 | 0.15 | -0.08 | -0.07 | 0.07 | 0.24 | 0.10 | 0.20 | 0.26 | -0.24 | 0.27 | 0.14 | 0.24 |
| *Rhodoplanes* | -0.34 | -0.32 | 0.48 | -0.64 | -0.18 | -0.40 | -0.25 | -0.46 | -0.60 | 0.68 | -0.28 | -0.18 | -0.63 |
| *Rhodospirillaceae_g* | 0.74 | 0.74 | 0.06 | 0.04 | 0.74 | 0.60 | 0.74 | 0.60 | 0.41 | -0.37 | 0.70 | 0.75 | 0.43 |
| *Rubrivivax* | -0.53 | -0.53 | -0.56 | 0.46 | -0.53 | -0.22 | -0.47 | -0.13 | 0.07 | -0.06 | -0.40 | -0.60 | 0.09 |
| *Sinobacteraceae_g* | 0.25 | 0.28 | 0.70 | -0.71 | 0.33 | -0.10 | 0.24 | -0.18 | -0.37 | 0.38 | 0.17 | 0.39 | -0.40 |
| *Solibacterales_f_g* | -0.43 | -0.37 | -0.38 | 0.39 | -0.44 | -0.24 | -0.39 | -0.10 | -0.04 | -0.02 | -0.33 | -0.51 | 0.05 |
| *Solirubrobacterales_f_g* | 0.60 | 0.65 | 0.08 | 0.11 | 0.61 | 0.42 | 0.56 | 0.39 | 0.33 | -0.41 | 0.49 | 0.56 | 0.37 |
| *Sphingobacteriaceae_g* | 0.25 | 0.29 | 0.64 | -0.70 | 0.37 | -0.03 | 0.24 | -0.16 | -0.35 | 0.38 | 0.18 | 0.40 | -0.42 |
| *Phycisphaerae WD2101_f_g* | 0.69 | 0.67 | 0.19 | -0.02 | 0.56 | 0.47 | 0.59 | 0.42 | 0.25 | -0.35 | 0.58 | 0.62 | 0.28 |
| *Xanthomonadaceae_g* | 0.16 | 0.17 | 0.72 | -0.75 | 0.23 | -0.22 | 0.12 | -0.30 | -0.56 | 0.55 | 0.00 | 0.26 | -0.55 |

**Supplementary Table 9.** Spearman linear correlation analyses between bacterial and fungal taxa relative abundance and chemical properties of substrates in greenhouse samples. Weak correlations (>|0.3|) are highlighted in red; moderate correlations (>|0.5|) are highlighted in yellow; strong correlations (>|0.7|) are highlighted in green. CEC: Cation exchange capacity; BCSR: Base cation saturation ratio.

| **Fungi** | **C total** | **N total** | **S total** | **pH** | **P** | **K** | **Ca** | **Mg** | **Mn** | **Fe** | **Na** | **CEC** | **BCSR** |
|---|---|---|---|---|---|---|---|---|---|---|---|---|---|
| *Acidea* | -0.13 | -0.08 | 0.73 | -0.71 | 0.02 | -0.40 | -0.08 | -0.47 | -0.67 | 0.70 | -0.18 | 0.03 | -0.68 |
| *Alternaria* | 0.60 | 0.55 | -0.12 | 0.25 | 0.50 | 0.53 | 0.54 | 0.54 | 0.47 | -0.42 | 0.57 | 0.54 | 0.48 |
| *Articulospora* | 0.62 | 0.57 | -0.10 | 0.21 | 0.51 | 0.55 | 0.56 | 0.57 | 0.51 | -0.41 | 0.57 | 0.57 | 0.48 |
| *Cadophora* | -0.15 | -0.23 | -0.33 | 0.13 | -0.28 | -0.07 | -0.28 | -0.05 | 0.04 | -0.03 | -0.13 | -0.30 | 0.04 |
| *Cephalothecaceae_g* | 0.16 | 0.19 | 0.10 | 0.06 | 0.13 | 0.10 | 0.18 | 0.14 | 0.00 | -0.13 | 0.14 | 0.17 | 0.09 |
| *Chrysosporium* | 0.35 | 0.38 | -0.28 | 0.42 | 0.31 | 0.40 | 0.37 | 0.49 | 0.45 | -0.50 | 0.35 | 0.30 | 0.49 |
| *Ciliophora* | 0.39 | 0.42 | -0.17 | 0.34 | 0.36 | 0.45 | 0.36 | 0.44 | 0.44 | -0.50 | 0.41 | 0.35 | 0.45 |
| *Cladosporium* | 0.49 | 0.42 | -0.15 | 0.25 | 0.40 | 0.47 | 0.42 | 0.49 | 0.42 | -0.37 | 0.49 | 0.40 | 0.45 |
| *Eurotiomycetes_o_f_g* | -0.28 | -0.26 | 0.20 | -0.10 | -0.29 | -0.44 | -0.31 | -0.30 | -0.48 | 0.25 | -0.34 | -0.38 | -0.28 |
| *Fusarium* | 0.23 | 0.19 | -0.03 | 0.08 | 0.14 | 0.26 | 0.24 | 0.35 | 0.19 | -0.09 | 0.30 | 0.24 | 0.22 |
| *Gibberella* | 0.57 | 0.50 | -0.15 | 0.28 | 0.47 | 0.56 | 0.54 | 0.58 | 0.54 | -0.40 | 0.58 | 0.52 | 0.52 |
| *Lecythophora* | 0.01 | 0.07 | 0.20 | -0.30 | 0.02 | -0.18 | -0.04 | -0.13 | -0.28 | 0.17 | 0.00 | 0.03 | -0.25 |
| *Leptosphaeria* | -0.48 | -0.56 | -0.58 | 0.41 | -0.60 | -0.15 | -0.46 | -0.04 | 0.18 | -0.08 | -0.36 | -0.60 | 0.17 |
| *Lindtneria* | 0.06 | 0.06 | 0.06 | -0.01 | 0.14 | -0.08 | 0.01 | -0.05 | -0.03 | -0.04 | 0.00 | 0.03 | -0.08 |
| *Meliniomyces* | 0.09 | 0.10 | 0.52 | -0.46 | 0.16 | -0.15 | 0.04 | -0.28 | -0.40 | 0.33 | -0.06 | 0.13 | -0.38 |
| *Mortierella* | 0.60 | 0.55 | -0.07 | 0.19 | 0.51 | 0.53 | 0.55 | 0.52 | 0.41 | -0.37 | 0.57 | 0.56 | 0.41 |
| *Pezoloma* | -0.23 | -0.18 | 0.58 | -0.71 | -0.12 | -0.48 | -0.23 | -0.56 | -0.72 | 0.68 | -0.29 | -0.10 | -0.75 |
| *Phaeosphaeriaceae_g* | 0.62 | 0.59 | -0.12 | 0.26 | 0.54 | 0.55 | 0.55 | 0.55 | 0.50 | -0.46 | 0.58 | 0.55 | 0.48 |
| *Pleosporale_f_g* | 0.54 | 0.48 | -0.17 | 0.34 | 0.43 | 0.51 | 0.50 | 0.55 | 0.47 | -0.46 | 0.50 | 0.46 | 0.53 |
| *Pleosporales_fam_Incertae_sedis_g* | 0.63 | 0.58 | -0.10 | 0.23 | 0.54 | 0.59 | 0.59 | 0.60 | 0.52 | -0.42 | 0.63 | 0.59 | 0.51 |
| *Pyrenopeziza* | -0.09 | -0.05 | 0.74 | -0.71 | 0.05 | -0.41 | -0.04 | -0.45 | -0.72 | 0.68 | -0.18 | 0.07 | -0.67 |
| *Pyrenophora* | 0.62 | 0.58 | -0.06 | 0.18 | 0.52 | 0.54 | 0.56 | 0.53 | 0.46 | -0.39 | 0.57 | 0.57 | 0.43 |
| *Russula* | -0.07 | -0.02 | 0.53 | -0.52 | 0.08 | -0.20 | -0.05 | -0.34 | -0.46 | 0.41 | -0.09 | 0.07 | -0.50 |
| *Sebacinales_f_g* | -0.39 | -0.45 | -0.35 | 0.17 | -0.54 | -0.19 | -0.48 | -0.14 | 0.00 | -0.02 | -0.31 | -0.56 | 0.00 |
| *Sordariales_f_g* | -0.59 | -0.62 | -0.56 | 0.42 | -0.60 | -0.18 | -0.49 | -0.12 | 0.10 | -0.05 | -0.41 | -0.60 | 0.06 |
| *Sphaerosporella* | 0.35 | 0.27 | -0.44 | 0.35 | 0.15 | 0.33 | 0.22 | 0.37 | 0.35 | -0.39 | 0.31 | 0.13 | 0.44 |
| *Tomentella* | 0.09 | 0.13 | 0.44 | -0.33 | 0.25 | -0.07 | 0.19 | -0.10 | -0.17 | 0.19 | 0.07 | 0.27 | -0.18 |
| *Trichoderma* | -0.24 | -0.21 | 0.32 | -0.37 | -0.16 | -0.46 | -0.20 | -0.36 | -0.48 | 0.42 | -0.25 | -0.17 | -0.45 |
| *Vibrisseaceae_g* | 0.35 | 0.35 | 0.61 | -0.61 | 0.37 | 0.05 | 0.27 | -0.09 | -0.24 | 0.25 | 0.20 | 0.43 | -0.29 |
