## Supplementary Table 10 for "Plant Genotype Influences Physicochemical Properties of Substrate as well as Bacterial and Fungal Assemblages in the Rhizosphere of Balsam Poplar"

**Supplementary Table 10.** Spearman linear correlation analyses between bacterial and fungal taxa relative abundance in rhizosphere and tree growth measurements. Weak correlations (>|0.3|) are highlighted in red; moderate correlations (>|0.5|) are highlighted in yellow; strong correlations (>|0.7|) are highlighted in green.

| | | Chlorophyll content | | | | |
|---|---|---|---|---|---|---|
| **Bacteria** | **Growth** | **Season 1** | **Season 2** | **Shoot diameter** | **Plant biomass** | **Blooming** |
| *Anaerolineae SBR1031 A4b_g* | 0.068 | -0.143 | -0.184 | -0.117 | -0.134 | 0.036 |
| *Acidobacteriaceae_g* | -0.197 | 0.003 | 0.217 | 0.006 | 0.062 | -0.033 |
| *Bradyrhizobium* | -0.096 | -0.110 | 0.157 | -0.161 | -0.101 | 0.046 |
| *Caulobacteraceae_g* | -0.002 | 0.231 | 0.246 | 0.001 | 0.131 | 0.026 |
| *Chitinophagaceae_g* | 0.094 | 0.066 | -0.168 | 0.168 | 0.250 | -0.121 |
| *Acidimicrobiales EB1017_g* | 0.051 | -0.097 | -0.216 | 0.057 | 0.038 | -0.065 |
| *Alphaproteobacteria Ellin329_f_g* | -0.048 | 0.163 | 0.249 | 0.080 | 0.227 | 0.015 |
| *Betaproteobacteria Ellin6067_f_g* | -0.091 | -0.035 | -0.075 | -0.124 | 0.122 | -0.029 |
| *Frankiaceae_g* | -0.013 | 0.054 | -0.156 | 0.068 | 0.239 | -0.244 |
| *Gaiellaceae_g* | -0.099 | 0.067 | 0.047 | -0.023 | 0.072 | -0.049 |
| *Gemmataceae_g* | 0.071 | -0.076 | -0.210 | -0.131 | -0.320 | -0.006 |
| *Geobacter* | -0.020 | -0.053 | 0.164 | -0.001 | -0.117 | 0.276 |
| *Acidobacteria-6 iii1-15_f_g* | 0.173 | -0.100 | -0.319 | -0.091 | -0.251 | -0.067 |
| *Isosphaeraceae_g* | -0.148 | 0.043 | 0.173 | -0.028 | -0.013 | 0.016 |
| *Myxococcales_f_g* | -0.025 | -0.097 | -0.293 | -0.031 | -0.008 | -0.178 |
| *Opitutaceae_g* | 0.143 | -0.055 | -0.157 | -0.100 | -0.357 | 0.054 |
| *Opitutus* | 0.060 | 0.060 | -0.126 | 0.016 | -0.231 | 0.157 |
| *Pedosphaerales_f_g* | -0.074 | -0.051 | 0.329 | 0.012 | -0.042 | 0.220 |
| *Pirellulaceae_g* | 0.122 | 0.164 | -0.288 | 0.080 | 0.020 | -0.115 |
| *Planctomyces* | 0.133 | -0.072 | 0.030 | -0.045 | -0.130 | -0.045 |
| *Rhizobiales_f_g* | -0.030 | 0.028 | -0.248 | 0.377 | 0.035 | -0.148 |
| *Rhodoplanes* | -0.067 | 0.007 | 0.012 | -0.222 | -0.110 | 0.016 |
| *Rhodospirillaceae_g* | 0.064 | 0.091 | -0.055 | -0.013 | 0.197 | -0.065 |
| *Rubrivivax* | 0.064 | -0.066 | -0.119 | -0.081 | -0.166 | 0.189 |
| *Sinobacteraceae_g* | -0.146 | 0.035 | 0.150 | -0.005 | 0.263 | -0.201 |
| *Solibacterales_f_g* | -0.041 | -0.208 | 0.010 | -0.017 | -0.142 | 0.045 |
| *Solirubrobacterales_f_g* | -0.156 | -0.023 | 0.050 | 0.073 | 0.274 | -0.114 |
| *Sphingobacteriaceae_g* | -0.050 | 0.183 | 0.150 | 0.085 | 0.119 | 0.002 |
| *Phycisphaerae WD2101_f_g* | -0.089 | -0.046 | -0.003 | 0.073 | 0.207 | -0.136 |
| *Xanthomonadaceae_g* | -0.131 | 0.035 | 0.247 | -0.053 | 0.043 | 0.000 |

**Supplementary Table 10.** Spearman linear correlation analyses between bacterial and fungal taxa relative abundance in rhizosphere and tree growth measurements. Weak correlations (>|0.3|) are highlighted in red; moderate correlations (>|0.5|) are highlighted in yellow; strong correlations (>|0.7|) are highlighted in green.

| | | Chlorophyll content | | | | |
|---|---|---|---|---|---|---|
| **Fungi** | **Growth** | **Season 1** | **Season 2** | **Shoot diameter** | **Plant biomass** | **Blooming** |
| *Acidea* | 0.152 | 0.152 | 0.106 | -0.108 | -0.006 | 0.152 |
| *Alternaria* | 0.025 | 0.025 | 0.090 | 0.040 | 0.246 | 0.025 |
| *Articulospora* | 0.021 | 0.021 | 0.054 | 0.090 | 0.232 | 0.021 |
| *Cadophora* | 0.177 | 0.177 | 0.164 | 0.164 | -0.004 | 0.177 |
| *Cephalothecaceae_g* | 0.039 | 0.039 | 0.126 | -0.022 | 0.103 | 0.039 |
| *Chrysosporium* | 0.046 | 0.046 | 0.063 | 0.032 | -0.026 | 0.046 |
| *Ciliophora* | -0.032 | -0.032 | 0.052 | 0.201 | 0.112 | -0.032 |
| *Cladosporium* | 0.033 | 0.033 | 0.053 | 0.052 | 0.224 | 0.033 |
| *Eurotiomycetes_o_f_g* | 0.267 | 0.267 | 0.259 | -0.113 | -0.046 | 0.267 |
| *Fusarium* | 0.129 | 0.129 | 0.104 | 0.046 | 0.144 | 0.129 |
| *Gibberella* | 0.113 | 0.113 | 0.131 | -0.007 | 0.266 | 0.113 |
| *Lecythophora* | 0.047 | 0.047 | 0.111 | 0.241 | 0.057 | 0.047 |
| *Leptosphaeria* | 0.012 | 0.012 | -0.077 | -0.022 | -0.211 | 0.012 |
| *Lindtneria* | -0.073 | -0.073 | 0.097 | 0.126 | -0.001 | -0.073 |
| *Meliniomyces* | -0.024 | -0.024 | 0.008 | -0.049 | -0.018 | -0.024 |
| *Mortierella* | 0.043 | 0.043 | 0.101 | -0.040 | 0.310 | 0.043 |
| *Pezoloma* | 0.076 | 0.076 | 0.056 | -0.061 | 0.022 | 0.076 |
| *Phaeosphaeriaceae_g* | 0.033 | 0.033 | 0.095 | 0.059 | 0.282 | 0.033 |
| *Pleosporale_f_g* | 0.022 | 0.022 | 0.087 | 0.142 | 0.141 | 0.022 |
| *Pleosporales_fam_Incertae_sedis_g* | 0.038 | 0.038 | 0.105 | 0.070 | 0.311 | 0.038 |
| *Pyrenopeziza* | 0.109 | 0.109 | 0.102 | -0.191 | -0.003 | 0.109 |
| *Pyrenophora* | 0.069 | 0.069 | 0.120 | 0.049 | 0.301 | 0.069 |
| *Russula* | 0.030 | 0.030 | 0.043 | -0.099 | -0.013 | 0.030 |
| *Sebacinales_f_g* | 0.024 | 0.024 | 0.025 | 0.154 | -0.096 | 0.024 |
| *Sordariales_f_g* | -0.132 | -0.132 | -0.142 | 0.006 | -0.246 | -0.132 |
| *Sphaerosporella* | 0.062 | 0.062 | 0.018 | 0.069 | 0.044 | 0.062 |
| *Tomentella* | -0.190 | -0.190 | -0.165 | -0.082 | 0.104 | -0.190 |
| *Trichoderma* | 0.117 | 0.117 | 0.114 | -0.071 | 0.056 | 0.117 |
| *Vibrisseaceae_g* | 0.042 | 0.042 | 0.034 | 0.105 | 0.098 | 0.042 |
