## Supplementary Table 11 for "Plant Genotype Influences Physicochemical Properties of Substrate as well as Bacterial and Fungal Assemblages in the Rhizosphere of Balsam Poplar"

**Supplementary Table 11.** Pairwise comparison between origin of cuttings in each substrate type in the greenhouse experiment for bacterial and fungal taxa relative abundance. Two-way ANOVAs were used to discern how substrate type (Control, Tailings or Waste rock), origin of cuttings (La Corne Mine, Westwood or Natural forest) and their interaction influenced taxa relative abundance. Additional analyses were performed for each substrate type separately and between substrate types to better assess the indivudial effects of origin and substrate type.

| **Bacteria** | | *Anaerolineae SBR1031 A4b_g* | *Acidobacteriaceae_g* | *Bradyrhizobium* | *Caulobacteraceae_g* | *Chitinophagaceae_g* | *Acidimicrobiales EB1017_g* |
|---|---|---|---|---|---|---|---|
| | Origin | 0.511 | 0.300 | 0.120 | 0.062 | 0.030 | 0.252 |
| | Substrate type | < 0.001 | < 0.001 | < 0.001 | < 0.001 | < 0.001 | < 0.001 |
| | Interaction | 0.562 | 0.732 | 0.259 | 0.461 | 0.380 | 0.003 |
| **Pairwise comparison between origin** | | | | | | | |
| | Natural | 0.9% B | 0.8% A | 0.9% A | 1.1% A | 13.0% A | 0.8% A |
| | La Corne Mine | 1.0% A | 0.7% A | 0.9% A | 1.0% A | 12.8% A | 0.7% A |
| | Westwood | 0.9% B | 0.7% A | 0.9% A | 0.9% A | 15.1% A | 0.8% A |
| | p-value | 0.688 | 0.873 | 0.995 | 0.742 | 0.323 | 0.574 |
| **Pairwise comparison by substrate type** | | | | | | | |
| Control | Natural | 0.9% A | 0.1% A | 0.4% AB | 0.5% A | 23.7% AB | 1.0% A |
| | La Corne Mine | 0.8% A | 0.1% A | 0.5% A | 0.6% A | 19.8% B | 0.9% A |
| | Westwood | 0.7% A | 0.1% A | 0.4% B | 0.5% A | 24.1% A | 0.7% A |
| | p-value | 0.331 | 0.576 | 0.030 | 0.179 | 0.031 | 0.064 |
| Tailings | Natural | 1.4% A | 0.1% A | 0.8% A | 0.4% A | 11.2% A | 1.2% AB |
| | La Corne Mine | 1.3% A | 0.0% A | 1.0% A | 0.6% A | 11.3% A | 0.9% B |
| | Westwood | 1.4% A | 0.0% A | 0.9% A | 0.5% A | 12.7% A | 1.3% A |
| | p-value | 0.750 | 0.199 | 0.743 | 0.112 | 0.465 | 0.014 |
| Waste rock | Natural | 0.6% A | 2.0% A | 1.3% A | 2.0% A | 6.6% A | 0.3% A |
| | La Corne Mine | 0.8% A | 1.9% A | 1.1% A | 1.7% A | 7.8% A | 0.3% A |
| | Westwood | 0.6% A | 1.8% A | 1.3% A | 1.7% A | 9.3% A | 0.4% A |
| | p-value | 0.465 | 0.954 | 0.377 | 0.509 | 0.250 | 0.111 |

**Supplementary Table 11.** Pairwise comparison between origin of cuttings in each substrate type in the greenhouse experiment for bacterial and fungal taxa relative abundance. Two-way ANOVAs were used to discern how substrate type (Control, Tailings or Waste rock), origin of cuttings (La Corne Mine, Westwood or Natural forest) and their interaction influenced taxa relative abundance. Additional analyses were performed for each substrate type separately and between substrate types to better assess the indivudial effects of origin and substrate type.

| **Bacteria** | | *Alphaproteobacteria Ellin329_f_g* | *Betaproteobacteria Ellin6067_f_g* | *Frankiaceae_g* | *Gaiellaceae_g* | *Gemmataceae_g* | *Geobacter* | *Acidobacteria-6 iii1-15_f_g* |
|---|---|---|---|---|---|---|---|---|
| | Origin | 0.812 | 0.604 | 0.129 | 0.134 | 0.094 | 0.025 | 0.391 |
| | Substrate type | < 0.001 | < 0.001 | < 0.001 | < 0.001 | < 0.001 | < 0.001 | < 0.001 |
| | Interaction | 0.584 | 0.377 | 0.153 | 0.055 | 0.433 | 0.567 | 0.306 |
| **Pairwise comparison between origin** | | | | | | | | |
| | Natural | 6.5% A | 2.0% A | 1.2% A | 0.9% A | 2.1% A | 0.9% A | 1.3% A |
| | La Corne Mine | 5.9% A | 1.9% A | 0.9% A | 0.9% A | 1.8% A | 2.9% A | 1.2% A |
| | Westwood | 6.3% A | 1.9% A | 1.1% A | 0.9% A | 2.1% A | 1.6% A | 1.3% A |
| | p-value | 0.744 | 0.846 | 0.149 | 0.946 | 0.686 | 0.154 | 0.852 |
| **Pairwise comparison by substrate type** | | | | | | | | |
| Control | Natural | 6.6% A | 1.8% A | 1.4% A | 0.8% A | 0.9% A | 0.1% A | 0.9% A |
| | La Corne Mine | 6.3% A | 2.1% A | 1.4% A | 0.9% A | 0.7% A | 0.4% A | 0.9% A |
| | Westwood | 5.8% A | 2.1% A | 1.4% A | 0.8% A | 0.9% A | 0.0% A | 1.0% A |
| | p-value | 0.655 | 0.453 | 0.977 | 0.653 | 0.472 | 0.248 | 0.436 |
| Tailings | Natural | 3.9% A | 1.9% A | 1.1% A | 0.5% AB | 4.1% A | 2.8% A | 2.5% A |
| | La Corne Mine | 3.6% A | 1.6% A | 0.6% B | 0.5% B | 3.2% A | 7.3% A | 1.9% A |
| | Westwood | 3.4% A | 1.5% A | 0.7% AB | 0.7% A | 4.0% A | 4.9% A | 2.4% A |
| | p-value | 0.722 | 0.185 | 0.027 | 0.003 | 0.033 | 0.191 | 0.127 |
| Waste rock | Natural | 8.3% A | 2.1% A | 1.1% A | 1.2% A | 1.6% A | 0.1% A | 0.7% A |
| | La Corne Mine | 8.2% A | 2.1% A | 0.8% A | 1.2% A | 1.4% A | 0.4% A | 0.7% A |
| | Westwood | 9.4% A | 2.0% A | 1.1% A | 1.1% A | 1.5% A | 0.2% A | 0.6% A |
| | p-value | 0.415 | 0.875 | 0.175 | 0.640 | 0.898 | 0.123 | 0.634 |

**Supplementary Table 11.** Pairwise comparison between origin of cuttings in each substrate type in the greenhouse experiment for bacterial and fungal taxa relative abundance. Two-way ANOVAs were used to discern how substrate type (Control, Tailings or Waste rock), origin of cuttings (La Corne Mine, Westwood or Natural forest) and their interaction influenced taxa relative abundance. Additional analyses were performed for each substrate type separately and between substrate types to better assess the indivudial effects of origin and substrate type.

| **Bacteria** | | *Isosphaeraceae_g* | *Myxococcales_f_g* | *Opitutaceae_g* | *Opitutus* | *Pedosphaerales_f_g* | *Pirellulaceae_g* | *Planctomyces* |
|---|---|---|---|---|---|---|---|---|
| | Origin | 0.624 | 0.024 | 0.750 | 0.085 | 0.014 | 1.000 | 0.357 |
| | Substrate type | < 0.001 | < 0.001 | < 0.001 | < 0.001 | < 0.001 | < 0.001 | 0.005 |
| | Interaction | 0.414 | 0.147 | 0.188 | 0.020 | < 0.001 | 0.047 | 0.545 |
| **Pairwise comparison between origin** | | | | | | | | |
| | Natural | 1.2% A | 2.5% A | 1.3% A | 1.0% A | 0.9% A | 1.0% A | 2.9% A |
| | La Corne Mine | 0.8% A | 2.1% A | 1.0% A | 0.8% A | 0.9% A | 1.0% A | 3.2% A |
| | Westwood | 0.8% A | 2.5% A | 1.3% A | 0.8% A | 1.0% A | 0.9% A | 2.7% A |
| | p-value | 0.325 | 0.365 | 0.656 | 0.138 | 0.573 | 0.951 | 0.451 |
| **Pairwise comparison by substrate type** | | | | | | | | |
| Control | Natural | 0.4% A | 3.1% A | 0.3% A | 0.9% A | 0.7% AB | 1.4% A | 2.6% A |
| | La Corne Mine | 0.4% A | 2.9% A | 0.4% A | 0.8% A | 1.0% A | 1.3% A | 2.7% A |
| | Westwood | 0.4% A | 3.7% A | 0.4% A | 0.7% A | 0.5% B | 0.9% A | 2.2% A |
| | p-value | 0.790 | 0.293 | 0.683 | 0.242 | < 0.001 | 0.030 | 0.290 |
| Tailings | Natural | 0.3% A | 3.2% A | 3.7% A | 1.8% A | 0.8% A | 1.2% A | 2.2% A |
| | La Corne Mine | 0.3% A | 2.0% B | 1.9% A | 1.0% B | 0.8% A | 1.1% A | 2.6% A |
| | Westwood | 0.4% A | 2.5% AB | 3.3% A | 1.1% B | 0.6% A | 1.3% A | 2.6% A |
| | p-value | 0.473 | 0.004 | 0.063 | 0.002 | 0.301 | 0.554 | 0.516 |
| Waste rock | Natural | 2.4% A | 1.7% A | 0.3% A | 0.5% A | 1.1% AB | 0.5% A | 3.6% A |
| | La Corne Mine | 1.7% A | 1.5% A | 0.4% A | 0.7% A | 0.9% B | 0.5% A | 4.2% A |
| | Westwood | 1.6% A | 1.4% A | 0.4% A | 0.6% A | 1.8% A | 0.6% A | 3.3% A |
| | p-value | 0.232 | 0.752 | 0.805 | 0.478 | 0.003 | 0.479 | 0.513 |

**Supplementary Table 11.** Pairwise comparison between origin of cuttings in each substrate type in the greenhouse experiment for bacterial and fungal taxa relative abundance. Two-way ANOVAs were used to discern how substrate type (Control, Tailings or Waste rock), origin of cuttings (La Corne Mine, Westwood or Natural forest) and their interaction influenced taxa relative abundance. Additional analyses were performed for each substrate type separately and between substrate types to better assess the indivudial effects of origin and substrate type.

| **Bacteria** | | *Rhizobiales_f_g* | *Rhodoplanes* | *Rhodospirillaceae_g* | *Rubrivivax* | *Sinobacteraceae_g* | *Solibacterales_f_g* |
|---|---|---|---|---|---|---|---|
| | Origin | 0.689 | 0.192 | 0.217 | 0.426 | 0.436 | 0.594 |
| | Substrate type | 0.014 | < 0.001 | < 0.001 | < 0.001 | < 0.001 | < 0.001 |
| | Interaction | 0.268 | 0.926 | 0.038 | 0.140 | 0.956 | 0.959 |
| **Pairwise comparison between origin** | | | | | | | |
| | Natural | 1.9% A | 4.4% A | 6.8% A | 0.7% A | 2.1% A | 1.5% A |
| | La Corne Mine | 1.9% A | 4.5% A | 7.6% A | 2.0% A | 1.7% A | 1.7% A |
| | Westwood | 2.0% A | 4.1% A | 6.9% A | 1.1% A | 1.8% A | 1.6% A |
| | p-value | 0.818 | 0.547 | 0.462 | 0.104 | 0.501 | 0.541 |
| **Pairwise comparison by substrate type** | | | | | | | |
| Control | Natural | 2.8% A | 3.0% A | 9.1% B | 0.1% A | 1.7% A | 1.3% A |
| | La Corne Mine | 2.2% A | 3.3% A | 11.6% A | 0.2% A | 1.5% A | 1.5% A |
| | Westwood | 2.2% A | 2.9% A | 9.5% B | 0.1% A | 1.6% A | 1.4% A |
| | p-value | 0.217 | 0.361 | 0.005 | 0.586 | 0.626 | 0.498 |
| Tailings | Natural | 1.7% A | 3.9% A | 5.3% A | 2.3% A | 1.0% A | 2.1% A |
| | La Corne Mine | 1.8% A | 4.0% A | 4.7% A | 5.2% A | 0.9% A | 2.2% A |
| | Westwood | 2.0% A | 3.8% A | 4.7% A | 3.0% A | 1.0% A | 2.1% A |
| | p-value | 0.484 | 0.892 | 0.489 | 0.037 | 0.676 | 0.937 |
| Waste rock | Natural | 1.5% A | 5.6% A | 6.3% A | 0.0% A | 3.1% A | 1.3% A |
| | La Corne Mine | 1.8% A | 6.2% A | 7.2% A | 0.2% A | 2.8% A | 1.4% A |
| | Westwood | 1.9% A | 5.3% A | 6.6% A | 0.3% A | 2.7% A | 1.4% A |
| | p-value | 0.489 | 0.382 | 0.607 | 0.622 | 0.741 | 0.926 |

**Supplementary Table 11.** Pairwise comparison between origin of cuttings in each substrate type in the greenhouse experiment for bacterial and fungal taxa relative abundance. Two-way ANOVAs were used to discern how substrate type (Control, Tailings or Waste rock), origin of cuttings (La Corne Mine, Westwood or Natural forest) and their interaction influenced taxa relative abundance. Additional analyses were performed for each substrate type separately and between substrate types to better assess the indivudial effects of origin and substrate type.

| **Bacteria** | | *Solirubrobacterales_f_g* | *Sphingobacteriaceae_g* | *Phycisphaerae WD2101_f_g* | *Xanthomonadaceae_g* |
|---|---|---|---|---|---|
| | Origin | 0.543 | 0.718 | 0.230 | 0.670 |
| | Substrate type | < 0.001 | < 0.001 | < 0.001 | < 0.001 |
| | Interaction | 0.185 | 0.677 | 0.139 | 0.498 |
| **Pairwise comparison between origin** | | | | | |
| | Natural | 0.9% A | 1.4% A | 1.5% A | 3.0% A |
| | La Corne Mine | 0.8% A | 1.1% A | 1.8% A | 2.2% A |
| | Westwood | 0.9% A | 1.0% A | 2.4% A | 2.3% A |
| | p-value | 0.628 | 0.409 | 0.066 | 0.705 |
| **Pairwise comparison by substrate type** | | | | | |
| Control | Natural | 1.2% A | 1.0% A | 2.5% A | 0.1% A |
| | La Corne Mine | 1.3% A | 0.9% A | 2.9% A | 0.1% A |
| | Westwood | 1.5% A | 0.8% A | 3.6% A | 0.1% A |
| | p-value | 0.176 | 0.854 | 0.103 | 0.339 |
| Tailings | Natural | 0.5% A | 0.2% A | 0.8% A | 0.0% A |
| | La Corne Mine | 0.5% A | 0.2% A | 1.0% A | 0.1% A |
| | Westwood | 0.6% A | 0.2% A | 0.9% A | 0.0% A |
| | p-value | 0.816 | 0.917 | 0.749 | 0.544 |
| Waste rock | Natural | 1.0% A | 2.5% A | 1.4% A | 7.2% A |
| | La Corne Mine | 0.8% A | 2.2% A | 1.7% A | 6.4% A |
| | Westwood | 0.8% A | 1.8% A | 2.5% A | 6.1% A |
| | p-value | 0.171 | 0.287 | 0.093 | 0.674 |

**Supplementary Table 11.** Pairwise comparison between origin of cuttings in each substrate type in the greenhouse experiment for bacterial and fungal taxa relative abundance. Two-way ANOVAs were used to discern how substrate type (Control, Tailings or Waste rock), origin of cuttings (La Corne Mine, Westwood or Natural forest) and their interaction influenced taxa relative abundance. Additional analyses were performed for each substrate type separately and between substrate types to better assess the indivudial effects of origin and substrate type.

| **Fungi** | | *Acidea* | *Alternaria* | *Articulospora* | *Cadophora* | *Cephalothecaceae_g* | *Chrysosporium* | *Ciliophora* | *Cladosporium* |
|---|---|---|---|---|---|---|---|---|---|
| | Origin | 0.672 | 0.397 | 0.562 | 0.541 | 0.349 | 0.950 | 0.123 | 0.121 |
| | Substrate type | < 0.001 | < 0.001 | < 0.001 | 0.108 | 0.094 | < 0.001 | < 0.001 | < 0.001 |
| | Interaction | 0.418 | 0.774 | 0.827 | 0.587 | 0.874 | 0.922 | 0.040 | 0.770 |
| **Pairwise comparison between origin** | | | | | | | | | |
| | Natural | 1.6% A | 1.2% A | 0.4% A | 0.1% A | 1.7% A | 14.3% A | 5.7% A | 0.4% A |
| | La Corne Mine | 1.0% A | 1.0% A | 0.4% A | 0.3% A | 1.4% A | 14.8% A | 5.9% A | 0.4% A |
| | Westwood | 1.1% A | 1.3% A | 0.5% A | 0.3% A | 1.5% A | 14.4% A | 8.6% A | 0.5% A |
| | p-value | 0.547 | 0.488 | 0.629 | 0.459 | 0.377 | 0.940 | 0.283 | 0.176 |
| **Pairwise comparison by substrate type** | | | | | | | | | |
| Control | Natural | 0,0% A | 2.4% A | 0.9% A | 0.1% A | 1.9% A | 18.9% A | 7.0% B | 0.8% A |
| | La Corne Mine | 0.0% A | 1.9% A | 0.8% A | 0.3% A | 1.7% A | 18.2% A | 14.5% AB | 0.6% A |
| | Westwood | 0.0% A | 2.2% A | 0.9% A | 0.1% A | 1.5% A | 17.7% A | 18.8% A | 0.8% A |
| | p-value | 0.807 | 0.432 | 0.497 | 0.424 | 0.677 | 0.846 | 0.031 | 0.519 |
| Tailings | Natural | 0.0% A | 0.7% A | 0.2% A | 0.2% A | 1.5% A | 13.2% A | 7.6% A | 0.3% A |
| | La Corne Mine | 0.0% A | 0.6% A | 0.2% A | 0.4% A | 1.0% A | 15.0% A | 2.6% A | 0.2% A |
| | Westwood | 0.0% A | 0.9% A | 0.3% A | 0.8% A | 1.3% A | 13.9% A | 4.0% A | 0.5% A |
| | p-value | 0.321 | 0.608 | 0.912 | 0.409 | 0.364 | 0.768 | 0.070 | 0.257 |
| Waste rock | Natural | 4.4% A | 0.5% A | 0.2% A | 0.1% A | 1.6% A | 11.3% A | 3.0% A | 0.2% A |
| | La Corne Mine | 3.0% A | 0.7% A | 0.3% A | 0.1% A | 1.4% A | 11.4% A | 1.7% A | 0.3% A |
| | Westwood | 3.1% A | 0.8% A | 0.3% A | 0.2% A | 1.5% A | 12.0% A | 3.5% A | 0.4% A |
| | p-value | 0.367 | 0.618 | 0.586 | 0.759 | 0.768 | 0.927 | 0.387 | 0.495 |

**Supplementary Table 11.** Pairwise comparison between origin of cuttings in each substrate type in the greenhouse experiment for bacterial and fungal taxa relative abundance. Two-way ANOVAs were used to discern how substrate type (Control, Tailings or Waste rock), origin of cuttings (La Corne Mine, Westwood or Natural forest) and their interaction influenced taxa relative abundance. Additional analyses were performed for each substrate type separately and between substrate types to better assess the indivudial effects of origin and substrate type.

| **Fungi** | | *Eurotiomycetes_o_f_g* | *Fusarium* | *Gibberella* | *Lecythophora* | *Leptosphaeria* | *Lindtneria* | *Meliniomyces* | *Mortierella* |
|---|---|---|---|---|---|---|---|---|---|
| | Origin | 0.032 | 0.194 | 0.533 | 0.453 | 0.564 | 0.527 | 0.761 | 0.692 |
| | Substrate type | < 0.001 | 0.692 | < 0.001 | 0.199 | 0.005 | 0.986 | 0.001 | < 0.001 |
| | Interaction | 0.516 | 0.759 | 0.817 | 0.408 | 0.293 | 0.485 | 0.008 | 0.578 |
| **Pairwise comparison between origin** | | | | | | | | | |
| | Natural | 0.5% A | 2.0% A | 0.5% A | 0.4% A | 0.7% A | 0.9% A | 2.5% A | 0.3% A |
| | La Corne Mine | 0.6% A | 2.1% A | 0.5% A | 0.4% A | 0.1% A | 0.5% A | 0.4% B | 0.3% A |
| | Westwood | 0.8% A | 2.5% A | 0.6% A | 0.3% A | 0.3% A | 0.7% A | 0.6% B | 0.3% A |
| | p-value | 0.195 | 0.720 | 0.728 | 0.615 | 0.350 | 0.516 | 0.010 | 0.751 |
| **Pairwise comparison by substrate type** | | | | | | | | | |
| Control | Natural | 0.3% A | 2.4% A | 0.9% A | 0.5% A | 0.0% A | 1.5% A | 0.0% A | 0.7% A |
| | La Corne Mine | 0.4% A | 1.8% A | 0.9% A | 0.3% A | 0.0% A | 0.6% A | 0.0% A | 0.5% A |
| | Westwood | 0.5% A | 2.2% A | 1.0% A | 0.3% A | 0.0% A | 0.4% A | 0.0% A | 0.6% A |
| | p-value | 0.313 | 0.195 | 0.789 | 0.401 | 0.390 | 0.307 | 0.841 | 0.492 |
| Tailings | Natural | 0.5% B | 1.8% A | 0.4% A | 0.4% A | 2.1% A | 0.4% A | 0.0% A | 0.1% A |
| | La Corne Mine | 0.7% AB | 1.6% A | 0.4% A | 0.2% A | 0.4% A | 0.3% A | 0.0% A | 0.2% A |
| | Westwood | 0.9% A | 2.7% A | 0.3% A | 0.3% A | 0.9% A | 1.0% A | 0.2% A | 0.2% A |
| | p-value | 0.022 | 0.469 | 0.943 | 0.529 | 0.240 | 0.395 | 0.441 | 0.924 |
| Waste rock | Natural | 0.8% A | 1.9% A | 0.2% A | 0.4% A | 0,0% A | 0.8% A | 6.7% A | 0.1% A |
| | La Corne Mine | 0.9% A | 3.0% A | 0.3% A | 0.7% A | 0,0% A | 0.5% A | 1.0% B | 0.2% A |
| | Westwood | 0.8% A | 2.6% A | 0.4% A | 0.4% A | 0.0% A | 0.7% A | 1.5% B | 0.3% A |
| | p-value | 0.903 | 0.832 | 0.330 | 0.292 | 0.508 | 0.775 | 0.002 | 0.360 |

**Supplementary Table 11.** Pairwise comparison between origin of cuttings in each substrate type in the greenhouse experiment for bacterial and fungal taxa relative abundance. Two-way ANOVAs were used to discern how substrate type (Control, Tailings or Waste rock), origin of cuttings (La Corne Mine, Westwood or Natural forest) and their interaction influenced taxa relative abundance. Additional analyses were performed for each substrate type separately and between substrate types to better assess the indivudial effects of origin and substrate type.

| **Fungi** | | *Pezoloma* | *Phaeosphaeriaceae_g* | *Pleosporale_f_g* | *Pleosporales _fam_Incertae_sedis_g* | *Pyrenopeziza* | *Pyrenophora* | *Russula* |
|---|---|---|---|---|---|---|---|---|
| | Origin | 0.459 | 0.242 | 0.628 | 0.259 | 0.310 | 0.466 | 0.538 |
| | Substrate type | < 0.001 | < 0.001 | < 0.001 | < 0.001 | < 0.001 | < 0.001 | < 0.001 |
| | Interaction | 0.208 | 0.961 | 0.707 | 0.834 | 0.405 | 0.756 | 0.806 |
| **Pairwise comparison between origin** | | | | | | | | |
| | Natural | 2.7% A | 0.4% A | 0.3% A | 1.6% A | 0.5% A | 0.4% A | 2.1% A |
| | La Corne Mine | 2.4% A | 0.3% A | 0.3% A | 1.5% A | 0.3% A | 0.4% A | 1.9% A |
| | Westwood | 2.1% A | 0.4% A | 0.3% A | 1.9% A | 0.3% A | 0.5% A | 2.4% A |
| | p-value | 0.856 | 0.372 | 0.653 | 0.490 | 0.533 | 0.643 | 0.910 |
| **Pairwise comparison by substrate type** | | | | | | | | |
| Control | Natural | 0.0% A | 0.7% A | 0.6% A | 3.3% A | 0,0% A | 1.0% A | 0.0% A |
| | La Corne Mine | 0.0% A | 0.7% A | 0.4% A | 2.8% A | 0,0% A | 0.8% A | 0.0% A |
| | Westwood | 0.0% A | 0.8% A | 0.5% A | 3.3% A | 0,0% A | 0.9% A | 0.0% A |
| | p-value | 0.778 | 0.526 | 0.511 | 0.525 | 0.999 | 0.506 | 0.551 |
| Tailings | Natural | 0.0% A | 0.2% A | 0.2% A | 0.8% A | 0,0% A | 0.1% A | 0.0% A |
| | La Corne Mine | 0.0% A | 0.2% A | 0.2% A | 0.9% A | 0,0% A | 0.2% A | 0.0% A |
| | Westwood | 0.0% A | 0.3% A | 0.2% A | 1.0% A | 0.0% A | 0.3% A | 0.3% A |
| | p-value | 0.046 | 0.442 | 0.473 | 0.873 | 0.479 | 0.650 | 0.464 |
| Waste rock | Natural | 7.1% A | 0.2% A | 0.2% A | 0.7% A | 1.2% A | 0.2% A | 5.5% A |
| | La Corne Mine | 7.2% A | 0.2% A | 0.2% A | 0.9% A | 0.9% A | 0.3% A | 5.6% A |
| | Westwood | 5.8% A | 0.2% A | 0.2% A | 1.3% A | 0.8% A | 0.3% A | 6.5% A |
| | p-value | 0.670 | 0.764 | 0.882 | 0.273 | 0.329 | 0.551 | 0.937 |

**Supplementary Table 11.** Pairwise comparison between origin of cuttings in each substrate type in the greenhouse experiment for bacterial and fungal taxa relative abundance. Two-way ANOVAs were used to discern how substrate type (Control, Tailings or Waste rock), origin of cuttings (La Corne Mine, Westwood or Natural forest) and their interaction influenced taxa relative abundance. Additional analyses were performed for each substrate type separately and between substrate types to better assess the indivudial effects of origin and substrate type.

| **Fungi** | | *Sebacinales_f_g* | *Sordariales_f_g* | *Sphaerosporella* | *Tomentella* | *Trichoderma* | *Vibrisseaceae_g* |
|---|---|---|---|---|---|---|---|
| | Origin | 0.872 | 0.594 | 0.021 | 0.518 | 0.820 | 0.272 |
| | Substrate type | < 0.001 | < 0.001 | < 0.001 | < 0.001 | 0.009 | < 0.001 |
| | Interaction | 0.476 | 0.821 | 0.338 | 0.858 | 0.517 | 0.604 |
| **Pairwise comparison between origin** | | | | | | | |
| | Natural | 0.5% A | 10.8% A | 3.8% B | 34.1% A | 1.7% A | 0.2% A |
| | La Corne Mine | 0.9% A | 12.3% A | 9.6% A | 28.5% A | 0.9% A | 0.3% A |
| | Westwood | 0.9% A | 10.6% A | 5.0% B | 31.7% A | 0.8% A | 0.3% A |
| | p-value | 0.517 | 0.897 | 0.007 | 0.516 | 0.197 | 0.981 |
| **Pairwise comparison by substrate type** | | | | | | | |
| Control | Natural | 0.3% A | 0.1% A | 6.5% A | 38.8% A | 0.4% A | 0.3% A |
| | La Corne Mine | 0.3% A | 0.1% A | 13.8% A | 30.1% A | 0.4% A | 0.2% A |
| | Westwood | 0.2% A | 0.1% A | 6.9% A | 30.0% A | 0.3% A | 0.1% A |
| | p-value | 0.707 | 0.588 | 0.044 | 0.428 | 0.924 | 0.256 |
| Tailings | Natural | 0.8% A | 34.1% A | 4.4% A | 21.6% A | 0.8% A | 0.0% A |
| | La Corne Mine | 2.1% A | 32.9% A | 11.4% A | 18.7% A | 0.6% A | 0.0% A |
| | Westwood | 1.8% A | 32.5% A | 5.2% A | 21.0% A | 0.7% A | 0.0% A |
| | p-value | 0.424 | 0.965 | 0.937 | 0.921 | 0.920 | 0.350 |
| Waste rock | Natural | 0.3% A | 0.2% A | 1.0% A | 40.5% A | 3.5% A | 0.4% A |
| | La Corne Mine | 0.2% A | 1.3% A | 3.9% A | 37.6% A | 1.5% A | 0.6% A |
| | Westwood | 0.7% A | 0.0% A | 3.2% A | 43.2% A | 1.3% A | 0.6% A |
| | p-value | 0.502 | 0.486 | 0.345 | 0.679 | 0.190 | 0.777 |
